# XAI-based Data Visualization in Multimodal Medical Data

**DOI:** 10.1101/2025.07.11.664302

**Authors:** Sahil Sharma, Muskaan Singh, Liam McDaid, Saugat Bhattacharyya

## Abstract

Explainable Artificial Intelligence (XAI) is crucial in healthcare as it helps make intricate machine learning models understandable and clear, especially when working with diverse medical data, enhancing trust, improving diagnostic accuracy, and facilitating better patient outcomes. This paper thoroughly examines the most advanced XAI techniques used in multimodal medical datasets. These strategies include perturbation-based methods, concept-based explanations, and example-based explanations. The value of perturbation-based approaches such as LIME and SHAP in explaining model predictions in medical diagnostics is explored. The paper discusses using concept-based explanations to connect machine learning results with concepts humans can understand. This helps to improve the interpretability of models that handle different types of data, including electronic health records (EHRs), behavioural, omics, sensors, and imaging data. Example-based strategies, such as prototypes and counterfactual explanations, are emphasised for offering intuitive and accessible explanations for healthcare judgments. The paper also explores the difficulties encountered in this field, which include managing data with high dimensions, balancing the tradeoff between accuracy and interpretability, and dealing with limited data by generating synthetic data. Recommendations in future studies focus on improving the practicality and dependability of XAI in clinical settings.

## 1 Introduction

The use of Artificial Intelligence (AI) in healthcare has had a significant impact on medical diagnosis, prognostics, and therapy planning. It transforms patient care by facilitating early disease detection, creating customised treatment strategies, optimising resource allocation, and reducing healthcare costs. The complex and diverse nature of medical data, including imaging (such as MRI and CT scans), genetics, electronic health records (EHRs), and sensor data, offers significant opportunities for AI models to extract valuable information [1]. While AI models can potentially improve clinical decision-making, they often function as opaque systems, hiding the reasoning behind their predictions [2]. Lack of transparency poses a significant obstacle to the acceptance and dependence of AI systems in medicine, where decisions can have substantial consequences for patient care [3].

In 2016, the Defense Advanced Research Projects Agency (DARPA) launched the Explainable Artificial Intelligence (XAI) program in response to these challenges^1^. As a result, XAI has focused on using visualisation tools to simplify the decision-making processes of complex AI models that manage multimodal medical data. Visualisation techniques are crucial in the healthcare sector for Explainable Artificial Intelligence (XAI) since they enable healthcare practitioners to interpret and clarify the decision-making process of AI models, providing a more profound understanding of their underlying mechanisms.

Visualisation methods are crucial since they enable medical practitioners to comprehend and evaluate AI’s judgements. These methods help to navigate the complexities of multimodal medical data which spans several forms including imaging (e.g., MRI, PET, and CT scans), genetic sequences, and structured datasets. Widely accepted for their ability to identify important features in models that examine medical images are methods including Layer-wise Relevance Propagation (LRP) [4], Gradient-weighted Class Activation Mapping (Grad-CAM) [5], and SHapley Additive exPlanations (SHAP) [6]. Conversely, in exploring the link between features and predictions in structured data models, methods like Partial Dependence Plots (PDP) [7] and Individual Conditional Expectation (ICE) plots [8] are very effective. Utilising the XAI-based visualisation techniques equips medical professionals to understand artificial intelligence decision-making and build trust in its findings. The transparency of AI recommendations is essential for accuracy and accountability, ensuring that the reasoning behind these ideas is precise and reliable [3]. By employing sophisticated visualisation tools, medical professionals can enhance their understanding of AI applications and improve the effectiveness of AI and patient care outcomes.

The study of artificial intelligence (AI) in healthcare involves analysing several types of data, such as imaging data, electronic health records, behavioural data, omics, and sensor data [9–13]. Many machine learning (especially ones based on deep learning) models in this field offer limited insight into their decision-making mechanisms [14] driving the increasing need for interpretable AI. Deep learning models are now making more efforts to clarify their decision-making process to healthcare professionals by utilising explainable AI (XAI) methodologies [15, 16]. Jin et al. [17] have contributed to understanding how deep learning is applied in clinical decision support systems. In addition, Joshi et al. [18] have thoroughly examined multimodal deep neural networks in several fields. The advancements above highlight the need for a methodical examination of XAI in medicine.

### Contribution

XAI has been surveyed and reviewed from multiple aspects of the medical domain [19, 20], such as medical image classification [21], visual analytics [22], interpretability in medical imaging [23], trustworthy AI in healthcare [1], IoMT [24], and evaluating XAI [25]. We relate this work to multimodal medical data [26, 27] and visualisation [28, 29]. This work presents the tools, frameworks, and challenges for data visualisation-based explainability in multimodal medical data. The review in this paper has been compared with existing surveys in Table 1.

**Table 1.**
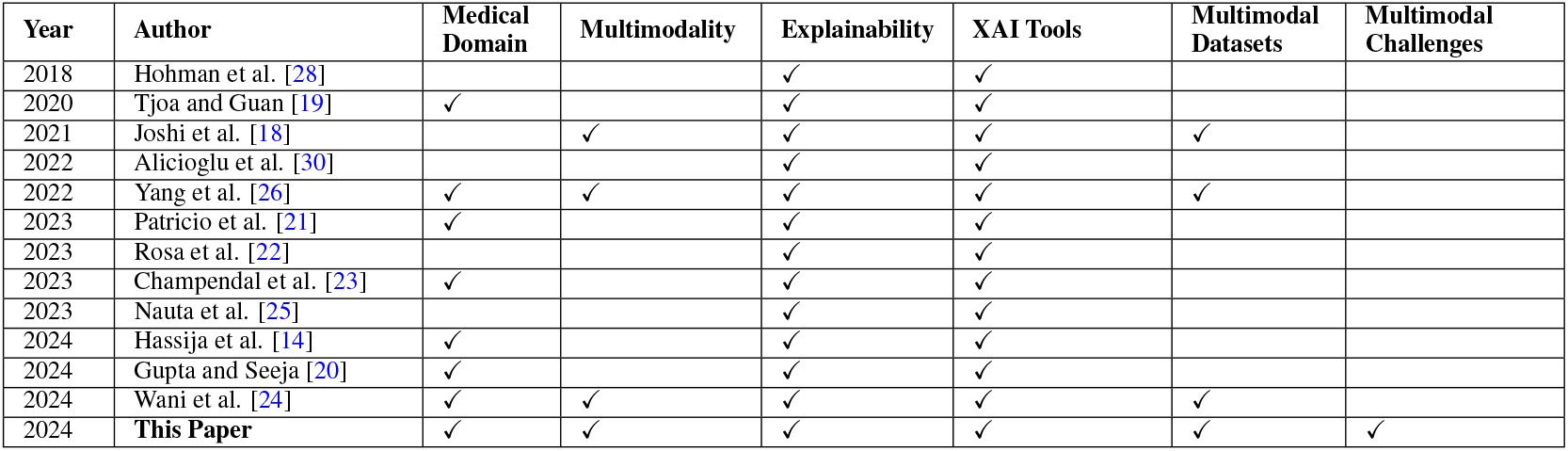
Comparison with Existing Surveys

| Year | Author | Medical Domain | Multimodality | Explainability | XAI Tools | Multimodal Datasets | Multimodal Challenges |
| --- | --- | --- | --- | --- | --- | --- | --- |
| 2018 | Hohman et al. [28] |  |  | ✓ | ✓ |  |  |
| 2020 | Tjoa and Guan [19] | ✓ |  | ✓ | ✓ |  |  |
| 2021 | Joshi et al. [18] |  | ✓ | ✓ | ✓ | ✓ |  |
| 2022 | Alicioglu et al. [30] |  |  | ✓ | ✓ |  |  |
| 2022 | Yang et al. [26] | ✓ | ✓ | ✓ | ✓ | ✓ |  |
| 2023 | Patricio et al. [21] | ✓ |  | ✓ | ✓ |  |  |
| 2023 | Rosa et al. [22] |  |  | ✓ | ✓ |  |  |
| 2023 | Champendal et al. [23] | ✓ |  | ✓ | ✓ |  |  |
| 2023 | Nauta et al. [25] |  |  | ✓ | ✓ |  |  |
| 2024 | Hassija et al. [14] | ✓ |  | ✓ | ✓ |  |  |
| 2024 | Gupta and Seeja [20] | ✓ |  | ✓ | ✓ |  |  |
| 2024 | Wani et al. [24] | ✓ | ✓ | ✓ | ✓ | ✓ |  |
| 2024 | **This Paper** | ✓ | ✓ | ✓ | ✓ | ✓ | ✓ |

This paper contributes to the existing literature in the following ways -

- A comprehensive review of recent literature on XAI-based visualisation over multimodal aspects of medical data.
- A detailed discussion on existing frameworks for XAI over multimodal medical data.
- An exhaustive list of open and restricted multimodal medical datasets used in the XAI literature.
- Discussion on current challenges in XAI data visualisation and future scope.

### Significance

In modern times, Responsible AI principles emphasise the necessity for transparency, accountability, and trust, especially in critical areas such as healthcare [31]. Our emphasis on XAI-based data visualisation corresponds to these ethical and clinical requirements. This multimodal medical data management guide builds on existing research to identify the best explainable approaches for specific case studies. This research integrates Explainable AI (XAI) approaches with multimodal medical data to improve AI models’ transparency, interpretability, and clinical usefulness. This study discusses crucial methods for enhancing clinical decision support systems.

These XAI methods will improve patient outcomes by enabling earlier and more accurate diagnoses. Integrating EHRs with omics and sensor data offers an integrated view on patient health, enabling doctors to recognise patterns that may otherwise remain undetected [32]. This study shows that XAI could overcome the knowledge gap between AI developers and doctors beyond clinical applications. This research discusses the AI models that help interpret medical decisions in multimodal settings. This enables collaboration and trust across different groups, leading to more robust and broadly accepted healthcare AI solutions. Standardised datasets and frameworks improve study uniformity and reproducibility, advancing the field. In addition to its technological contributions, this research could change healthcare practices by making AI-driven insights more accessible and useful for medical practitioners globally.

### A Problem statement

Applying Explainable AI (XAI) to multimodal medical data poses difficulties in guaranteeing the transparency and interpretability of AI models. Multimodal medical data refers to the information derived from various sources, such as electronic health records (EHRs), behavioural data, omics data, sensor data (including wearable devices and IoT systems), and medical imaging. Every data source offers distinct and significant information essential for precise diagnosis and treatment.

This study outlines a mathematical framework that addresses the challenges of integrating varying subsets of modalities and ensuring explainability in the predictions. Through this formulation, any combination of at least two modalities qualifies as multimodal data, allowing flexible integration and explanation of the model’s outputs. We define the set of all possible modalities in Equation 1.

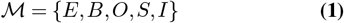

where *E* represents electronic health records (EHR), *B* denotes behavioural data, *O* denotes the Omics data, *S* as Sensor-based data, and *I* the Image-based data. Each patient/sample may have only a subset of these modalities available. We consider a dataset 𝒟 consisting of *N* samples.

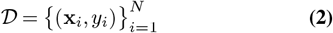

where **x**_*i*_ represents the multimodal input for the *i*-th sample, and *y*_*i*_ denotes the corresponding target label such as if the disease is present or not. For each sample *i*, the set of available modalities is denoted by ℳ_*i*_ ⊆ ℳ. If |ℳ_*i*_ | ≥ 2, the sample qualifies as multimodal. The input vector for sample *i* is given by Equation 3.

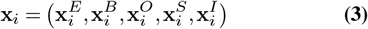

where:

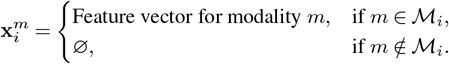

This construction allows flexibility in handling missing modalities while maintaining compatibility with the multi-modal framework.

We define a predictive model *f* that maps the multimodal input **x**_*i*_ to the target output *ŷ*_*i*_.

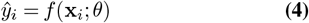

where *θ* denotes the set of trainable parameters. Depending on the architecture, the model can perform *early fusion* (concatenating raw features), *intermediate fusion* (merging modality-specific representations mid-network), or *late fusion* (aggregating predictions from individual modalities). To ensure interpretability, we introduce an explainability function *g*, which explains *e*_*i*_ for the prediction *ŷ*_*i*_.

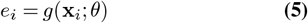

By converting the internal model reasoning into a form understandable to doctors, the explainability function *g* emphasises how various modalities or features affect the outcome. By methodically assessing the contributions of characteristics and modalities inside the predictive model, one ensures openness and relevance to clinical workflows.

The function *g* operates at feature level as well as modality level. *g* identifies the importance of specific features within a modality, providing insights into the key variables influencing the prediction. For instance, in the case of EHR data, it may highlight critical lab values or diagnoses. Also, *g* assesses the relative importance of entire modalities, offering a holistic view of which data types (e.g., imaging, omics, or behavioural data) had the most significant impact on the model’s decision. This is particularly valuable in multimodal settings, where clinicians need to understand the role of each data source.

The output *e*_*i*_ can take four forms depending on the modality and clinical requirements. The first type is visual explanations, such as saliency maps for image-based modalities to localize regions of diagnostic importance. The next is the quantitative scores, where feature importance or Shapley values for tabular data rank variables based on their contribution. The third type is temporal explanations, where we have attention scores or temporal heatmaps for time-series data, revealing critical time points or trends. The fourth is the counterfactual scenario, where we hypothetically illustrate how small input changes could alter predictions, aiding clinical decision-making.

*g* can include domain-specific knowledge to improve interpretability even more by matching explanations with accepted clinical guidelines and hence providing actionable insights. For omics data, *g* might link feature contributions to biological processes, offering fundamental knowledge of the predictions. All things considered, the explainability feature (g) not only decodes the inner workings of the predictive model but also closes the gap between sophisticated AI outputs and clinician-friendly interpretations, so guaranteeing transparent, practical, and in line with real-world medical decision-making predictions of the model.

This concept recognises the importance of interpretability in AI-powered medical diagnostics, ensuring that the predictions made by the model are precise and can be efficiently conveyed to clinicians to facilitate well-informed decisionmaking. Figure 1 represents the basic idea of what explainability can offer in the multimodal medical setting.

**Fig. 1.**
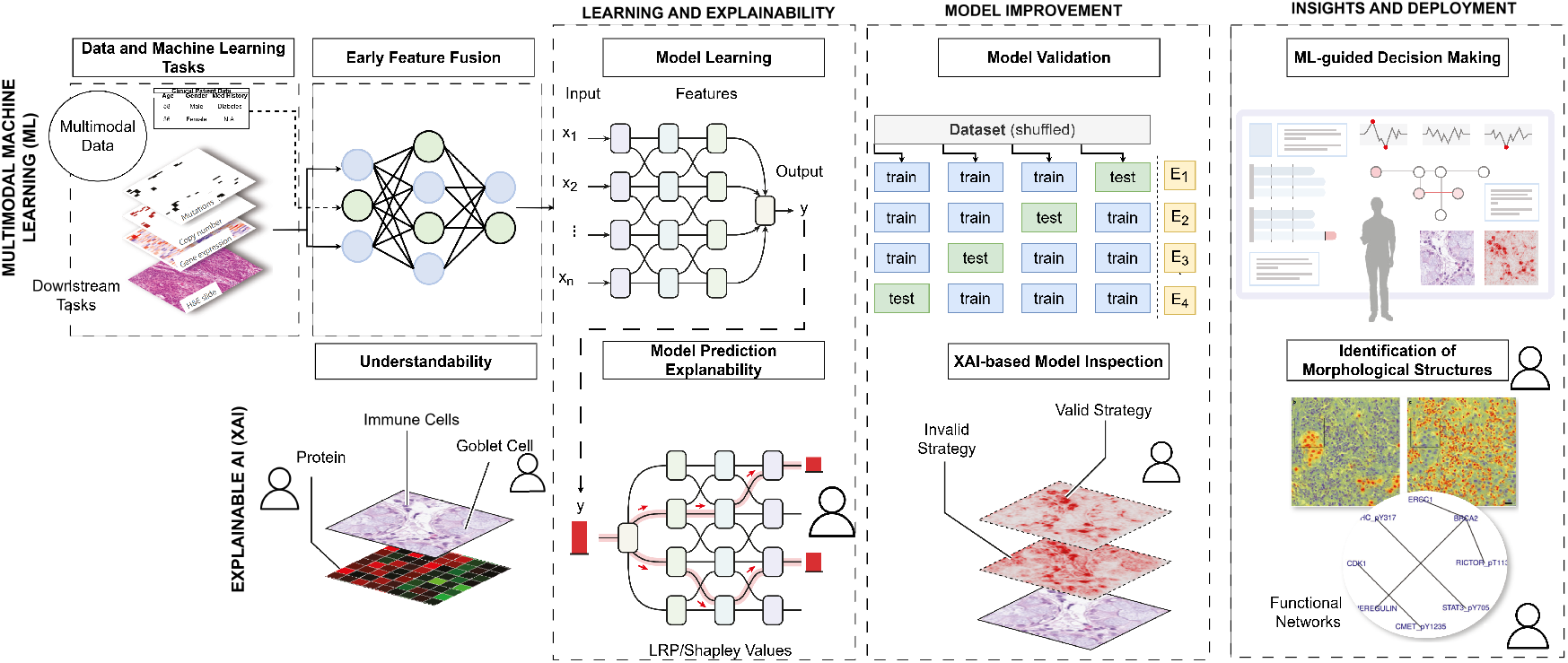
Overview of multimodal ML and XAI pipeline. Adapted from [33]. This image is licensed under CC BY 4.0.

### B Review method

This study reviews and analyses the groundbreaking research in XAI-based data visualisation in multimodal medical data. To conduct this systematic review, we first lay down the bibliography search strategy in which specific search engines were used to find the relevant publications listed. It is essential to filter out the publications using specific keywords based on the date of the bibliographic search. Once the papers were collected from the search engines, the most important studies were chosen according to the criteria for inclusion and exclusion.

To establish a search strategy that is both replicable and clear, it was essential to define the following elements: (1) The specific search engines that will be utilised to conduct the scientific database search; (2) the limitations imposed to narrow down the results; and (3) the criteria for including or excluding publications to focus solely on the most important ones. ***Search engines*** The search engines used in the current review are Engineering Village^2^, Scopus^3^, Web of Science^4^, PubMed^5^, and Google Scholar^6^. These research engines offer a comprehensive coverage of scientific publications. Furthermore, we have included additional publications in the reference lists of the main articles reviewed as worthy candidates for this review.

#### Search limits

A preliminary analysis in state of the art in the field of XAI in multimodal medical data enabled the identification of the relevant search terms and keywords covering multi-aspects of the topic, such as interpretability, attention visualisation, feature attribution, saliency methods, model agnostic and model specific, and sensitivity of the medical data involved.

The identified fundamental keywords have been further developed and expanded to include a broader range of topics in multimodal medical data and explainable artificial intelligence (XAI). The search terms were systematically categorised for precise searches across several document fields, including the title, abstract, keywords, and topic fields. The search method was established using logical operators and specific phrases about Explainable Artificial Intelligence (XAI) in multimodal medical contexts. The primary focus of the search was on strategies related to explainability. The search query was constructed as follows: (XAI OR “explainable AI” OR “explainable artificial intelligence”) AND “multimodal” AND (medic* OR healthcare) AND (“attention visualization” OR “feature attribution” OR “saliency methods” OR “model agnostic” OR “model specific” OR “in-model” OR “post-hoc”). Figure 2 shows the count of publications over the years 2016 to 2024 using the above query.

**Fig. 2.**
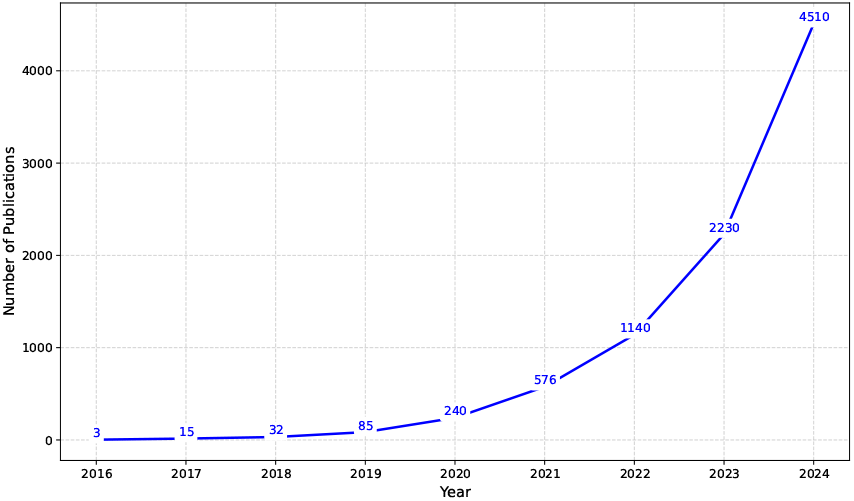
Overview of XAI publications retrieved from Google Scholar.

We limited our search to publications starting from 1st January 2016 to 31st December 2024 (date of the bibliographic search). The search was restricted to peer-reviewed conferences and journal articles written exclusively in English. These papers offer a comprehensive perspective on established and verified methodology and expertise.

Once the papers were collected from the search engines, the most important studies were chosen by the criteria for inclusion and exclusion.

#### Inclusion criteria

The publications considered in this systematic review satisfied the criterion of utilising multimodal medical data for XAI. Furthermore, they satisfied at least one of the subsequent criteria:

- They employ a strategy or technique to visualise the model using multimodal medical data.
- They evaluate and contrast different techniques or methodologies for explainability in a multimodal setting.

#### Exclusion criteria

To reject all possible irrelevant publications retrieved, additional exclusion criteria were added to those implicitly defined by the search strategy and inclusion criteria. Hence, the publications that were excluded from this systematic review fulfilled at least one of the following conditions:

- The publication does not deal with medical XAI.
- Multimodal data types have not been considered.
- The publication is a short version of another retrieved publication.
- The publication has not been peer-reviewed.
- The publication is not written in English.
- The publication has worked on audio/video data.

### C Taxonomy

The rest of the paper are structured into the following sections (Figure 3). Section 2 covers the background about multimodal medical data and XAI methods. In Section 3, different approaches to modelling and visualisation have been discussed. Section 4 explores different multimodal frameworks in XAI data visualisation. The current practices, challenges, and future directions in the XAI data visualisation are presented in Section 5 and finally, the conclusion in Section 6. Table 2 presents the abbreviations and their full forms used throughout the presented literature.

**Table 2.**
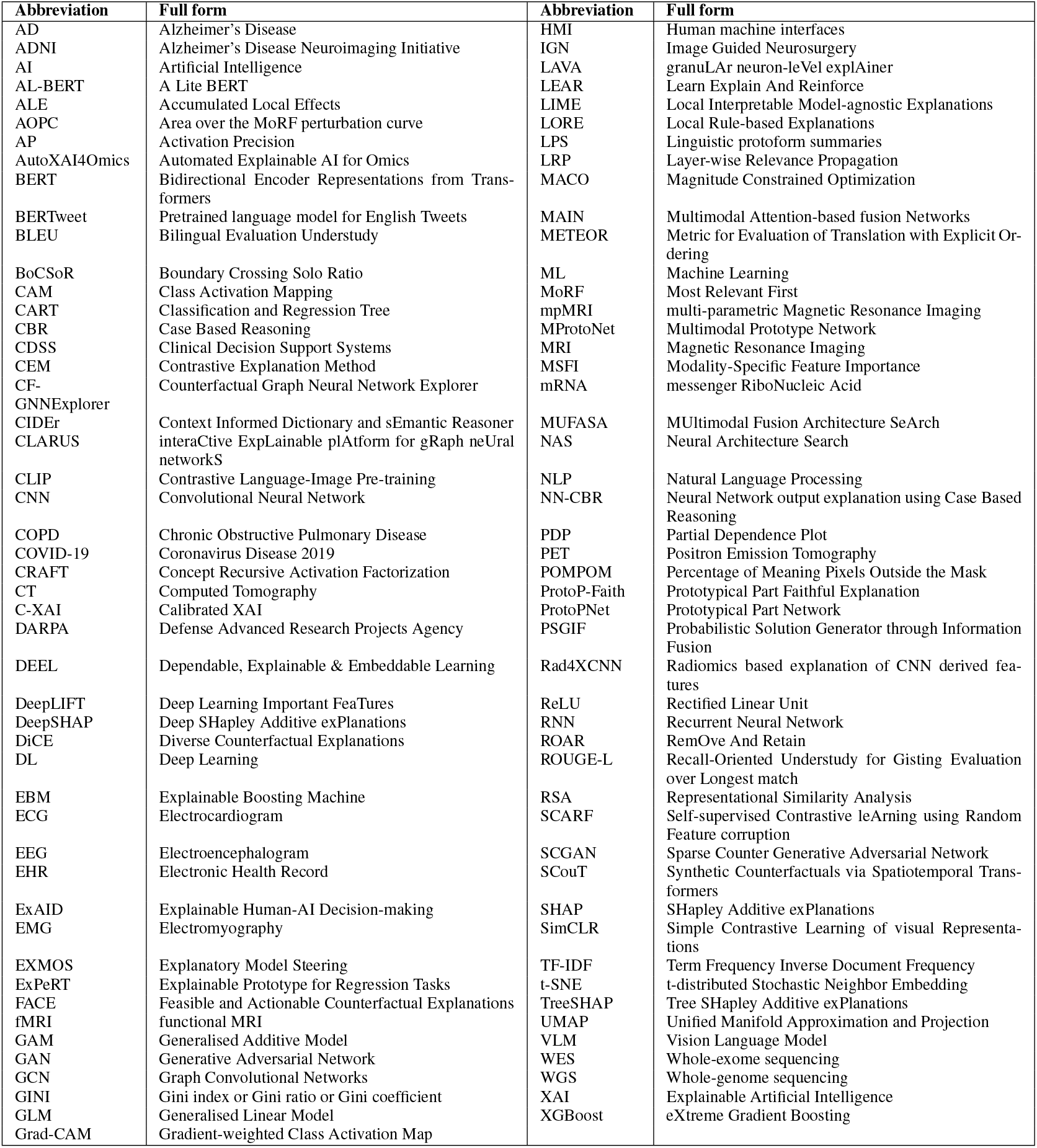
Abbreviations and their full forms used in the literature

**Fig. 3.**
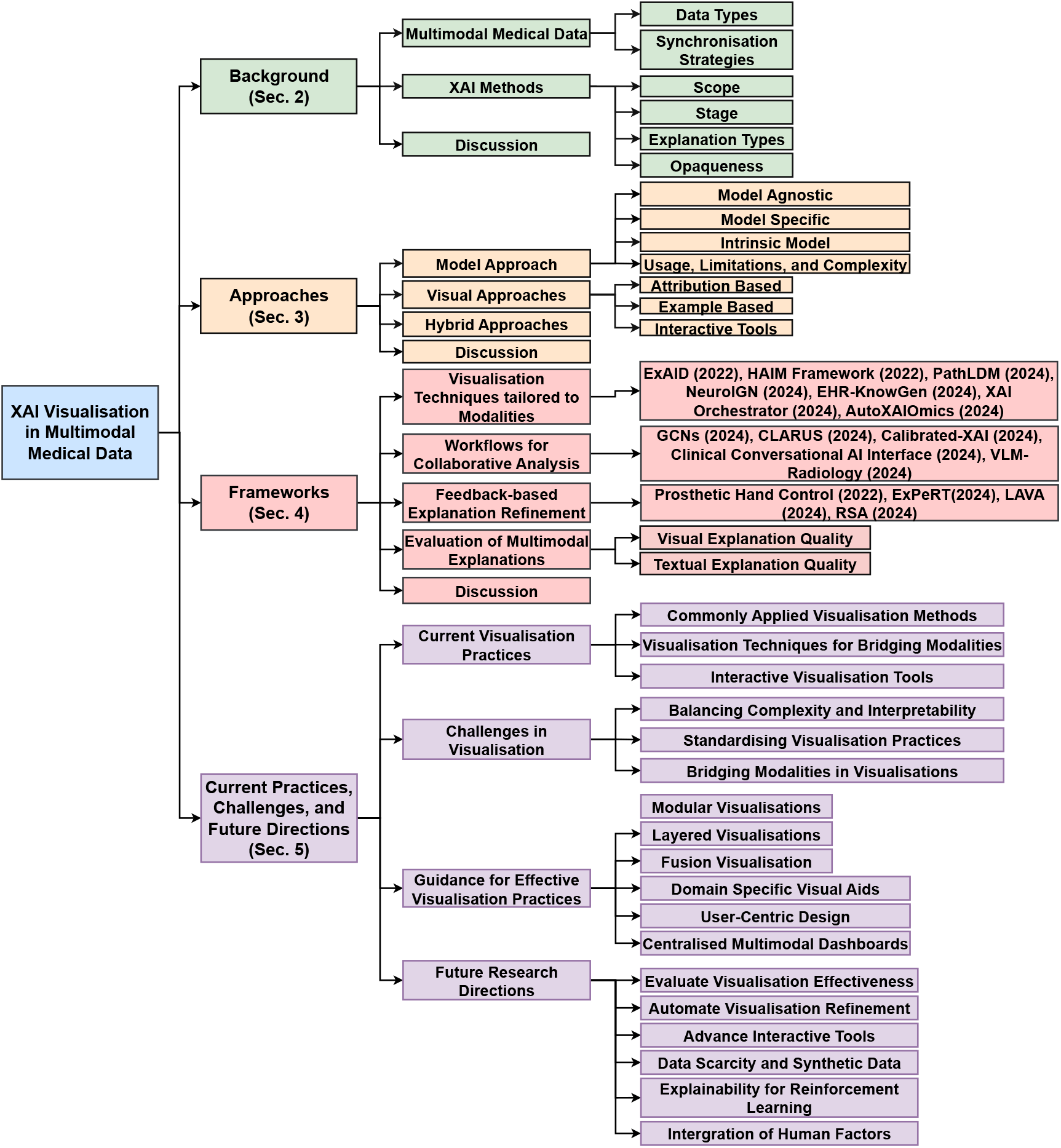
Taxonomy of XAI Visualisation in Multimodal Medical Data

## 2 Background

### A Multimodal Medical Data

Various types of data are used in clinical practice [34, 35] as shown in Figure 4 and Table 3, which highlight their relevance to machine learning representations. These cover various kinds of data, such as EHR, behavioural, omics, sensor, and imaging data. We present a comprehensive summary of the primary data modalities employed in modelling research to emphasise the importance of data selection in multimodal machine learning.

**Table 3.** General Overview of Data Types, Representations, and ML/DL Techniques in Healthcare

| Category | Subcategory | Examples | Representation for ML/DL | Common Models/ Techniques | Applications | Anatomical Locations |
| --- | --- | --- | --- | --- | --- | --- |
| EHRs | Numerical Data | Vital Signs (BP, HR, BMI), Lab Results, Dosages [36, 37] | Tabular, normalised values | Decision Trees [38], Regression, DNNs, Temporal Convolutional Networks [39] | Risk prediction, trend analysis | Heart (BP, HR), Body (BMI), Blood |
|  | Categorical Data | Patient Demographics, ICD Codes, Medication Categories [36] | One-hot encoding, embeddings | Logistic Regression, Neural Embeddings | Disease classification, population profiling | Organs, Systems (Cardiovascular, Metabolic) |
|  | Event Sequence Data | Admissions Logs, Disease Progression Timelines [40] | Temporal event embeddings | RNNs, Transformers, Sequence Models | Temporal modeling, disease pathways | Organs (Liver, Lungs, Heart) |
|  | Textual Data | Clinical Notes, Discharge Summaries [37] | Tokenised text, BERT embeddings | NLP Models (BERT, GPT), Sequence Models | Medical NLP, summarisation, diagnostics | Organs, Body parts (Context-specific) |
| Behavioural Data | Therapy Logs | Patient Diaries [41], Interaction Logs | Time-stamped sequences | RNNs, Transformers | behavioural modeling, therapy adherence | Brain (Cognition, Therapy) |
|  | Sentiment Data | Therapy Notes, Text Sentiment [42] | Sentiment embeddings, Bag-of-Words | Sentiment Analysis, RNNs | Psychological profiling, patient engagement | Brain (Emotional Regulation) |
|  | Social Interaction Data | Digital Communications [43, 44], Chat Logs [45] | Text embeddings, interaction graphs | GNNs, NLP Models | Social behaviour modeling, stress detection | Brain (Neurological Stress) |
| Omics | Genomics/ Proteomics | Gene Sequences [46], Protein Levels | One-hot sequences, k-mer embeddings | GNNs, Feature-Based Models | Biomarker discovery, gene-drug interactions | Cellular (DNA, Proteins) |
|  | Epigenomics | Methylation Profiles [47], Histone Data | Epigenetic feature matrices | Feature-Based Models, Neural Networks | Gene regulation, cancer research | Cellular (Epigenetic Markers) |
|  | Metabolomics/ Lipidomics | Metabolite Profiles [48], Lipid Biomarkers | Tabular, high-dimensional data | PCA, DNNs, Clustering | Metabolic disorder research, cardiovascular health | Liver, Heart, Blood |
|  | Spectroscopy Data | SERS [49, 50], Mass Spectrometry [51] | High-dimensional spectral features | Feature Selection, Random Forests, DNNs | Protein interaction mapping, molecular insights | Proteins, Tissues, Blood |
| Sensor Data | Physiological Signals | ECG, EEG, PPG, Wearable Streams [36] | Time-series signals, FFT-transformed features | RNNs, Transformers | Anomaly detection, physiological monitoring | Heart (ECG), Brain (EEG), Skin |
|  | Wearable Fusion Data | Accelerometer, Gyroscope, Multimodal Streams [36] | Multimodal embeddings | Multimodal Networks, Late Fusion | Comprehensive health monitoring | Musculoskeletal System, Limbs |
|  | Environmental Data | GPS [52], Air Quality [53], UV Levels [54] | Spatial embeddings, time-series signals | Geospatial Models, Transformers | Exposure analysis, epidemiology | Lungs (Air Quality), Skin (UV Exposure) |
| Medical Imaging | Standard Imaging | X-rays [55], CT Scans, MRI | Pixel arrays, preprocessed DICOM files | CNNs, Pretrained Models (ResNet, VGG) | Disease detection, lesion segmentation | Chest (X-rays), Brain (MRI), Abdomen |
|  | Advanced Imaging | Hyperspectral [56], Photoacoustic Imaging [57] | Spectral voxel arrays | 3D CNNs, Multispectral Networks | Tissue diagnostics, cancer detection | Skin (Dermatology), Breast (Cancer Detection) |
|  | Emerging Modalities | Optical Coherence Tomography (OCT) [58, 59], Thermal Imaging [60] | Voxel-based 3D arrays, heatmaps | 3D CNNs, GANs | Ophthalmology, inflammation detection | Eye (OCT), Skin (Thermal Imaging) |

**Fig. 4.**
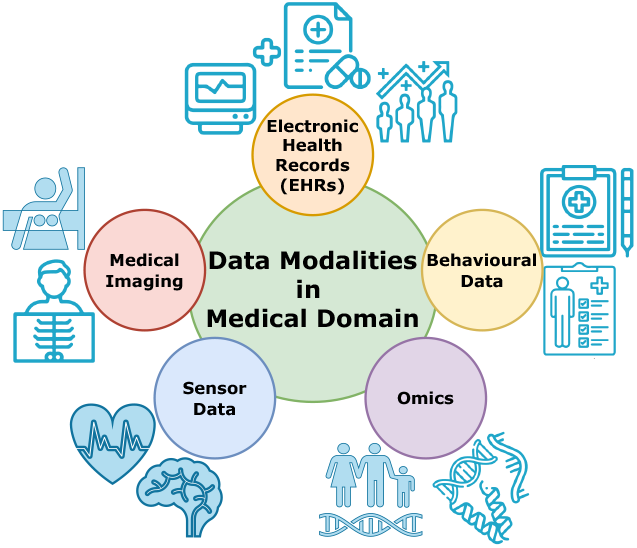
Data Modalities for Medical Domain

#### A.1 Data Types

##### EHRs

Electronic Health Records (EHRs) are digital representations of patients’ paper charts, containing a comprehensive compilation of patient health information generated through different interactions in the healthcare environment. Healthcare providers have adopted Electronic Health

Records (EHRs) as a central repository for medical data, resulting in a significant increase in data complexity [61]. Even though these datasets are large and specific to each patient, they are often disintegrated and poorly organised. This lack of integration complicates the analysis, especially because of the variety of data such as drugs, lab results, physiological assessments, and history records [62]. Machine Learning (ML) solves EHR dataset complexity by exploring complex correlations among varied factors [63].

EHRs are essential for multimodal integration in intelligent healthcare, particularly for life-threatening disorders like diabetes [64]. Managing and analysing unstructured data from sensors and EHRs is difficult. Accurate forecasts require data fusion, and deep learning is effective for larger healthcare datasets. Wearable sensors and EHRs can improve healthcare datasets by gathering patient data. Using an ensemble ML method, a recommendation system is created to anticipate and provide timely recommendations for multidisciplinary diabetes patients accurately. MUltimodal Fusion Architecture SeArch (MUFASA), a novel Neural Architecture Search (NAS) extension, was proposed by Xu et al. [65] to maximize multimodal fusion techniques in EHR data. The Transformer architecture has shown potential in utilising EHR internal structure. MUFASA outperformed Transformer, Evolved Transformer, RNN versions, and classic NAS models [66] in public EHR data experiments. Medical codes and clinical notes have distinct properties, making EHR data representation difficult. Inter-modal correlations, which models sometimes miss, are another difficulty. An et al. [67] introduced the Multimodal Attention-based fusIon Networks (MAIN) model to tackle significant healthcare prediction difficulties utilising EHR data. The MAIN model uses independent feature extraction modules for each modality, such as self-attention and time-aware Transformer for medical codes and a CNN model for clinical notes. The module includes a low-rank multimodal fusion approach and a cross-modal attention layer for inter-modal correlation extraction. To predict diagnosis, the model uses attention mechanisms and neural networks to aggregate modality representations and correlate them to build visit and patient representations [68]. MAIN provides a comprehensive multimodal fusion and correlation extraction platform in EHR-based prediction tasks. Vital sign features like blood pressure (BP) and oxygen levels are critical for in-hospital mortality prediction utilising XAI methods to realise their significance [37].

##### Behavioural Data

In healthcare, behavioural data is indispensable because it provides invaluable insights into individuals’ lifestyles, routines, and behaviours, substantially impacting their overall health and well-being. Behavioural data comprises a wide range of human behavioural aspects, such as compliance with medical treatments, sleep patterns, dietary practices, stress levels, and social interactions [69]. Using behavioural data, intelligent healthcare systems can continuously observe and assess the actions of individuals, enabling medical practitioners to gain a holistic comprehension of their patients’ daily routines and inclinations [70]. By utilising this information to identify patterns, trends, and deviations from typical conduct, it becomes possible to detect prospective health problems early and implement interventions promptly. Behavioural data can be utilised to monitor stress levels and emotional well-being, allowing healthcare professionals to recognise triggers and patterns that potentially affect the mental health of individuals [71]. This data has also been utilised to inform the creation of relaxation techniques, stress management methods, and individualised interventions aimed at assisting people in preserving their mental health [72]. In addition, behavioural data has the potential to enhance patient involvement and enable self-care. By utilising feedback mechanisms and interactive platforms, users can actively monitor their behaviours, set objectives, and receive personalised recommendations informed by the data they have provided. Due to this empowerment, individuals who manage their health and well-being may experience increased motivation and accountability. However, acquiring and analysing behavioural data raises significant ethical concerns like privacy, data security, and informed consent. Ensuring the protection of individuals’ privacy, storing their data securely, and obtaining appropriate consent for data collection and utilisation are all critical obligations.

##### Omics Data

Omics data refers to the collective information obtained from several sectors, such as genomics, proteomics, metabolomics, and other related disciplines [73]. This data type offers extensive knowledge about health and disease’s molecular and genetic foundations, facilitating individualised medicine and precise therapies. Genomics analyses an organism’s entire DNA sequence, encompassing all its genes, which is crucial for understanding the genetic predispositions associated with different diseases. Proteomics explicitly examines proteins, essential components of living organisms and vital bodily functions [74]. Metabolomics studies metabolites, tiny chemicals that play a role in metabolism, allowing a quick overview of an organism’s physiological condition [75].

The utilisation of omics data has resulted in notable progress in identifying biomarkers for diseases, understanding disease mechanisms, and formulating novel therapeutic approaches [76]. Whole-genome sequencing (WGS) and whole-exome sequencing (WES) have been crucial in finding genetic variants linked to different diseases. Integrating genomic data with clinical data in cancer research has facilitated the advancement of precision oncology, which customises treatments based on the genetic characteristics of an individual’s tumour. Furthermore, multi-omics data, which merges data from several omics disciplines, has yielded a more profound understanding of the intricate relationships within biological systems, enabling more efficient disease prevention and management techniques [77].

The contribution of omics data to personalised medicine is remarkable, as it enables the tailoring of healthcare according to an individual’s distinct genetic composition. Implementing this individualised strategy can enhance the effectiveness of therapies and decrease the probability of adverse side effects. Moreover, prominent endeavours like the 100,000 Genomes Project in the UK [78] and the Precision Medicine Initiative’s All of Us Research Program in the USA [79] exemplify the worldwide effort to utilize omics data to advance healthcare. These programs aim to establish comprehensive genomic databases that may be used to identify genetic variations associated with diseases and develop more efficient, individualised therapies.

##### Sensor Data

The ability of sensor data to measure and monitor a vast array of physiological parameters and activities in real-time is crucial to revolutionising healthcare. Various technologies, including wearable devices, implantable sensors, and remote monitoring systems, provide this information. Wearable devices, capable of being worn externally or incorporated into apparel and accessories, play a crucial role in collecting up-to-the-minute information regarding health-related metrics such as vital signs, physical activity, and sleep patterns [80, 81]. The knowledge obtained from this information is critical, as it enhances the overall understanding of an individual’s well-being and enables individualised surveillance and proactive healthcare approaches [82].

The continuous flow of sensor data is crucial for ongoing health assessment in the healthcare industry, as it enables timely intervention and early detection of prospective health problems [83]. In addition, the implementation of ubiquitous technologies permits remote patient monitoring, thereby expanding the scope of modern healthcare. This functionality enables healthcare practitioners to facilitate timely interventions when required [84]. The uninterrupted integration of intelligent healthcare systems with sensor data from wearable devices improves the timely identification of potential health complications. The integration is crucial for delivering timely, individualised interventions and recommendations, notably enhancing the quality of patient care and health results [85].

Utilising this up-to-date data stream empowers healthcare practitioners to formulate individualised treatment strategies that effectively attend to the specific requirements of each patient. For example, sensor data is critical in managing chronic ailments like cardiovascular diseases and diabetes by surveilling heart rate variability and glucose levels. In addition, using real-time sensor data enables telemedicine and telehealth services to expand, enabling medical practitioners to oversee and control patient conditions from a distance [86]. This feature provides essential healthcare services to individuals residing in remote areas or with limited mobility, thereby minimising the need for frequent hospital visits [87].

Furthermore, the healthcare industry relies heavily on the data management lifecycle, encompassing essential phases such as acquiring raw data, structuring data, fusing data, and developing predictive models to optimize the usefulness of the gathered information. By following this systematic procedure, the data obtained from sensors, wearable devices, and electronic health records (EHRs) can be efficiently merged and evaluated, resulting in improved healthcare outcomes. The process depicted in Figure 5 exemplifies the all-encompassing methodology for managing data, forming the foundation of accountable AI for healthcare systems.

**Fig. 5.**
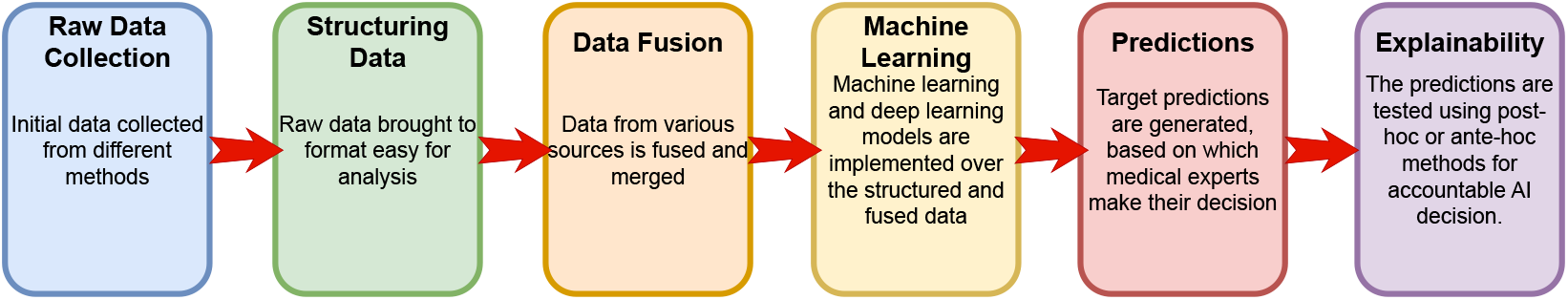
The lifecycle of raw data to XAI for multimodal medical data

##### Medical Imaging

Medical imaging is an essential component of modern healthcare systems, providing vital diagnostic information crucial for managing and treating various medical conditions [88–91]. Implementing advanced imaging technologies, including MRI, CT scans, PET scans, and ultrasound, permits the precise visualisation of anatomical components within the body [92]. This visualisation is critical in enabling healthcare practitioners to diagnose illnesses precisely, detect irregularities, and assess the effectiveness of treatments [93].

Integrating data-driven technologies and medical imaging streamlines the diagnostic process and enables a more individualised approach to patient care. By utilising predictive analytics to forecast disease progression and treatment response, it becomes possible to develop individualised therapeutic strategies that address specific health requirements. Therefore, the potential benefits of incorporating medical imaging and digital innovations into healthcare systems are substantial, as they guarantee improved patient outcomes and streamlined healthcare provision [94].

##### Multimodal Medical Data

Within healthcare, multimodal data fusion exhibits a significant change by integrating various forms of data to augment patient care and facilitate clinical decision-making [95]. Patients showing various symptoms in clinical practice are generally subjected to diagnostic tests utilising a variety of modalities to ascertain the underlying health conditions precisely. As a result, multimodal data fusion plays a crucial role in medical diagnostics, as it provides a solid foundation for integrating and analysing data from various sources. Various techniques, including early, hybrid, and late fusion (elaborated in subsection A.2), may be utilised to implement data fusion, depending upon the characteristics of the data and the particular clinical demands. Early fusion combines unprocessed data in the preliminary phase, providing a comprehensive dataset for subsequent examination. By merging characteristics at various processing phases, hybrid fusion balances the data’s comprehensiveness and the features’ specificity. In contrast, late fusion incorporates predictions or decisions from distinct models, capitalising on the unique attributes of each data source at the level of the decision.

This process is further improved by incorporating Explainable Artificial Intelligence (XAI) models, which generate outputs from fused data sets that are transparent and interpretable. XAI models facilitate healthcare professionals’ comprehension of the reasoning behind diagnostic recommendations by simplifying the decision-making processes of complex AI systems.

Integrating XAI into the multimodal data fusion process improves the precision of diagnoses and enables comprehension of the complex interconnections in the data. Implementing this methodology can substantially enhance the accuracy of disease detection and therapeutic strategy. Therefore, the combination of multimodal data fusion methodologies with XAI models signifies an advanced approach in the field of medical diagnostics, consistent with the main objectives of improving the quality of patient care and improving the credibility and dependability of AI implementations in healthcare environments. Table 4 provides an overview of various multi-modal medical datasets, including their access status (open or restricted). The table details the sample size and data modalities and includes hyperlinks for further information.

**Table 4.** Overview of Multimodal Medical Datasets

| Study | Country | Year | Data modalities | Access | Sample size | Link |
| --- | --- | --- | --- | --- | --- | --- |
| 100,000 Genomes Project [78] | UK | 2012 | Whole genome sequencing (WGS), Clinical data | Restricted | 100,000 genomes | genomicsengland.co.uk |
| American Gut Project [96] | USA | 2012 | Clinical, Diet, Microbiome | Open | ~25,000 | americangut.org |
| Atherosclerosis Risk in Communities (ARIC) [97] | USA | 1987 | Clinical data, Biomarkers, Genetic data, ECG data, Demographic data, Dietary history, Medication history, Environmental data, Administrative data | Restricted | ~15,792 participants at baseline | ahajournals.org |
| Avon Longitudinal Study of Parents and Children (ALSPAC) [98] | UK | 1991 | Questionnaires, Clinical data, Genetic data, Environmental data | Restricted | ~14,500 participants and their parents | bristol.ac.uk |
| Biobank Japan [99] | Japan | 2003 | Questionnaires, Clinical, Laboratory, Genome-wide genotyping | Restricted | ~270,000 | biobankjp.org |
| China Kadoorie Biobank [100] | China | 2004 | Questionnaires, Physical measurements, Biosamples, Genome-wide genotyping | Restricted | ~500,000 | ckbiobank.org |
| Cohorts for heart and aging research in genomic epidemiology (CHARGE) Consortium [101] | USA, Europe | 2009 | Genome-wide association studies (GWAS), Standardised health-related phenotypes, Clinical data | Restricted | ~38,000 individuals across 5 cohort studies | chargeconsortium.com |
| Framingham Heart Study [102] | USA | 1948 | Questionnaires, Clinical exams, Lab tests, Genetic data, Imaging, Omics data, Noninvasive and biomarker data | Restricted | ~15,000 across three generations | framinghamheartstudy.org |
| ImageCLEFmed 2019 [103] | Switzerland | 2019 | Medical images, Text annotations | Open | 70,786 images, 222,758 captions | imageclef.org/2019 |
| ImageCLEFmed 2020 [104] | Switzerland | 2020 | Medical images, Textual concepts | Open | 80,723 images, 416,180 captions | imageclef.org/2020 |
| ImageCLEFmed 2021 [105] | Switzerland | 2021 | Medical images, Captions | Open | 88,324 images, 427,205 captions | imageclef.org/2021 |
| ImageCLEFmed 2022 [106] | Italy | 2022 | Captions, VQA, Text | Open | 88,324 images and captions | imageclef.org/2022 |
| ImageCLEFmed 2023 [107] | Greece | 2023 | Medical images, Captions, Dialogue Summarisation, VQA | Open | 81,825 images. Non-radiological images removed | imageclef.org/2023 |
| ImageCLEFmed 2024 [108] | France | 2024 | Medical images (radiology), textual captions | Restricted | Training (70,108), Validation (9,972), Test (17,237) images | imageclef.org/2024 |
| Medical Information Mart for Intensive Care (MIMIC) [109, 110] | USA | 2008-2019 | Clinical/EHR, Images | Open | ~380,000 | mimic.physionet.org |
| MediCaT [111] | USA | 2020 | Medical images, Captions, Textual references | Open | 217,354 images and captions | github.com |
| Michigan Predictive Activity and Clinical Trajectories (MIPACT) [112] | USA | 2018-2019 | Wearables, Clinical/EHR, Physiological, Laboratory | Restricted | ~6,000 | researchtools.mipactstudy.org |
| Million Veteran Program [113] | USA | 2011 | EHR/clinical, Laboratory, Genome-wide | Restricted | 1 million | research.va.gov/mvp |
| Multimodal Brain Tumour Segmentation Challenge (BraTS) [114–116] | USA | 2018 | MRI scans, Segmentation masks, Clinical data | Open | ~2000 | med.upenn.edu/cbica/brats-2020/data.html |
| North American Prodrome Longitudinal Study [117] | USA | 2008 | Clinical, Genetic | Restricted | ~1,000 | napls.ucsf.edu |
| PMC-15M [118] | USA | 2024 | Medical images, Text | Open | 15 million image-text pairs | huggingface.co |
| PMC-OA [119] | USA | 2023 | Biomedical images, Text | Open | 1.6M image-caption pairs | huggingface.co |
| PMC-VQA [120] | USA | 2023 | Medical images, Textual Q&A | Open | 226,946 VQA pairs and 149,075 images | github.com/xiaoman-zhang/PMC-VQA |
| Precision Medicine Initiative's All of Us Research Program (PMI-AURP) [79] | USA | 2017 | Questionnaires, Social Determinants of Health (SDH), EHR/clinical, Laboratory, Genome-wide, Wearables | Open | 1 million (target) | allofus.nih.gov |
| Project Baseline Health Study [121] | USA | 2015 | Questionnaires, EHR/clinical, Laboratory, Wearables | Restricted | 10,000 (target) | projectbaseline.com |
| Radiology objects in context (ROCO) [122] | Germany | 2018 | Radiology images, Contextual data | Open | 81,500 images and captions | github.com/razorx89/roco-dataset |
| Radiology Objects in Context Version 2 (ROCOv2) [123] | Germany | 2024 | Radiology images, Contextual data | Open | 110,000 images and captions | zenodo.org/records/10821435 |
| Semantically-Labeled Knowledge-Enhanced Dataset (SLAKE) [124] | China | 2021 | Medical images, Textual Q&A | Open | 14028 images, VQA, Semantic Masks | huggingface.co/datasets/Bo-Kelvin/SLAKE |
| TOPMed [125] | USA | 2014 | Clinical, WGS | Open | ~180,000 | nhlbiwgs.org |
| The UK Household Longitudinal Study (UKHLS)<sup>7</sup> | UK | 2009 | Questionnaires, Clinical data, Biomarkers, Genetic data | Open | ~40,000 households (~100,000 individuals) | understandingsociety.ac.uk |
| UK Biobank [126] | UK | 2006 | Questionnaires, EHR/clinical, Laboratory, Genome-wide genotyping, whole-exome sequencing (WES), WGS, Imaging, Metabolites | Open | ~500,000 | ukbiobank.ac.uk |
| Visual Question Answering in Radiology (VQA-RAD) [127] | USA | 2018 | Radiology images, Textual Q&A | Open | 315 images, 3515 QA pairs | osf.io/89kps/ |

#### A.2 Synchronisation Strategies

Synchronisation strategies in multimodal machine learning refer to the techniques employed to fuse data from various sources or modalities shown as Disparate Health Data in Figure 6 to construct a unified and all-encompassing model. These methodologies guarantee the proper combination of information from many modalities to boost the model’s prediction capability. The primary methods for synchronisation in multimodal machine learning are early fusion, hybrid/intermediate fusion, and late fusion, each possessing unique attributes and uses.

**Fig. 6.**
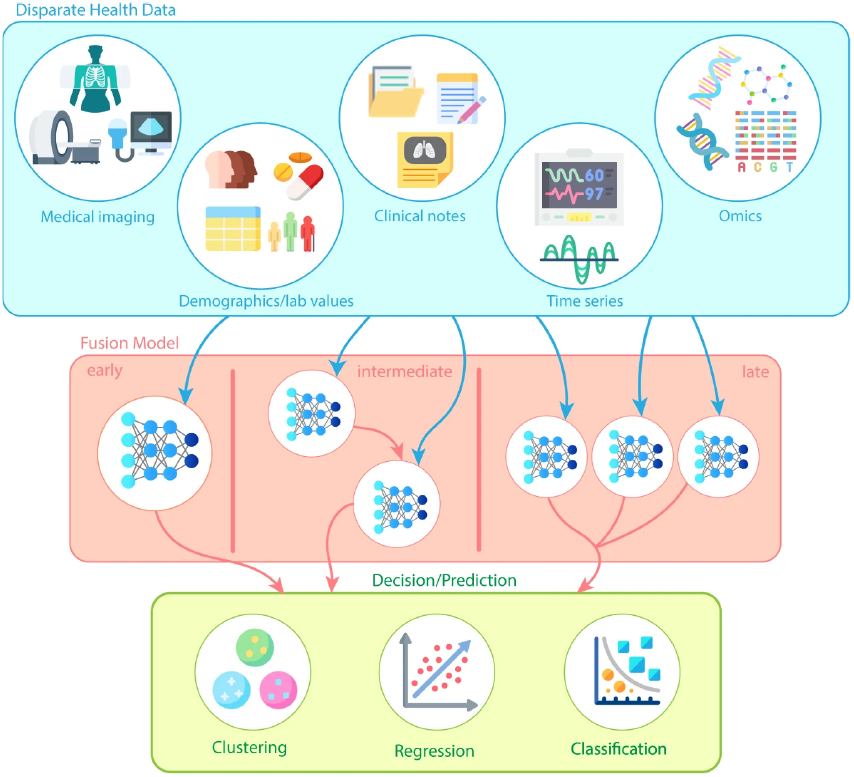
Illustration of how Early Fusion, Hybrid Fusion, and Late Fusion integrate diverse health data modalities - Medical Imaging, Clinical Notes, Demographics/Lab Values, Time Series, and Omics. Fusion strategies are applied at different stages: early input concatenation, intermediate feature-level integration, or late decision aggregation, supporting clustering, regression, and classification predictions [128]. This image is licensed under CC BY 4.0.

##### Early fusion

Also known as data-level fusion [129] or feature-level fusion [130, 131], early fusion precedes training of a machine learning model [128, 132, 133]. It combines multiple input modalities - Medical Imaging, Clinical Notes, Demographics and Lab Values, Time Series Data, and Omics Data in the preliminary phases (depicted in Figure 6). This methodology may employ unprocessed data or require a variety of feature extraction stages, ranging from straightforward aggregation to intricate model-driven extractions. The approach to feature combination generally differs according to the particular modalities and models employed. An illustration would be the aggregation of time series data for models such as XGBoost [134] or the clustering of multiple images as channels in a convolutional neural network (CNN) [135]. One of the primary obstacles encountered in early fusion methods for multimodal deep learning is effectively addressing the discrepancy in “data richness” between modalities [136]. To illustrate, in integrating vision and language data, light feature extractors are employed to transform the two sets of information into a compatible feature space. Nevertheless, vision data processing is frequently more intensive due to its complexities. This can create potential imbalances in which a model places an unbalanced emphasis on the vision data compared to the language data.

Furthermore, integrating the low-level features obtained from specific modalities may be challenging due to their potential lack of semantic richness. Word embeddings derived from language processing cannot directly interpret visual data, such as image boundaries, which are indispensable for computer vision models [137]. Standardising features across various modalities before their implementation in machine learning applications specific to tasks requires the development of modality-specific feature encoding strategies.

Across multiple domains, early fusion has demonstrated efficacy in clinical settings. For instance, An XGBoost model incorporating demographic data, lab results, image features, and vital signs was utilised to predict cardiomegaly [35]. Similarly, characteristics extracted from CT and PET scans have been integrated with demographic information to diagnose lung cancer [138], and MRI and PET images have been combined with genetic and demographic data to predict Alzheimer’s disease [139]. These applications underscore the potential of early fusion in augmenting the precision and effectiveness of healthcare predictive models.

##### Hybrid fusion

Hybrid fusion [140], or Intermediate fusion [128, 129], is a complex procedure comprising several stages. Initially, distinct models are employed to analyze data from various modalities to extract features. The concatenated features are input into the final model for prediction (see Figure 6). This methodology distinguishes itself from early fusion by individually optimising the features from each modality before combination. During training, feature representations are refined iteratively through each feature extraction model by back-propagating the loss function [132].

An often employed approach to intermediate fusion implementation entails pre-training models on specific modalities to develop resilient initial feature representations [35, 141]. Following the pre-training phase, it is customary to freeze the model weights and aggregate the outputs of all models. Training the final model on this concatenated dataset follows. Significantly, to authenticate as intermediate fusion, at least once during the training process, it is critical to de-freeze and modify a subset of the model weights to improve feature integration and learning across modalities [132].

Clinician-wide implementation of this technique is due to its strength in incorporating various data types. In oncology, for instance, hybrid fusion is utilised to predict cancer by combining pathological images with demographic data [142–144]. It enables incorporating X-ray images with demographic information, biomarkers, and clinical measurements in cardiology [145, 146]. In addition, it has demonstrated efficacy in neurology when demographic and clinical data are integrated with MRI images to forecast brain disorders [147, 148]. These applications demonstrate how hybrid fusion can substantially improve predictive accuracy through the utilisation of the distinct advantages of each data modality; consequently, this permits medical interventions that are more individualised and accurate.

##### Late fusion

The late fusion, alternatively referred to as decision-level fusion [149], is implemented once the individual models have made the predictions and their decisions are fused at the final stage either by an auxiliary model or an aggregation function (see Figure 6). This process ensures that each data source is handled independently while enabling a final, unified decision that takes advantage of the strengths of all modalities [150, 151].

An essential benefit of late fusion is its resilience when confronted with incomplete data sets. The Contrastive Language Image Pre-Training (CLIP) model undergoes inference with associated text data even when pre-trained on image and text data and can operate efficiently as an image classifier [152]. Nevertheless, an undesirable aspect of late fusion is its failure to capture inter-modal interactions that occur during model training and feature extraction. This may lead to losing valuable information when inter-modal connections are vital [132, 153, 154]. Moreover, integrating outcomes from numerous models can present a complex undertaking [149].

Notwithstanding these obstacles, late fusion has been effectively implemented across diverse medical domains. For instance, combining cognitive assessment scores, demographic information, and predictions extracted from MRI scans have been utilised to diagnose cognitive impairments [155]. Combining biomarker data with magnetic resonance image (MRI) analyses has similarly enabled late fusion to improve cancer prediction in oncology [143]. Furthermore, within the framework of the COVID-19 pandemic, late fusion techniques have been employed to correlate demographic and clinical indicators with CT scan results to predict the contraction and advancement of the virus [156, 157]. These applications demonstrate the strategic value of late fusion in scenarios that need modality-specific studies before a merged diagnosis. This ensures a flexible and comprehensive data-driven prediction strategy.

### B XAI Methods

Explainable Artificial Intelligence (XAI) approaches can be categorised according to different criteria. Figure 7 presents the classification of various XAI methods based on their scope, stage, types, and opaqueness.

**Fig. 7.**
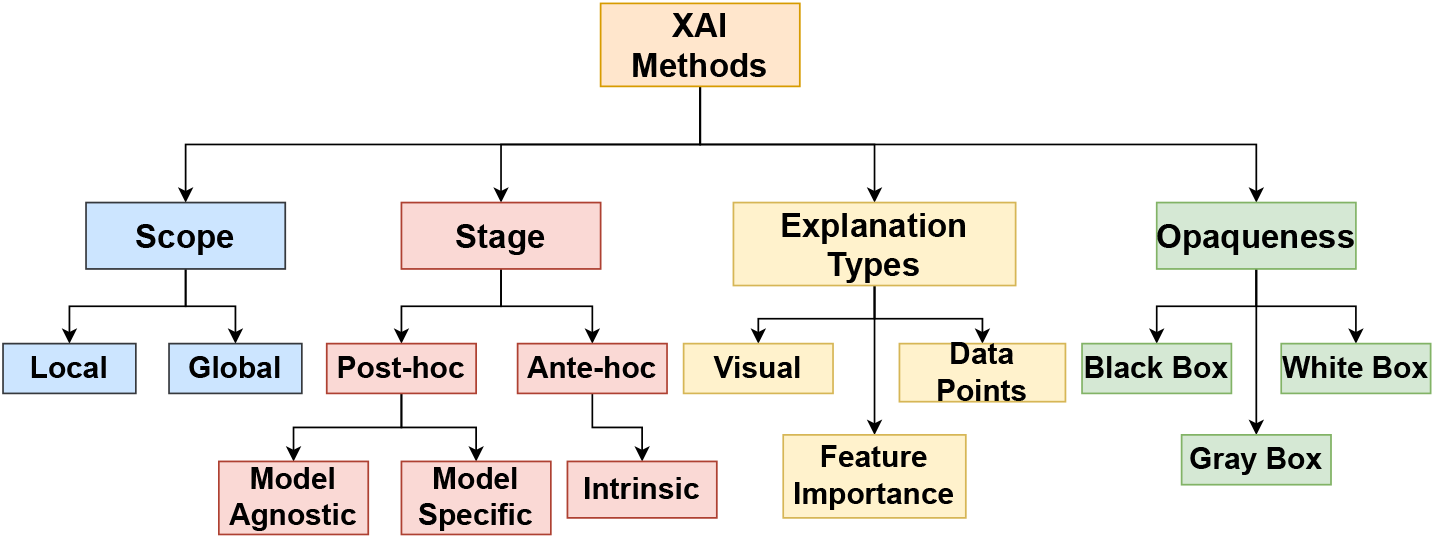
Various XAI Methods

#### B.1 Scope

The scope of their explanations can categorize XAI techniques. We present the local and global explanability in brief.

***Local*** explanations focus on explaining individual predictions of a model by understanding how a specific input leads to its particular output [158]. Using Local Interpretable Model-agnostic Explanations *(LIME)*, medical imaging professionals can focus on the essential parts of a chest X-ray image that impact the diagnosis [159]. For example, LIME could show that areas with apparent opacities are critical for pneumonia diagnosis [160]. SHapley Additive exPlanations *(SHAP)* can explain individual predictions by computing the contribution of each feature. For example, it can explain the prediction for diabetes in a particular patient by identifying relevant features like glucose levels and age [161].

***Global*** explanations seek to offer a thorough comprehension of the overall behaviour of the entire model [158]. Decision trees provide a model structure that is easy to understand, making them appropriate for analysing how patient characteristics impact the diagnosis of illnesses such as breast cancer in the total patient population [162]. In deep learning, models applied to multimodal medical datasets, such as combining MRI images with clinical data, feature importance rankings provide insights into how the model assigns priority to different data sources when making decisions [163].

#### B.2 Stage

This subsection discusses explainable techniques based on the timing of their execution in the machine learning model building lifecycle. A post-hoc technique uses the explainable AI (XAI) model after the primary model has completed its training. Alternatively, when the XAI model is incorporated during the creation of the main model, it is referred to as an ante-hoc approach [164].

##### Post-hoc

Post-hoc techniques are applied after the primary model training. These strategies are essential for comprehending and analysing the decisions made by intricate models once their training period has concluded. Post-hoc strategy can be categorised into two types: model-agnostic and model-specific approaches.

*Model Agnostic* approaches apply to models of any type. *LIME* is a technique that can be applied to explain models that make predictions about diabetic retinopathy using fundus images and clinical data, regardless of the specific model being employed [165, 166]. *SHAP* can explain a multimodal system that utilises MRI and genetic data to forecast the evolution of Alzheimer’s disease [167, 168] as well as for Parkinson’s disease [169].

*Model Specific* approaches are tailored to specific model architectures. When applied to multimodal medical data, the Contextual Attention Model *(CAM)* improves the interpretability of black-box models [170]. By incorporating attention mechanisms into neural networks, CAM accurately recognises significant elements of various data inputs, including clinical metrics and imaging data that are taken into account by a model during diagnosing or predicting. This is done by generating visual representations such as heat maps, which emphasize crucial regions in the data that substantially influence the model’s outputs.

Gradient-weighted Class Activation Mapping *(Grad-CAM)* is a technique employed with convolutional neural networks *(CNNs)* to generate visual heat maps. Pathology uses this technique to emphasize areas relevant to identifying malignancy in histopathology images. *TreeExplainer* is a form of SHAP specifically designed for tree-based models like XG-Boost. TreeExplainer can explain the predictions made by a gradient-boosted model trained on clinical characteristics to predict heart disease. Another model-specific method is the eXirt (Explanations based on Item Response Theory) technique [171], which employs Item Response Theory (IRT) to evaluate feature significance in tree-ensemble models. eXirt systematically perturbs input features, treated as test items, to quantify their impact on model decisions. It offers global insights, including feature rankings across patient populations and instance-specific explanations. This approach is especially advantageous for multimodal medical datasets, where comprehending the interplay between clinical data, imaging biomarkers, and genetic information is essential. Detailed discussion about model-agnostic and model-specific XAI techniques have been presented in Section A.

##### Ante-hoc

In contrast, ante-hoc strategies incorporate explainability into the model as it is being developed. These inherent models possess inherent interpretability, providing trans-parency and comprehensibility from the beginning without requiring supplementary explanatory techniques.

*Intrinsic Models* Unlike model-agnostic or model-specific techniques, intrinsic models automatically incorporate explainability within their architecture. The inherent transparency of this feature enables users to comprehend the reasoning behind forecasts directly from the model’s structure and operation without the necessity for further explanatory methods. Some instances of intrinsic models in healthcare have been presented as follows.

*Linear models* are widely employed in medical research to categorize risks and forecast diseases using clinical data [172, 173]. *Logistic regression* can forecast the probability of a patient acquiring hypertension, considering characteristics such as age, weight, and cholesterol levels. The model’s coefficients offer direct insights into the significance and influence of each element on the outcome [174–176]. *Decision trees* are considered intrinsic models because of their straightforward, rule-based structure that divides data into branches based on specific conditions. A decision tree can be employed in healthcare to diagnose patients by analysing their symptoms. For example, a tree can classify respiratory symptoms to identify illnesses like asthma [177, 178] or chronic obstructive pulmonary disease (COPD) [179, 180], with each decision point providing a logical explanation for each branch.

The Explainable Boosting Machine (*EBM*) is an interpretable machine learning model. It integrates the transparency required in critical domains like healthcare with the precision of gradient boosting. EBMs represent the additive contribution of each feature to the final decision [181]. This simplifies the process of decision-making. This is advantageous in healthcare, where clinicians must incorporate and interpret multimodal medical data to make informed decisions. Such data may include clinical notes, patient demographics, laboratory results, and medical images. EBMs can, for instance, process and incorporate features from various data types to predict heart failure [182–184]. This includes features extracted from ECG images, NLP-processed clinical notes, and direct numerical inputs from lab results [185].

This approach facilitates the model’s efficient handling of diverse and complex data and offers transparent insights into the individual impact of each factor.

The openness of intrinsic models is highly valued as it promotes confidence and enables more straightforward validation and regulatory approval, particularly in sensitive industries such as healthcare. They provide a counterpoint to more intricate models, such as deep neural networks, which frequently have unclear decision-making processes and necessitate external explanation techniques to clarify their process.

#### B.3 Explanation Types

Explanation types are classified based on how explanations can be conveyed.

##### Visual

Explanations are communicated through images or visual depictions. Saliency maps are used in dermatology to detect skin lesions. They visually represent the areas in a dermatology image that contribute to a model’s prediction of melanoma [186, 187]. Grad-CAM utilises heatmaps to superimpose retinal images to identify the regions highly symptomatic of diabetic retinopathy throughout the diagnostic process [188–190].

##### Feature Importance

Explanations are derived by identifying the key features that impact the predictions more. Permutation importance is a metric that may be used to measure the extent to which changing each feature affects the performance of a model in predicting the risk of cardiovascular disease using both imaging and clinical data [191–193]. When radiological images are paired with patient metadata, feature importance scores can demonstrate that clinical factors such as age or gender are frequently just as important as image features [194, 195].

##### Data Points

Using explanation by data points is a highly effective method for determining the extent to which each data point contributes to the prediction. This method uses the visualisation of the data points to identify key features and data points that form clusters or are outliers, giving more clarity about the data point’s contribution to the prediction. One can modify the data points with the what-if tool^8^ to generate counter-factual explanations, as this tool helps understand model’s sensitivity to a variety of features. Data points technique can help diagnose lung diseases using CT scans [196, 197]. When forecasting the risk of breast cancer by combining mammography and genomic data, counterfactual explanations can demonstrate how altering genetic markers could potentially impact a patient’s risk classification [198, 199].

#### B.4 Opaqueness

##### Black Box Models

The complexity and opacity of these models make it challenging to interpret their predictions directly. Deep neural networks, random forests, and other intricate machine learning models are classified in this category. Posthoc explanation techniques, such as saliency maps (visual representations), feature importance rankings, and surrogate models (such as LIME and SHAP), are employed to explain black-box models.

##### Gray Box Models

Gray box models are an intermediary alternative to black and white box models. They are helpful when dealing with multimodal medical data. These models strike a harmonious equilibrium by providing clarity that enables researchers and clinicians to comprehend the decision-making process without delving into the intricacies of complex systems. For example, when examining the correlation between EHRs and medical images, gray box models may incorporate ensemble models or simplified neural networks [200–203]. By enabling users to trace the impact of particular patient data points on diagnostic outcomes, these configurations maintain a degree of interpretability regarding the significance of features or the influence of various data modalities [204]. Ensuring this degree of transparency is of the utmost importance in medical environments, where comprehension of the reasoning behind an automated system’s determination can significantly influence treatment decisions and patient confidence. These methods provide limited comprehensibility while retaining a certain level of intricacy. Examples are less complex iterations of neural networks or ensembles, wherein the interpretability of feature importance or specific model architecture is preserved.

##### White Box Models

Due to their renowned transparency, white box models are frequently implemented in environments where it is critical to comprehend the decision-making process, such as healthcare with multimodal medical data. These models comprise basic machine learning algorithms such as decision trees, linear models, and rule-based systems, which are inherently clear-cut and, therefore, “interpretable by design” [205, 206]. The clarity of this transparency is especially beneficial in the context of XAI for medical applications, as it enables healthcare practitioners to observe precisely how various data categories, including lab results, patient histories, and images are processed and how they factor into the formulation of diagnostic conclusions. Nonetheless, white box models may also be constrained by their simplicity. Potentially less accurate results could result from their inability to process complex data patterns as efficiently as more sophisticated models in situations involving intricate multimodal data interactions. Although white-box models are transparent, their use in healthcare should be carefully balanced with the need for deep analysis to handle the complexity of medical data effectively.

Various XAI models have been listed and classified based on their scope, stage, explanation type, opaqueness, and if they are agnostic in Table 5.

**Table 5.** Overview of XAI Models

| XAI Model | Scope | Stage | Explanation Type | Opaque | Agnostic |
| --- | --- | --- | --- | --- | --- |
| CART [207] | Global | Ante-hoc | Rule-based, Visual | White | No |
| Decision Trees [208] | Global | Ante-hoc | Rule-based, Visual | White | No |
| Explainable Boosting Machine (EBM) [181] | Global | Ante-hoc | Feature Importance | White | No |
| J48 [209] | Global | Ante-hoc | Rule-based, Visual | White | No |
| SimCLR [210] | Global | Ante-hoc | Feature Importance | Gray | No |
| Transparency by Design (TbD) [211] | Global | Ante-hoc | Rule-based | White | No |
| ALE [212] | Global | Post-hoc | Feature Importance | Gray | Yes |
| DIME [213] | Global | Post-hoc | Feature Importance, Visual | Gray | Yes |
| PDP [7] | Global | Post-hoc | Feature Importance | Gray | Yes |
| CAM [214] | Local | Ante-hoc | Visual | Gray | No |
| Activation Maximization [215] | Local | Post-hoc | Visual | Gray | No |
| Anchor Explanations [216] | Local | Post-hoc | Rule-based | Black | Yes |
| CBR [217] | Local | Post-hoc | Example-based | Gray | Yes |
| CEM [218] | Local | Post-hoc | Example-based | Gray | Yes |
| CF-GNNExplainer [219] | Local | Post-hoc | Feature Importance | Gray | No |
| CLARUS [220] | Local | Post-hoc | Example-based | Gray | Yes |
| Concept Attribution [221] | Local | Post-hoc | Feature Importance, Visual | Gray | No |
| C-XAI [222] | Local | Post-hoc | Feature Importance | Gray | No |
| DeepLIFT [223] | Local | Post-hoc | Feature Importance | Gray | No |
| DeepSHAP [224] | Local | Post-hoc | Feature Importance | Gray | Yes |
| DiCE [225] | Local | Post-hoc | Example-based | Gray | Yes |
| FACE [226] | Local | Post-hoc | Example-based | Gray | Yes |
| Grad-CAM [227] | Local | Post-hoc | Visual | Gray | No |
| Grad-CAM++ [228] | Local | Post-hoc | Visual | Gray | No |
| ICE [8] | Local | Post-hoc | Feature Importance | Gray | Yes |
| LIME [229] | Local | Post-hoc | Surrogate Models | Black | Yes |
| LORE [230] | Local | Post-hoc | Example-based, Rule-based | Black | Yes |
| Linguistic protoform summaries (LPS) [231] | Local | Post-hoc | Feature Importance | Gray | Yes |
| LRP [4] | Local | Post-hoc | Feature Importance | Gray | No |
| LSP [232] | Local | Post-hoc | Feature Importance, Visual | Gray | No |
| NN-CBR [233] | Local | Post-hoc | Example-based | Gray | Yes |
| Saliency Map [234] | Local | Post-hoc | Visual | Gray | No |
| SCGAN [235] | Local | Post-hoc | Feature Importance, Visual | Gray | No |
| Sharp-LIME [236] | Local | Post-hoc | Feature Importance, Visual | Gray | Yes |
| Smooth-Grad [237] | Local | Post-hoc | Feature Importance | Gray | Yes |
| Smooth-GradCAM++ [238] | Local | Post-hoc | Visual | Gray | No |
| Integrated Gradients [239, 240] | Local, Global | Post-hoc | Feature Importance | Gray | No |
| KernelSHAP [224] | Local, Global | Post-hoc | Feature Importance | Gray | Yes |
| SHAP [6] | Local, Global | Post-hoc | Feature Importance | Gray | Yes |
| TCAV [241] | Local, Global | Post-hoc | Concept-based, Feature Importance | Gray | Yes |
| TreeExplainer [242] | Local, Global | Post-hoc | Feature Importance | Gray | No |
| TreeSHAP [243] | Local, Global | Post-hoc | Feature Importance | Gray | No |

## C Discussion

The extensive array of multimodal medical data - EHRs, behavioural data, omics data, sensor readings, and medical imaging presents considerable problems and valuable prospects for XAI. Every category of data possesses distinct characteristics that necessitate tailored explanatory methods. Structured EHR data can be explained using feature-based approaches like SHAP, which emphasise the impact of individual variables. Conversely, medical imaging frequently needs more visually intuitive techniques such as Grad-CAM, which produces heatmaps to highlight regions of interest within the image. The methods for combining these varied data types, namely early fusion, hybrid fusion, and late fusion introduce an additional layer of complexity. These fusion techniques facilitate the integration of information from many modalities, enabling researchers to harness the complementary advantages of each data source. The selection of fusion strategy critically influences the application timing and methodology of XAI techniques inside the data processing pipeline. The stage at which data is fused substantially influences the interpretability of the resultant models, with diverse methods providing differing degrees of transparency. This discussion establishes the foundation for investigating the application of XAI methodologies in complex, multimodal contexts. A persistent issue is a correspondence between the characteristics of the data and the selection of explanatory procedures, highlighting the necessity for adaptability in XAI methods to tackle the distinct challenges presented by the integration of varied forms of medical data. By accommodating these complexities, XAI can provide significant insights into models constructed from multimodal datasets.

## 3 Approaches

### A Model Approach

Selecting and utilising suitable model techniques in Explainable Artificial Intelligence (XAI) is essential for attaining transparency and interpretability in multimodal medical data. This section explores different model-based approaches, classifying them as model-agnostic, model-specific, and intrinsic models. Every category provides unique approaches to include explainability into AI systems, allowing doctors and researchers to have a deeper understanding of and more confidence in the results produced by complicated models. Through a thorough analysis of various methods, we aim to provide a comprehensive framework that improves the understandability of AI applications in healthcare. This will ultimately lead to better diagnostic accuracy and patient outcomes. The subsequent sections will thoroughly examine these categories, beginning with model-agnostic approaches.

#### A.1 Model Agnostic

Model-agnostic techniques are crucial in contexts where the internal mechanisms of models are either hidden or too complex to dissect directly, such as in medical fields using proprietary or advanced algorithms. The advantages of these methods lie in their flexibility across different models, their capability to offer explanations at varying levels of detail, and their adaptability to diverse data forms, including multimodal inputs [229].

These techniques are broadly categorised into surrogate models and feature interaction methods. Surrogate models are subdivided into local and global types. Local surrogate models create simplified versions of complex models to explain specific predictions, which is beneficial for detailed, patient-specific analyses. An example is *Local Interpretable Model-agnostic Explanations (LIME)* [229], which constructs straightforward models to elucidate individual decisions. On the other hand, global surrogate models use simpler models, such as linear models or decision trees, to approximate and thus demystify the functioning of more complex models. These models provide a macro-level understanding of a model’s decision-making processes. Metrics like the R-squared value have been evaluated to determine how well they replicate the variance of the original model’s outputs. This macro perspective is invaluable in healthcare, assisting in assessing AI systems’ reliability across various clinical applications.

Feature interaction methods, another category of model-agnostic approaches, quantify how changes in one or more variables influence predicted outcomes while keeping other variables constant. This includes techniques such as *partial dependence plots (PDP), individual conditional expectation (ICE), accumulated local effects (ALE)*, and *H-statistics*. These methods are beneficial for elucidating feature interactions in complex, multimodal medical datasets, such as the interplay between clinical biomarkers and imaging data, which could impact diagnostic algorithms. Their universal applicability makes them essential tools for clinicians and medical researchers seeking to validate and understand AI model predictions comprehensively.

Figure 8 organises the various model-agnostic XAI methods, underscoring their role in enhancing transparency and trust in AI-supported medical diagnostics and decision-making systems.

**Fig. 8.**
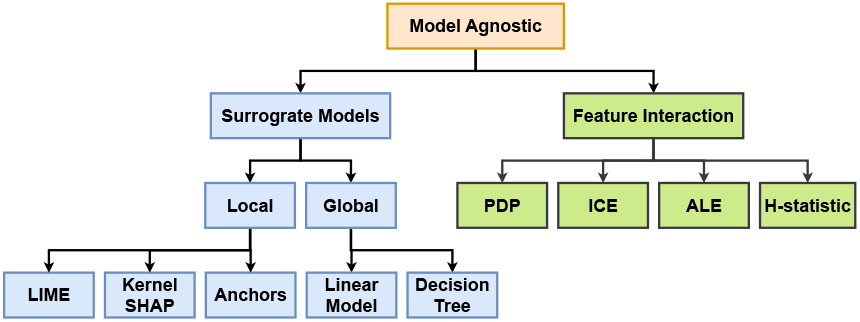
Various Model Agnostic XAI Methods

##### Surrogate Models

Surrogate models are simpler versions of complex machine learning models that estimate and interpret their predictions. They serve as interpretable alternatives, allowing users to comprehend the decision-making procedures of the original models without sacrificing their complexity or accuracy. Surrogate models are essential in XAI as they reduce complex data linkages and make them understandable to humans, facilitating explanations. The *LIME* method is a well-known approach that uses surrogate models to clarify the individual predictions made by a black-box model. The attribute level classification for the test sample is presented in Figure 9.

**Fig. 9.**
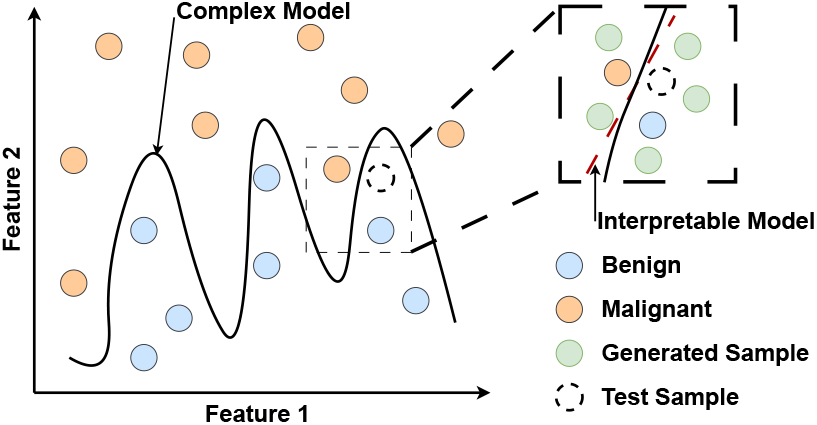
Complex model *f* decision boundaries are black curved lines. LIME explains a new test sample (dashed circle) by fitting an interpretable model *g* (red dashed line) to its variations (green circles), which are random samples. The fitted model shows how each attribute classifies that test sample. Adapted from [21]. This image is licensed under CC BY 4.0.

In *LIME* method, the aim is to elucidate the predictions of a complex model, represented as *f*, by employing an interpretable surrogate model *g*. This elucidation is specific to an instance *x*. The optimal surrogate model *g*^*^ is identified by minimising a function that includes a loss function ℒ (*f, g, π*_*x*_) and a complexity measure Ω(*g*). The loss function assesses how closely *g* can approximate *f* around *x*, using weights determined by the proximity of each perturbed data point *z* to *x*. The proximity is calculated with 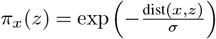, where distance is scaled by a parameter *σ*, which controls how quickly proximity influence diminishes with increasing distance.

If the surrogate model *g* is a linear model, it is expressed as 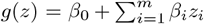. In this formula, *β*_0_ is the intercept, and *β*_*i*_ are the coefficients for the features *z*_*i*_, indicating how much each feature contributes to the prediction at *x*. This framework balances simplicity, enhancing interpretability and accuracy, ensuring closeness to *f*. Its model-agnostic nature allows it to be applied universally across different types of complex models *f*, providing a flexible tool for explanation in machine learning. Graziani et al. [236] proposed and utilised *Sharp-LIME*, which leverages nuclei annotations, to enhance the understandability and reliability of explanations provided by *LIME* in histopathology through improved segmentation techniques.

*SHAP* uses cooperative game theory to provide consistent and fair explanations for any machine learning model. The core concept relies on Shapley values [244], a solution concept in cooperative game theory that determines each player’s contribution to the overall outcome. The concept of SHAP is presented in Figure 10, illustrating how the binary classification of skin cancer is determined using five factors.

**Fig. 10.**
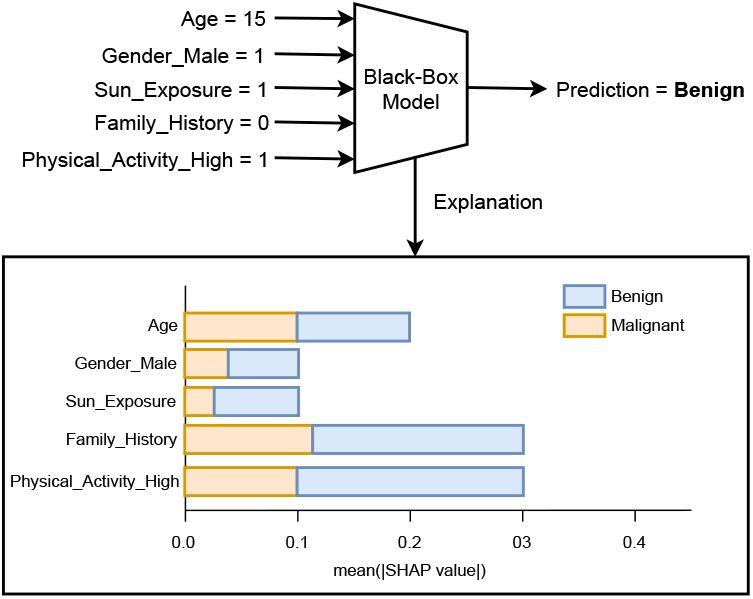
Given input of five factors including Age, Gender_Male, Sun_Exposure, Family_History, and Physical_Activity_High for Benign vs. Malignant Skin Cancer. Explanation shown using mean SHAP values for the given Benign prediction. Adapted from [21]. This image is licensed under CC BY 4.0.

The goal is to explain the output of a complex model, denoted by *f*, for a given instance *x* using the concept of Shapley value. This approach quantifies the contribution of each feature in the model to the prediction. The Shapley value for a feature *i* is computed as follows:

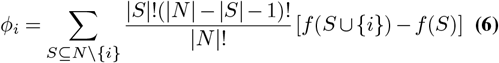

In this equation, *S* represents any subset of all features *N*, excluding the feature *i*. The terms *f* (*S*) and *f* (*S* ∪ {*i*}) refer to the model’s predictions when only the features in *S* are included and when both *S* and *i* are included, respectively. The factorial terms calculate the number of ways features can be ordered, thus weighting the importance of feature *i* relative to different subsets *S*.

For a model with five features, *x*_1_, *x*_2_, *x*_3_, *x*_4_ and *x*_5_, the total prediction *f* (*x*) can be broken down as:

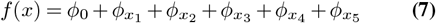

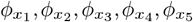 represent the contributions of the respective features based on their Shapley values. The Shapley value method ensures that the contributions are fairly distributed among all features based on their actual impact on the model’s output, reflecting the value added or subtracted by including each feature.

Among *LIME* and *SHAP, LIME* does not ensure fair distribution of the prediction impact across features. In contrast, the Shapley value method, on which *SHAP* is based, guarantees a fair allocation of effects across features underpinned by robust theoretical principles. The axioms of efficiency, symmetry, dummy, and additivity provide a strong and rational basis for these explanations [245].

*Anchor algorithms* The Anchor XAI technique is a model-agnostic approach that integrates black-and-white box methodologies elements. It is intended to explain the outputs of classifier models, including those employed in multimodal medical data AI systems. This method is based on identifying *“anchors”*, which are decision principles that firmly establish the rationale behind the predictions made by black-box classifiers. These anchors function similarly to IF-THEN rules, identifying regions within the feature space where predictions remain consistent despite changes in other feature values of the input instance [216]. These regulations facilitate a more profound comprehension of intricate AI systems that analyze a variety of data categories, including images, text, and structured data, and they also increase confidence in their decisions. Anchor XAI is classified as a grey box approach, which means it is independent of the internal mechanics of the models it clarifies. This is similar to black-box techniques, and it simultaneously offers clear, understandable decision rules that increase transparency and interpretability, attributes typically associated with white-box techniques. Anchor XAI enhances the interpretability of complex black box models without requiring modifications to the models or disclosing their detailed inner workings. Consequently, it is particularly advantageous for multimodal medical data AI systems applications.

##### Feature Interaction Methods

Gaining insight into the complex interaction of many components within a machine learning model is essential for comprehending and elucidating the model’s predictions. Feature interaction approaches offer insights into these interactions by displaying the correlations between features and the projections made by the model. This section examines essential tools, including Partial Dependence Plots (PDPs), Individual Conditional Expectation (ICE) plots, Accumulated Local Effects (ALEs), and H-statistics, that are crucial for displaying and evaluating the effects and interactions of features.

*Partial Dependence Plots (PDPs)* [7] are visual tools used to illustrate how the average prediction of a model changes as a particular feature’s value varies while keeping all other features constant. These plots are typically created for each feature in a dataset. Conceptually, the partial dependence curve provides the best estimate of a prediction if only one feature’s value is known. To generate a PDP for a specific feature, we use the following formula:

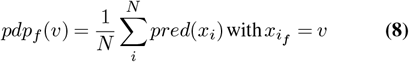

where *N* is the number of instances in the dataset, *pred* is the prediction function defined by the model, *f* represents the feature being analysed, *v* is a value within the feature’s domain, *M* is a hyperparameter that determines the explanation’s granularity, 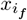 is the feature value of the *i*-th instance, set to the desired value *v*.

The drawback of the PDP is that the realistic maximum number of features in the partial dependence function is two. Other drawbacks are the feature independence assumption and heterogeneous effects might be hidden. In most cases, the trained models would have some inter-dependence of features. There might be a case for a particular feature that 50% of the data points have a positive impact, and another 50% harm the prediction. In that case, PDP, being the aggregated line, would be a horizontal line cancelling each other’s effect, and we might think that feature does not contribute to prediction. To overcome the limitations of Partial Dependence Plots (PDPs), which can mask the varying impacts across different data points, *Individual Conditional Expectation (ICE)* [8] plots were introduced. These plots separate the aggregated line of a PDP into individual lines for each data point in the dataset, thereby revealing the specific effects for each instance. An ICE plot is essentially a collection of lines where each line represents the model’s predictions across the range of a feature for a single instance. ICE plots are valuable because they expose the diversity of responses within a dataset that PDPs might obscure. However, these plots have drawbacks, particularly as the number of instances increases. With a large dataset, ICE plots can become cluttered, making it difficult to recognise individual lines and identify significant patterns (see Figure 11).

**Fig. 11.**
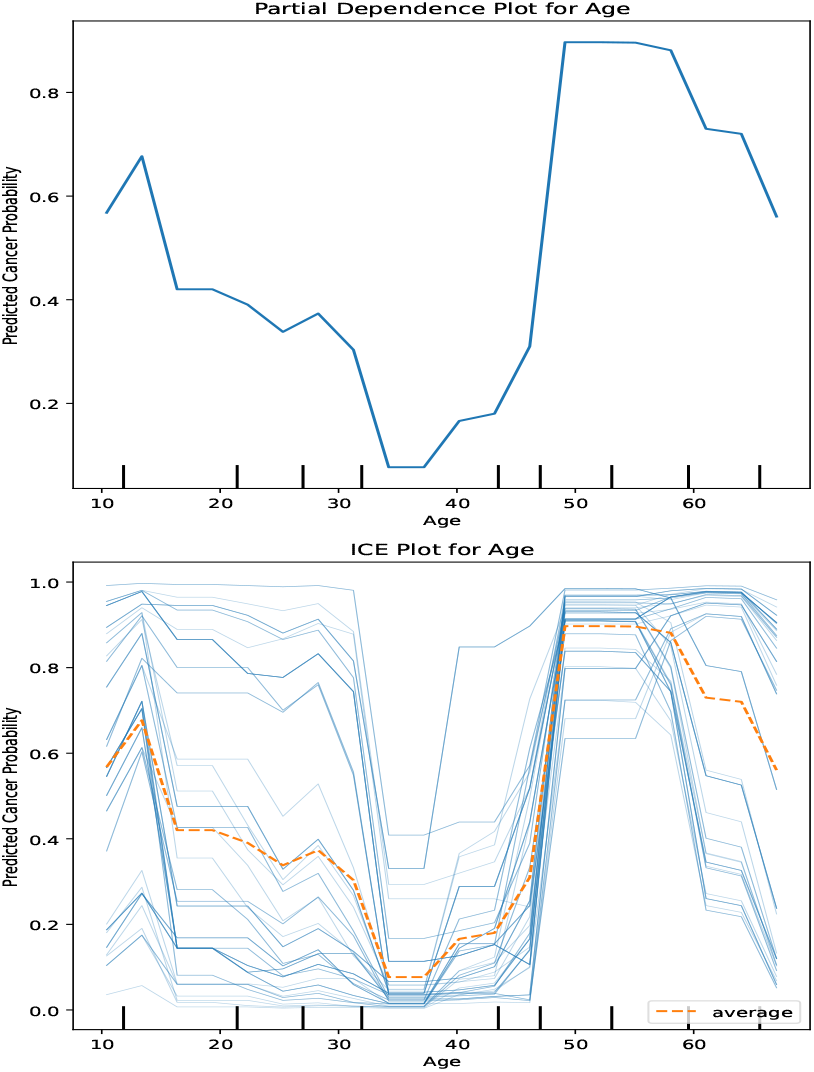
With reference to features in Figure 10, the PDP and ICE plots have been generated for feature *Age*. PDP shows the cancer probability plot for patients with age 38 vs. ICE plot shows the probability of multiple patients as age varies with other feature values as constant.

*Accumulated local effects (ALE)* [212] offer a complete analysis of the impact of individual features on the average prediction output of a machine learning model. In contrast to PDPs, ALE plots concentrate on the local changes in the prediction when a feature changes, preventing the introduction of bias in averaging the distribution of all other features. This makes ALE plots a method for feature effect analysis that is particularly robust and accurate in situations where features are correlated.

Instead of recalculating predictions across the entire data set for each point in the plot as in PDP, ALE is quicker to compute because it concentrates solely on changes within small intervals of the feature of interest. This localised computation expedites the process and improves the clarity and specificity of the insights regarding the relationship between feature values and prediction changes. Consequently, ALE plots are the optimal choice for practitioners who require a method that is both efficient and unbiased in evaluating the significance of features in intricate machine-learning models.

Friedman’s *H-statistic* [246] is used to measure the strength of interactions among features within ensemble models, assessing how these interactions contribute to the variation in predictions.

From a global perspective in model-agnostic techniques, *Feature Importance* [247] and *Multidimensional Feature Visualisation* [248] approaches explore various aspects of feature importance and relationships, potentially guiding feature engineering and model refinement procedures. The importance of a feature is assessed by observing how the accuracy of predictions changes when the relationship between that feature and the actual outcome is disrupted. Specifically, by permuting the feature, we can break the existing link between the feature and the outcome. If this permutation increases prediction error, the model relies significantly on that feature for accurate predictions.

#### A.2 Model Specific

Model-specific explanation methods are designed to work with specific model architectures and show how these models think and make decisions. These methods are beneficial for getting to know the complicated models used in healthcare, where accurate interpretations can help doctors make decisions.

One well-known example is Class Activation Maps (CAM) [214], which find essential image parts that affect how well it predicts a particular class. This method is made explicitly for convolutional neural networks (CNNs) with a global average pooling layer before their last classification layer. For example, CAMs are very helpful in medical imaging for finding areas of interest in X-rays that might show signs of disease.

Gradient-weighted Class Activation Mapping (Grad-CAM) [227] builds on CAM by using the gradients of any target idea. These gradients flow into the final convolutional layer to make a coarse localisation map that shows where the predictions are most likely to be accurate. Grad-CAM can be used with CNN designs and for more medical imaging tasks, like finding the edges of tumours on magnetic resonance imaging, drawing attention to trouble spots in dermatological images, or performing glaucoma diagnosis [249] (see Figure 12).

**Fig. 12.**
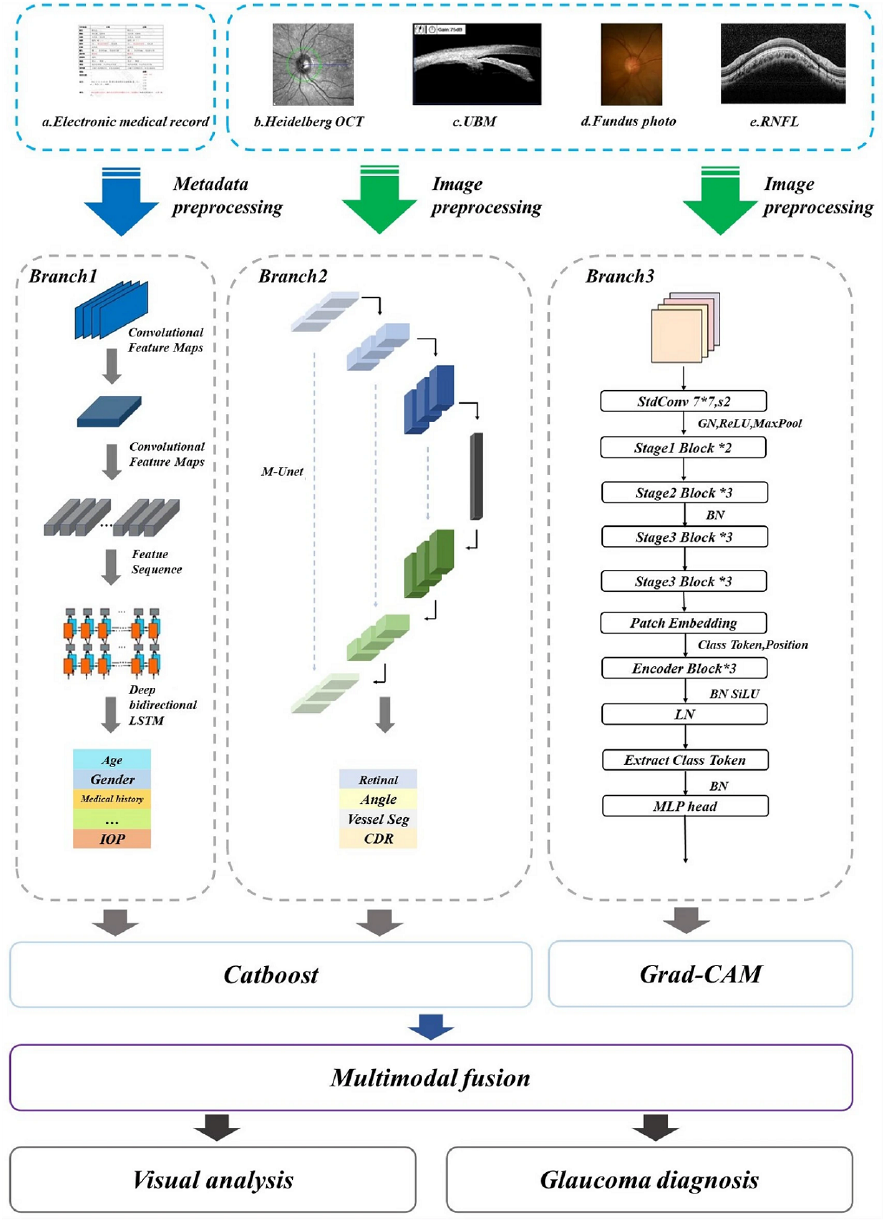
Grad-CAM based Glaucoma Multimodal Neural Network showcasing the use of Grad-CAM [249]. This image is licensed under CC BY 4.0.

Also, Deep Learning Important FeaTures (DeepLIFT) [223] and Layer-wise Relevance Propagation (LRP) [4] give us more information specific to the model. DeepLIFT looks at how active each neuron is compared to its “reference activation” and provides each neuron with a contribution number. This can be very helpful when specific biomarkers or imaging features are significant. But LRP returns the output choice to the input layer, showing which pixels or parts of an image help with a specific diagnosis. Figure 13 presents the example utilising LRP for multimodal disease prediction and explanation.

**Fig. 13.**
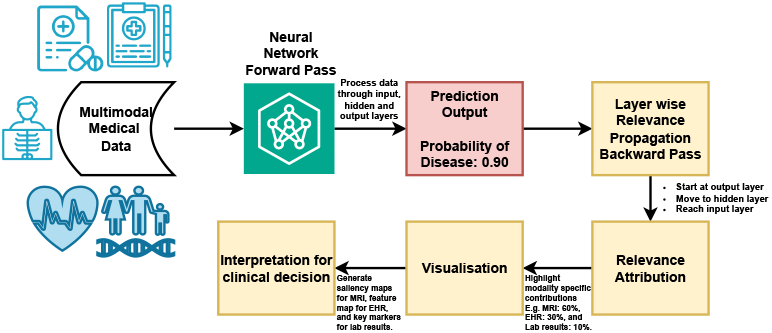
Explainable Multimodal Disease Explanation using LRP

By using these methods, doctors can better understand AI’s results and check them against what they know from clinical experience. This builds trust in automatic systems and leads to better patient outcomes.

*Class Activation Mapping (CAM)* [214] is a technique to identify regions within an image that are influential in classifying the image into a specific category. CAM is generated when the feature maps from the last convolutional layer of a CNN (noted as *F* ^*k*^ where *k* is the feature map index) are pooled using global average pooling and then connected to the output layer. This pooled output represents the average intensity of the features detected by the convolutional filters. Each class score *S*_*c*_ (prediction score for class *c*) at the output is calculated as a weighted sum of these pooled feature maps:

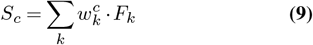

where 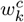 is the weight linking the *k*-th feature map to the class *c*, and *F*_*k*_ is the average pooled value of the *k*-th feature map. The Class Activation Map *M*_*c*_ for a class *c* shows the region’s importance for classifying the image. It is computed as:

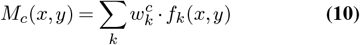

where *f*_*k*_(*x, y*) is the activation at each pixel (*x, y*) in the *k*-th feature map. By upscaling *M*_*c*_ to the size of the original image, CAM visualises the regions critical for identifying the class *c* in the image, helping to understand why the network classifies the image in a certain way.

##### Gradient-weighted Class Activation Mapping (Grad-CAM)

[227] is a technique that improves CAM by using gradient information from CNN to highlight essential areas in an image for a specific class prediction. Grad-CAM can be applied to any CNN architecture without requiring particular features like a global average pooling layer or a dense output layer. This flexibility allows it to be used widely without altering existing CNN structures. For each class *c*, Grad-CAM calculates the gradient of the class score *S*_*c*_ (which represents the likelihood of the class) for the feature map activations at the last convolutional layer. This gradient reflects how much each element of the feature map influences the class score. The weights 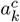 for Grad-CAM are computed by globally averaging these gradients across all spatial locations (*x, y*) of the feature map:

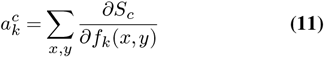

These weights measure the importance of feature map *k* in predicting class *c*.

The Class Activation Map for Grad-CAM, *Grad*_*M*_*c*_(*x, y*), is then calculated as a weighted sum of the feature maps using these derived weights, passed through a ReLU function to only keep positive influences:

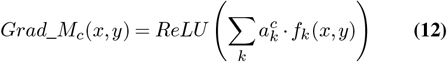

This operation effectively highlights the regions of the image most relevant to identifying the class *c*, ignoring areas with negative influence on the prediction. The resulting activation maps are scaled up to the size of the original image to show the discriminative image regions that support the CNN’s decision, providing a visual explanation of why the network predicts a particular class. This method is beneficial in applications such as medical imaging. Similarly, Grad-CAM++ [228], Smooth-Grad [237], and Smooth-GradCAM++ [238] are the variants of Grad-CAM.

*Layer-wise Relevance Propagation (LRP)* [4] is distinguished for its ability to trace back the predictions of neural networks to the contributions of individual input features across different data modalities. This method aims to distribute a prediction backwards through the network to assign relevance scores to each input feature, based on a “conservation” principle, where the sum of relevance scores is consistent across layers. The methodology is to perform a standard forward pass to obtain the output score *S*_*c*_ for a given class *c*. With the backward pass, for each layer, redistribute the relevance scores to the previous layer. The redistribution rules depend on the type of layer. There are three rules.

- LRP-*ϵ* Rule (for ReLU Layers):

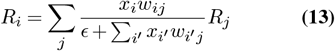

where *R*_*i*_ is the relevance of input neuron *i, x*_*i*_ is its activation, and *w*_*ij*_ is the weight connecting neuron *i* to output neuron *j*.
- LRP-*αβ* Rule (for ReLU Layers):

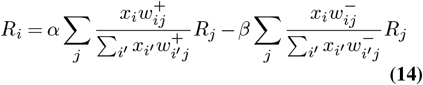

where 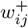 and 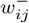 represent positive and negative weights, respectively, and *α* + *β* = 1.
- LRP-*γ* Rule: A simplified variant using a positive bias factor *γ*.

*Integrated Gradients* [239] compute feature attributions by integrating gradients over a baseline-to-input path. Select an input *x* and a baseline *x*^′^ (typically a zero or average input). Create interpolated inputs between *x* and *x*^′^:

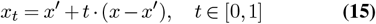

Compute the gradients of the model output *F* (*x*) concerning the interpolated inputs and average them:

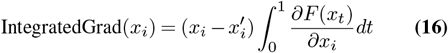

Multiply the averaged gradients by the difference between the input and baseline to obtain feature attributions:

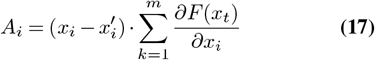

where *m* is the number of interpolation steps. The integrated gradients provide a more reliable attribution method than simple gradient methods. They also handle issues of gradient saturation by averaging over multiple steps.

*eXirt method* [171] presents an explanatory framework utilising Item Response Theory (IRT) for tree-ensemble models, providing a systematic approach to evaluating various features’ impact on a model’s decision-making process. In contrast to conventional post-hoc explainability methods like SHAP or TreeExplainer, eXirt assesses the significance of input features through a probabilistic framework based on assessment models. eXirt systematically perturbs features treated as test items to analyse their impact on model predictions, quantifying their significance through three key properties: discrimination, difficulty, and guessing. Discrimination quantifies the effectiveness of a feature in distinguishing between various diagnostic categories, while difficulty indicates the point at which a feature significantly affects model predictions. Guessing addresses the model’s reliability on a feature even after being altered.

eXirt utilises the Three-Parameter Logistic (3PL) IRT model to mathematically formalise the probability that a feature *i* contributes to a model’s prediction for a specific instance *j* as follows:

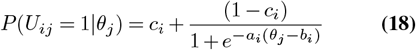

*a*_*i*_ denotes the discrimination capability of feature *i, b*_*i*_ measures its difficulty, and *c*_*i*_ indicates the likelihood of the model depending on that feature despite its minimal expected influence. The variable *θ*_*j*_ represents the estimated ability of the model in prediction, functioning as a latent measure of feature importance.

eXirt constructs a global feature importance ranking by aggregating individual feature perturbations across multiple instances to derive an overall interpretability score. The final importance score *T* (*f, r*) is calculated as follows:

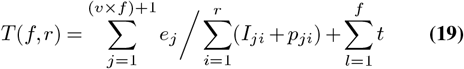

*v* represents the number of perturbations applied to a specific feature, *f* denotes the total number of input features, and *I*_*ji*_ indicates the estimated parameters derived from the response matrix. The notation *p*_*ji*_ denotes the ability estimate of the relevant feature, whereas *t* represents the total score obtained from classification results, defined as:

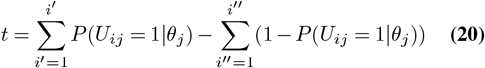

*i*^′^ denotes instances that are correctly classified, while *i*^′′^ indicates instances that are misclassified. This scoring mechanism guarantees that features with substantial influence on predictions are assigned a higher interpretability ranking, whereas those with negligible impact are ranked lower.

The advantage of eXirt is particularly evident in multimodal medical data, where features originate from diverse sources. In high-dimensional datasets, conventional feature attribution methods may inadequately represent the intricate interactions among various modalities. For instance, in the context of a diagnostic model for *Alzheimer’s disease*, eXirt could indicate that hippocampal atrophy (by MRI), genetic risk factors like the presence of the APOE4 allele, and cognitive assessment scores would collectively influence the classification outcomes of the model. eXirt could enhance interpretability by quantifying the role of each feature through a structured, response-driven method, thereby providing essential reliability for high-stakes decision-making, including clinical diagnoses.

In contrast to other model-specific explanation methods, eXirt incorporates a probabilistic perspective into explainability, aligning feature importance with established psychometric evaluation principles. This provides a structured and theoretically informed method for feature ranking, allowing healthcare practitioners to comprehend the rationale behind prioritising specific inputs in a prediction. eXirt enhances trust in AI-driven decision-support systems by integrating explainability with reliability, enabling clinicians and medical researchers to make informed and accountable interpretations of model predictions.

#### A.3 Intrinsic Model

Intrinsic models have an inherent interpretability, which is crucial when dealing with complex multimodal medical data, including clinical notes, imaging, and genetic information. Models like linear models, logistic regression, decision trees, and explainable boosting machines each provide varying degrees of interpretability and detail, making them suitable for medical applications. Linear models are fundamental, assuming a linear relationship between the input variables and the outcome. For example, a linear regression could predict patient recovery time based on a combination of numerical data, such as age and white blood cell count, and coded categorical data, such as multiple medical conditions [250, 251]. In a linear regression model used to predict heart disease risk using multimodal medical data, the equation might look like this:

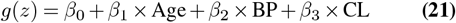

*g*(*z*) represents the predicted risk of heart disease, *β*_0_ is the baseline risk when all factors are zero, and *β*_1_, *β*_2_, and *β*_3_ are coefficients that indicate how much each factor - age, blood pressure (BP), and cholesterol level (CL) affects the risk. Each coefficient reflects the expected increase in disease risk associated with a one-unit increase in its corresponding feature, helping medical professionals understand the impact of these variables on heart health.

Logistic regression is an extension of linear regression used for classification problems, such as determining whether a disease is present or absent. It predicts the probability of a binary outcome by using the formula:

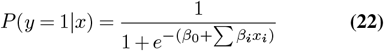

This model incorporates various types of data, such as imaging features and genetic markers, where *β*_0_ is the baseline effect when all other variables are zero, and *β*_*i*_ are the co-efficients indicating how significantly each variable *x*_*i*_ influences the likelihood of the disease. This setup provides a clear understanding of how different factors contribute to the probability of a disease occurring.

Decision trees efficiently manage multimodal data by categorising patients using the GINI index, a measure of data purity at each node. The equation for the GINI index is:

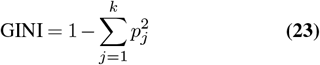

where *p*_*j*_ represents the proportion of each class *j* within a node and *k* is the number of classes. For instance, in predicting heart disease, a decision tree might evaluate splits like patient age, using the GINI index to find the most effective division. If splitting by age, one group could be under 45 years old with a GINI of 1 − (0.7^2^ + 0.3^2^) = 0.42 and another 45 or older with a GINI of 1 − (0.4^2^ + 0.6^2^) = 0.48. The tree selects the feature and threshold that minimize the GINI, continually refining the data split at each node until it meets specific criteria like maximum depth or minimum node size, ensuring a clear and statistically robust pathway from patient data to diagnosis.

Explainable Boosting Machines (EBM) is a predictive tool that integrates the advantages of boosting methods in machine learning with the transparency and interpretability of decision trees and generalised additive models [181]. Typically, EBMs utilize a sequence of compact decision trees, each tailored to a distinct data category. For instance, one tree is designated for imaging features, another for clinical parameters, and another for genetic markers. By employing this methodically organised strategy, EBMs can preserve a significant degree of interpretability, given that every tree concentrates on elucidating precisely how its specific collection of features influences the overall prognosis. Furthermore, EBMs optimize performance by focusing on particular characteristics within each data domain. This guarantees that the model not only generates precise predictions but also provides valuable insights into the correlation between different categories of data and the anticipated results. EBMs are especially advantageous in intricate settings such as healthcare, where comprehending the impact of various data streams is vital.

The transparency of these models builds trust among medical practitioners and patients by clearly illustrating the reasoning behind clinical decisions, which is particularly important when decisions have significant health outcomes.

#### A.4 Usage, Limitations, and Complexity

Comprehending the assumptions and trade-offs of explainable AI (XAI) methodologies is crucial for their effective use in clinical environments. Techniques such as LIME, SHAP, Grad-CAM, and CAM are extensively employed, each possessing unique advantages and drawbacks, as outlined in Table 6. These methods are customised for distinct data types and applications, exhibiting differing computational expense and interpretability degrees.

**Table 6.** XAI Methods: Usage, Limitations, Computational Cost, and Datatypes

| Method | Typical Clinical Use | Key Limitations/Assumptions | Computational Cost | Datatype |
| --- | --- | --- | --- | --- |
| **LIME** [229] | Explaining individual diagnoses (e.g., sepsis prediction from lab parameters) | **Limitations:** May require dense sampling, leading to high computational costs; sampling introduces uncertainty, potentially resulting in variable explanations for the same input.<br>**Assumptions:** Assumes local linearity and feature independence during perturbations, which may not hold in complex medical data. | Moderate; requires repetitive perturbations around the instance | Structured EHRs |
| **SHAP** [6] | Local and global feature attribution (e.g., stroke prediction integrating demographic and imaging data) | **Limitations:** Computationally expensive for large feature sets; KernelSHAP is particularly costly in high dimensions.<br>**Assumptions:** Assumes feature independence, which may not hold in medical data. | Potentially high; TreeSHAP/DeepSHAP versions mitigate cost for tree/DNN models | Structured/Unstructured EHRs, Image Data |
| **eXirt** [171] | Global and local explainability in tree-ensemble models (e.g., ranking feature importance in multimodal disease risk prediction) | **Limitations:** Designed specifically for tree-ensemble models; may not generalize well to other architectures.<br>**Assumptions:** Assumes perturbation-based response behavior follows IRT principles; feature dependencies may influence interpretability. | Moderate to high; requires iterative feature perturbations and response matrix computations | Structured EHRs, Omics, Imaging Biomarkers |
| **Grad-CAM** [227] | Localizing lesions in medical images (e.g., tumor boundaries on MRIs) | **Limitations:** Focuses on the last convolutional layer, leading to coarse heatmaps that may miss fine-grained details.<br>**Assumptions:** Assumes the most relevant features are in the final convolutional layer. | Relatively low; one forward + one backward pass for the target class | Image Data |
| **CAM** [214] | Highlighting discriminative regions in CNN-based classification (e.g., suspicious tissues in histopathology) | **Limitations:** Requires global average pooling before the final layer; limited to specific architectures.<br>**Assumptions:** Assumes the discriminative power is captured in a linear combination of feature maps. | Low to moderate; depends on CNN architecture constraints | Image Data |
| **Integrated Gradients** [240] | Pixel-level or feature-level attribution in structured data (e.g., tabular data on lab results) and medical images | **Limitations:** Choice of baseline significantly impacts results; gradient saturation can still be an issue.<br>**Assumptions:** Assumes interpolation along a straight path between baseline and input provides meaningful attributions. | Moderate to high; requires multiple forward/backward passes for interpolation steps | Structured EHRs, Image Data |
| **DeepLIFT** [223] | Feature attribution in DNNs (e.g., analyzing which genes or lab values cause a high-risk prediction) | **Limitations:** Baseline selection can introduce bias; may be less interpretable for non-standard activation layers.<br>**Assumptions:** Assumes differences in neuron activations relative to a baseline provide meaningful explanations. | Similar to integrated gradients, it does multiple passes through the network; overall moderate | Omics, Structured EHRs |
| **Layer-wise Relevance Propagation (LRP)** [4] | Decomposes CNN or DNN predictions back to input features (e.g., highlighting critical pixels for skin lesion classification) | **Limitations:** Highly sensitive to the redistribution rule used for relevance propagation.<br>**Assumptions:** Assumes relevance can be decomposed layer by layer in a meaningful way. | Moderate; one backward pass per prediction for relevance propagation | Image Data |
| **Anchor Explanations** [252] | Local rule-based explanation for classification (e.g., describing minimal conditions that firmly fix a diagnosis outcome) | **Limitations:** May require dense sampling, leading to high computational costs; sampling introduces uncertainty, potentially resulting in variable explanations for the same input.<br>**Assumptions:** Assumes high-precision rules can anchor predictions effectively. | Moderate; relies on sampling-based coverage checks for rule anchors | Structured EHRs |
| **Contrastive Explanation Method (CEM)** [218] | Local explanations focusing on Pertinent Positives/Negatives (e.g., minimal needed features for diagnosing pneumonia vs. minimal absent features) | **Limitations:** Computationally expensive due to iterative optimization; struggles with high-dimensional data.<br>**Assumptions:** Assumes minimal perturbations that flip predictions provide meaningful contrastive explanations. | High; requires iterative optimization to identify positives/negatives | Structured EHRs, Omics |
| **ProtoPNet** [253] | Example-based reasoning in CNNs (e.g., linking new chest X-rays to "prototype" X-rays learned during training) | **Limitations:** Performance depends on prototype diversity and interpretability; tuning prototype dimensions is challenging.<br>**Assumptions:** Assumes learned prototypes effectively represent real-world data clusters. | Moderate to high; prototypes require memory/space, but explanation is more intuitive (case-based) | Image Data |
| **Case-Based Reasoning (CBR)** [233] | Uses a "library" of medical cases to explain new predictions via similar past examples (e.g., referencing previous patient files) | **Limitations:** The case library must be large and diverse to ensure meaningful explanations; similarity metrics can be misleading if not carefully chosen.<br>**Assumptions:** Assumes past cases provide relevant and generalizable explanations for new cases. | Varies; typically moderate; case retrieval cost depends on library size and indexing strategy | Structured/Unstructured EHRs |
| **Counterfactual Explanations** [225, 226] | Clarifies what minimal changes would alter the diagnosis (e.g., slight changes in lab values to reduce the risk class) | **Limitations:** Finding actionable and plausible counterfactuals is challenging; correlated features can make generation unreliable.<br>**Assumptions:** Assumes minimal feature modifications provide meaningful insight into model decisions. | High in naive search-based approaches; advanced methods (GAN-based) can mitigate cost but add complexity | Structured EHRs, Behavioral Data |

LIME and SHAP are effective methods for interpreting predictions, particularly in tabular datasets and structured electronic health records (EHRs). Both methods depend on perturbing or sampling data points, which may result in mistakes within strongly correlated feature spaces, characterised by overlapping laboratory markers or co-occurring symptoms. SHAP is sometimes seen as more theoretically robust. However, it is typically computationally demanding, especially with extensive datasets or intricate models. eXirt offers a systematic method for explainability by incorporating Item Response Theory (IRT) principles to evaluate feature significance in tree-ensemble models. eXirt systematically perturbs input features, treated as test items, to generate a response matrix that quantifies their influence on model decision-making. This methodology is especially pertinent for multimodal medical datasets, where the interconnections among structured electronic health records, genomic data, and imaging biomarkers are essential for diagnostic modelling. In contrast to LIME or SHAP, which assess individual feature contributions in a model-agnostic way, eXirt produces probabilistic feature rankings informed by difficulty, discrimination, and guessing parameters, rendering it particularly effective for high-dimensional and heterogeneous medical datasets. Nonetheless, its dependence on perturbations and response modelling may lead to sensitivity regarding feature dependencies and necessitate moderate to high computational expenses.

Conversely, Grad-CAM and CAM are particularly effective for image-centric applications, including lesion localisation in MRI scans and identifying questionable areas in histopathological images. Grad-CAM offers imprecise explanations centred on the last convolutional layer, whereas CAM necessitates architectural attributes such as global average pooling, potentially restricting its applicability to specific neural networks. Alternative strategies tackle various faces of interpretability. Integrated Gradients and DeepLIFT provide pixel- or feature-level attributions and can process structured data, medical imaging, or omics datasets. Nevertheless, their reliance on baseline selection may introduce biases, and their computational expense might escalate for models necessitating multiple interpolation stages. Layer-wise Relevance Propagation (LRP), intended for convolutional neural networks (CNNs) and deep neural networks (DNNs), disaggregates predictions to input features; even so, it is sensitive to redistribution rules that necessitate careful adjustment.

Rule-based methodologies such as Anchor Explanations and case-based reasoning prioritise simplicity and intuitiveness, frequently demonstrating efficacy with structured electronic health records (EHRs). Anchor Explanations employ minimal prerequisites to elucidate an outcome effectively but encounter difficulties with continuous or highly connected variables. Case-based reasoning depends on a repository of comparable examples for forecasting and explanation, portraying the quality of its outputs as significantly reliant on the diversity of the repository and the similarity metrics utilised.

Approaches such as ProtoPNet and counterfactual explanations offer distinct interpretive frameworks. ProtoPNet links predictions with acquired prototypes, providing a more comprehensible reasoning framework, particularly for image data. Counterfactual methodologies elucidate the smallest modifications to input variables that could modify a diagnosis, proving especially beneficial for behavioural data or structured electronic health records (EHRs). Generating actionable counterfactuals is computationally intensive and necessitates accurate management of feature correlations and domain limitations.

According to Table 6, the appropriate XAI approach depends upon the particular clinical application, the nature of the data under analysis, and the available computational resources. By comprehending these details, researchers and clinicians can more effectively utilise XAI techniques to enhance transparency and confidence in AI-driven healthcare systems.

### B Visual Approach

Visual explanation techniques improve the interpretability of complex AI models, especially in medical applications. By visually emphasising essential input features, clinicians can naturally and efficiently understand AI predictions, thus improving trust and providing informed decisionmaking in critical circumstances. Significant methods involve attribution-based explanations that assess the influence of particular input features on model outputs. This is achieved through techniques including saliency based explanations, perturbation based explanations, concept-based explanations, decomposition methods, and contrastive methods as shown in Figure 14. Example-based explanations provide a distinct perspective by utilising prototypes, counterfactual explanations, and case-based reasoning to contextualize model reasoning. Interactive tools like Intel’s Explainable AI Toolkit, Microsoft’s InterpretML, and Google’s What-If Tool enable users to engage with models, facilitating the analysis of feature importance and the observation of prediction changes in response to input variations. These techniques collectively establish a comprehensive framework for enhancing explainable artificial intelligence (XAI) in healthcare. They facilitate the integration of various data types, such as imaging, clinical parameters, and genomic information, to tackle the specific challenges associated with multi-modal medical data.

**Fig. 14.**
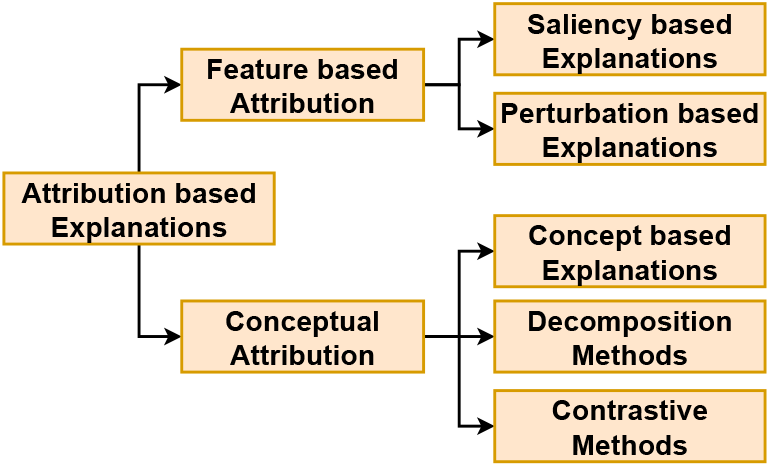
Classification of Attribution-based Explanation Methods based on level of abstaction

#### B.1 Attribution-based Explanations

Based on the level of abstraction, the attribution methods can be classified into two categories - Feature-based attribution (low level) and Conceptual attribution (high level).

Attribution-based methods aim to quantitatively evaluate the contribution of individual input features to a model’s predictions. These techniques are particularly important in medical applications, as they help identify the most critical elements of the data, thereby enhancing trust in the model’s outputs and supporting clinical decision-making. In multimodal settings, attribution methods are instrumental in analysing the impact of diverse data sources and their interactions on predictions. Broadly, feature attribution methods can be categorised into two primary types: saliency-based methods and perturbation-based methods. Additionally, conceptual attribution approaches can be divided into three categories: concept-based explanations, decomposition methods, and contrastive methods.

##### Saliency-based Explanations

Saliency map-based methods are famous for XAI in computer vision tasks, as they can highlight regions of the input data that affect the model’s output most strongly [234]. It is referred to as explanations through heatmap, sensitivity map, feature importance map or feature attribution map in literature [254–257]. These methods are extremely valuable in multimodal medical data - such as combining the clinical metadata with the medical imaging (e.g. CT, PET, MRI, or X-ray), as these methods can bring the clinician’s attention to the most valuable features or anatomical structures relevant for the diagnostic decision. By making the reasoning patterns of an underlying model more transparent, saliency maps help reduce the black-box concerns to a great extent [258].

From the methodology-based standpoint, backpropagation is what saliency maps rely on. Early models measure each pixel’s importance using Vanilla Gradient extractions with regard to the input pixels [215]. Later improvements change the backward pass to produce finer, more interpretable gradient maps using Guided Backpropagation [259] and the DeConvNet method [260]. Grad-CAM and Grad-CAM++ [227, 228] include class-specific gradient signals in convolutional feature maps, therefore allowing coarse localisation of relevant regions in 2D or 3D medical scans. Modern approaches, such as Integrated Gradients [239], solve gradient saturation concerns. By averaging across several noisy inputs, techniques such as SmoothGrad [237] and Smooth-GradCAM++ [238] lower noise in saliency maps; Layer-wise significance Propagation (LRP) [4] methodically assigns significance to every pixel or voxel by propagating predictions backwards through the network layers.

Saliency maps can highlight the essential words and phrases in textual data such as electronic health records (EHRs). For a trained classification model, phrases such as “shortness of breath” or “chest pain” in clinical notes, when highlighted by the saliency-based XAI model, give more confidence to the medical decisions of the experts [261, 262]. Saliency maps help experts understand patients’ medical history in less time, which can be crucial for doctors to serve more patients per day. In the case of time-series data such as Electrocardiograms (ECGs) and Electroencephalograms (EEGs), the saliency maps, can highlight specific waveform features and time stamps, which helps medical experts save time when diagnosing a heart condition or a brain seizure [263, 264]. In models integrating multimodal data, such as imaging, EHR data, and lab test results, saliency maps can be applied individually on each data type and presented to the medical expert for their decision to be aided [265]. Applied across MRI modalities including T1, T1C, T2, and FLAIR, Figure 15 shows the use of saliency maps produced using guided back-propagation and Shapley Value Sampling. Modality-Specific Feature Importance (MSFI) scores were derived from these maps and compared with doctor ratings.

**Fig. 15.**
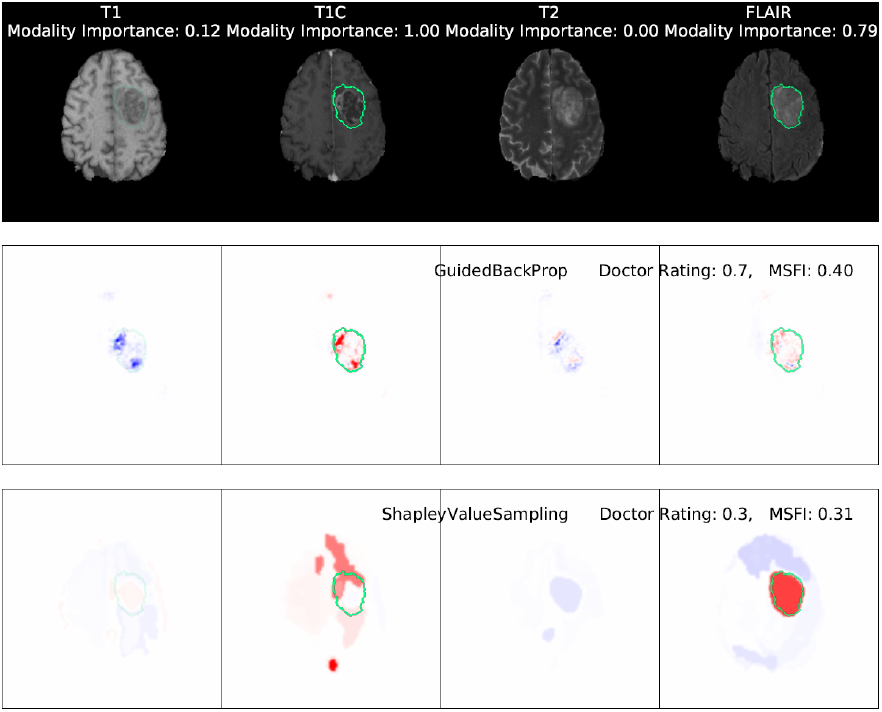
MRI images of a high-grade glioma, alongside saliency maps generated by two methods: Guided Backpropagation and Shapley Value Sampling [266]. This Figure is licensed under CC-BY-NC-ND.

##### Perturbation-based Methods

Perturbation-based methods aims at changing the input feature in such as way that it affects the model output. The most significant perturbation-based approaches include LIME [229], SHAP [6], KernelSHAP [224], Occlusion [260], meaningful perturbations [267], real-time perturbations [268], extremal perturbations [269], randomised input sampling for explanation (RISE) [270], and hierarchical perturbations (HiPe) [271]. Malhi et al. [272] used perturbation approaches to explain the prediction of medical diagnosis algorithms, namely the LIME method, to explain a classifier’s decisions to detect bleeding in gastral endoscopic images. Similarly, Punn et al. [273] employed the LIME technique to explain the predictions of multiple cutting-edge deep learning models that categorize pulmonary illnesses in chest X-ray images. Magesh et al. [274] employed LIME to justify the choices of a Parkinson’s disease classifier model. For melanoma recognition, Young et al. [275] used the Kernel SHAP [6] interpretability approach to test its reliability in explaining a melanoma classifier from dermoscopy images. They concluded that the interpretability technique emphasised unimportant elements to the final prediction. The authors also ran sanity checks on the interpretability approaches, confirming that they frequently offer alternative interpretations for models with equal performance. This can be explained by the model’s ability to learn some spurious associations, causing interpretability approaches to overestimate the relevance of spurious regions emphasised on the resulting saliency maps. In the same context, Wang et al. [276] suggested a multimodal CNN for skin lesion identification that took into account both patient meta-data and skin lesion images. They used SHAP to examine each feature’s contribution to patient metadata. Similarly, Eitel et al. [277] used an occlusion-based interpretability method [260] and tested its robustness for Alzheimer’s disease categorisation. GeneXAI framework [278] identified top genes in stages I and II using LIME and SHAP utilising early fusion for extracting the multimodal features.

##### Concept based Explanations

Concept-based explanation methods aim to establish a connection between human and machine perception by connecting model decisions with concepts that are more abstract and understandable to humans [279]. They help decode machine learning (ML) models that analyse multimodal medical data [280], which may include formats such as EHRs, behavioural data, omics data, sensors data, or images. Using concept-based explanations, we can associate model decisions with clinically significant concepts across diverse data types, providing a clear understanding of diagnoses and prognoses.

Concept-based explainability methods provide a structured framework for this interpretation. At the neuron level, methods like Network Dissection reveal how specific neurons encode features such as anatomical structures or molecular signatures. Layer-level methods, including Concept Activation Vectors (CAVs), extend this understanding by identifying how groups of neurons collectively represent higher-level concepts like disease patterns or pathological markers. Concept Bottleneck Models (CBMs) [281] further refine this by explicitly mapping these concepts to dedicated neurons, offering direct insights into a model’s decision-making process. Finally, probing methods evaluate the extent to which layers or neurons capture nuanced relationships, such as multimodal interactions between imaging data and clinical variables. Together, these approaches offer granular and holistic interpretability that is invaluable for medical diagnostics.

Table 7 summarises these methods into categories, types, and examples to provide a comprehensive overview of the diverse methodologies in concept-based explanations. It systematically organises the types of concepts (e.g., symbolic, unsupervised, prototypes, and multimodal) and explanations (e.g., class-concept relations, node-concept associations). A brief description and example methods or techniques are presented in each type. This categorisation highlights the broad applicability of concept-based methods across domains, from post-hoc techniques like TCAV [241] and CRAFT [282] to explainable-by-design models such as CBMs [281] and ProtoPNet [253]. For instance, symbolic concepts highlight human-aligned interpretations of features like colour or shape [241], while temporal concepts explore time-sequenced data like motion patterns in video analysis [283]. Similarly, multimodal concepts bridge data types, such as linking imaging and text, to enhance semantic understanding [213, 284]. By mapping these methodologies to their respective examples, the table provides an essential reference for navigating the field of concept-based AI explainability.

**Table 7.** Concept and Explanation Types in Concept-based Explanations

| Category | Type | Description | Examples/ Methods |
| --- | --- | --- | --- |
| Concept Types | Symbolic Concepts | Human-defined, interpretable features such as colour, shape, or material. These concepts are directly aligned with human understanding. | TCAV [241], CAR [286], Network Dissection [287] |
|  | Unsupervised Concept Bases | Automatically discovered clusters of features in a model’s latent space. Requires no manual annotations. | ACE [288], ICE [289], CRAFT [282] |
|  | Prototypes | Specific instances that represent a concept or class. Common in case-based reasoning to identify critical features or examples in data. | ProtoPNet [253], ProtoTree [290], ProtoPShare [291] |
|  | Temporal Concepts | Concepts derived from time-sequenced data, such as motion patterns or trajectories in video models. | Temporal TCAV [283], Spatial-Temporal Concept Analysis [292] |
|  | Multimodal Concepts | Concepts represented across multiple modalities, such as vision-language models, to capture richer semantic relationships. | Concept Transformers [284], DIME [213] |
|  | Textual Concepts | Concepts represented by embeddings derived from textual data, enabling alignment with natural language. | Label-free CBM [281], Vision-Language Models [293, 294] |
| Explanation Types | Class-Concept Relation | Describes the influence of a concept on a specific class decision, aiding in model debugging and feature attribution. | TCAV [241], CaCE [295], Concept Importance Estimation [289] |
|  | Node-Concept Association | Associates individual neurons or filters with specific concepts to trace internal representation alignment. | Net2Vec [296], Concept Neurons [297], Guided Concept Projection Vectors [298] |
|  | Concept Visualisation | Visualises how a model represents and uses concepts in its decision-making process, often highlighting interpretable regions in input data. | ProtoConcept [299], CRAFT [289], Activation Maps [282] |
|  | Ontological Commitment | Captures relationships between concepts (e.g., hierarchical or logical) to verify consistency with human cognition or predefined rules. | Concept Hierarchies [300], Region-Based Representations [301, 302] |
|  | Temporal Dynamics | Explains how concepts evolve over time, critical in dynamic or sequential decision-making systems like video models. | Optical Flow-Based Concept Extraction [283, 303] |

A model may analyse imaging data and genetic information in the context of cancer diagnosis. By combining particular genetic markers with specific textural features in a tumour image [285], concept-based explanations may recognise how this information influences the classification of cancer type by the model. In this context, concepts such as genetic mutation types derived from genomic sequences or tumour texture extracted from MRI scans are crucial for rendering the model’s reasoning comprehensible to clinicians. By organising such methods into categories, Table 7 guides researchers and practitioners to identify suitable tools for specific interpretability challenges. In treating chronic diseases such as diabetes, models may analyse patient medical records, laboratory findings, and continuous glucose monitoring information. In this context, concepts such as past treatments’ responses to glucose level trends or historical glucose level trends could be utilised to comprehend and forecast patientspecific reactions to various treatment strategies.

Using concept-based explanations in healthcare contexts clarifies the model’s decision-making process and augments the confidence and practicality of AI applications. By enabling healthcare professionals to cross-check the AI insights with established clinical knowledge and practical experience, they ensure the scientific validity and clinical relevance of the model’s predictions.

##### Decomposition Methods

Unlike LIME and SHAP, which focus directly on input features, Layerwise Relevance Propagation (LRP) begins at the network’s output layer. It redistributes the output prediction via the network layers until it reaches the input layer. This redistribution is based on each neuron’s contribution to the next layer’s activity, which effectively decomposes the output into contributions from all neurons in the network. This procedure shows the paths via which significant neurons (and, eventually, input features) influence the output, resulting in a precise map of how input features are processed across the network to contribute to the final choice. DeepLIFT [304] decomposes the output predictions back through the network by comparing the influence of each neuron activation to a reference activation. DeepLIFT, like LRP, allows us to understand the contribution of each input feature across layers of a deep neural network, making it extremely useful in the context of deconstruction. Integrated Gradients [240] correlate the network’s output prediction to its inputs by integrating the gradients of the output to the inputs along a path from a baseline to the input. This technique thoroughly dissects input contributions, similar to how LRP tracks input influence through the network.

##### Contrastive Methods

Contrastive Explanation Methods (CEM) [218] include the concept of what should be minimally and adequately present to justify its classification (Pertinent Positives), and what should be minimally and necessarily absent (Pertinent Negatives). For instance, to validate a high-risk stroke classification based on a male patient aged 60 with hypertension and significant arterial plaque, pertinent positives such as these must be present due to their strong correlation with stroke risk. Conversely, the absence of pertinent negatives like protective genetic factors and adequate blood pressure management supports this high-risk assessment. By contrasting this profile with a similar demographic but classified as low risk where negative factors might be absent and positive factors less pronounced, clinicians can better understand and explain why this patient is categorised as high risk. This systematic analysis aids clinicians in discussing preventive measures and treatments with patients by highlighting the underlying reasons for the model’s predictions.

White et al. 2023 [305] proposed CLEAR *Image*^9^ framework, which utilised generative adversarial network (GAN) to generate a synthetic image which acts as a contrast image in image classification. CLEAR’s contrastive, counterfactual, and measurable explanations outperformed Grad-CAM and LIME. Using pointing games, it presented novel image explanation, segmentation, and evaluation methods. Hager et al. 2023 [306] presented a self-supervised contrastive learning framework which worked on the tabular and image data for training unimodal encoders. They combined SimCLR [210] and SCARF [307], two leading contrastive learning techniques. The dataset contained MRI images and 120 feature tabular data from 40000 UK Biobank subjects [308]. They used attribution and ablation-based experimentation to discover tabular features, which contributed to understanding shape and size, improving the quality of learned features.

#### B.2 Example based Explanations

Methods that describe a model’s decisions through a subset of related instances are called example-based explanation methods. In addition to clarifying algorithmic decisions, clinicians frequently utilize these techniques to justify the reasoning that informs their decision-making. Classification of example-based explanation methods includes (i) prototypes, (ii) counterfactual explanations, and (iii) case-based reasoning.

##### Prototypes

The prototypes-based explanation approach implemented during model training entails acquiring knowledge from a collection of “prototypes” that serve as visual aids to depict distinct categories or classes (e.g., denoted by the colours blue and orange in Figure 16). These prototypes exemplify each class in an ideal manner. When a new image is evaluated during the testing phase, the system extracts and compares its features to those of the learned prototypes. It employs a technique similar to cosine similarity to determine the degree to which the characteristics of the new image resemble those of each prototype. Subsequently, the ultimate classification prediction for the test image is ascertained using scores similar to the above. The system essentially predicts the class of the new image through a comparison with the best-matching examples learned during the training process.

**Fig. 16.**
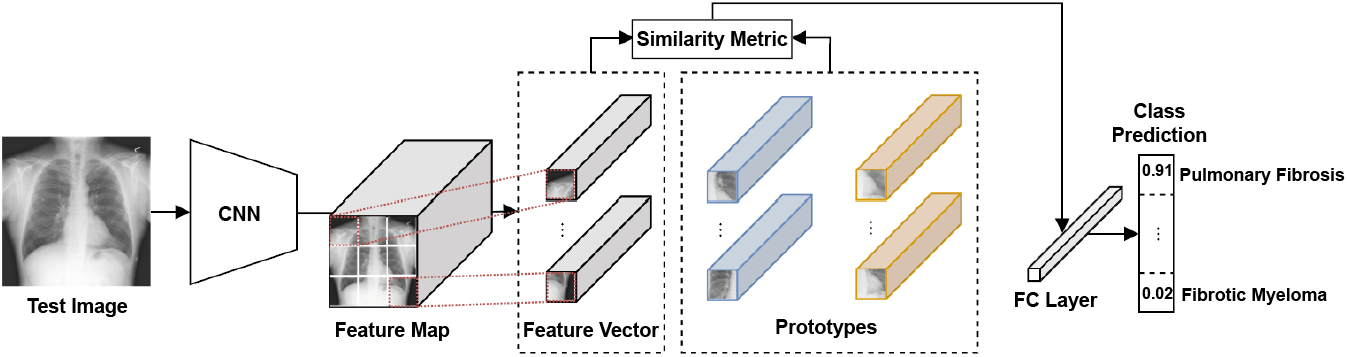
Prototype Explanation using CNN for Chest X-ray Test Image. Adapted from [21]. This image is licensed under CC BY 4.0.

Yildirim et al. 2024 [309] integrated large language models with vision encoders in radiology, can improve multimodal medical explainable AI (XAI), particularly within prototype-based methods. This work enables the creation of interpretable prototypes that represent typical or atypical medical cases, by utilising vision-language models (VLMs) for tasks such as generating radiology findings and answering visual queries. Clinicians were involved in the iterative, multidisciplinary design process, which guarantees that these prototypes are technically accurate and clinically pertinent. This process increased their acceptance and utility in medical practice. The potential for the expansion of prototype-based XAI across healthcare is underscored by these collaborative endeavors, which provide a comprehensive framework for developing interpretable and trustworthy AI tools.

Cheng et al. 2023 [310] introduced a novel prototype representation learning framework emphasising the granularity of multimodal interactions by concentrating on global and local alignments between medical images and reports. Their framework incorporated a local alignment module for more precise representation and a cross-modality conditional reconstruction module to facilitate the exchange of information between modalities during the training phase, in contrast to conventional methods that primarily rely on global multimodality alignment. This was accomplished by reconstructing obscured images and reports. A sentence-wise prototype memory bank is implemented for lengthy reports, increasing the network’s emphasis on critical clinical linguistic aspects and detailed visual features. Furthermore, the framework introduced a generation paradigm that is non-auto-regressive and is designed for the reconstruction of non-sequential reports.

##### Counterfactual Explanations

Counterfactual XAI for multi-modal data need the utilisation of a black-box model *f* (·), a diverse dataset *x* consisting of different types of data such as images and text, and a specific target class *y*_*t*_. This approach utilises a concept library *C* = {*c*_1_, …, *c*_*N*_} consisting of *N* keywords. It generates a vector *w* that represents the relevance scores for each idea. The score *w*_*i*_ represents the degree to which the concept *c*_*i*_ needs to be adjusted to align the model’s prediction with *y*_*t*_. A perturbation function *p*(·) takes as input the model *f*, dataset *x*, concept library *C*, and scores *w*, and generates a modified prediction *y*_*p*_. The objective is to identify the most favourable scores *w*^*^ that reduce the disparity between *y*_*t*_ and *y*_*p*_, by employing cross-entropy loss. This approach is valuable in multimodal situations for demonstrating the impact of diverse data kinds on predictions. It aids in identifying the required modifications for successful classification and understanding the factors contributing to precise predictions for a selected *y*_*t*_ [311, 312].

For instance, a machine learning model has been taught to identify diseases using various types of medical data, such as X-ray images and textual medical observations. A potential counterfactual explanation for this model could entail modifying the input data, such as adjusting specific clinical factors or pixel values in an image, to change the model’s diagnosis from “pneumonia” to “no pneumonia.” These tweaks aid in pinpointing the slightest alterations required to alter the forecast, providing doctors with concrete insights into the decision-making criteria of the model.

Figure 17 shows a generative model for medical diagnosis counterfactual explanations. Multimodal data, including a test image and written report, are sent into an encoder, compressing them into a latent space representation (*z*). This representation uses a decoder to reconstruct a counterfactual example using image and text data. This example shows how little modifications modify the model’s prediction. This counterfactual output helps the clinicians understand the scenario and explain the reasoning to the patient.

**Fig. 17.**
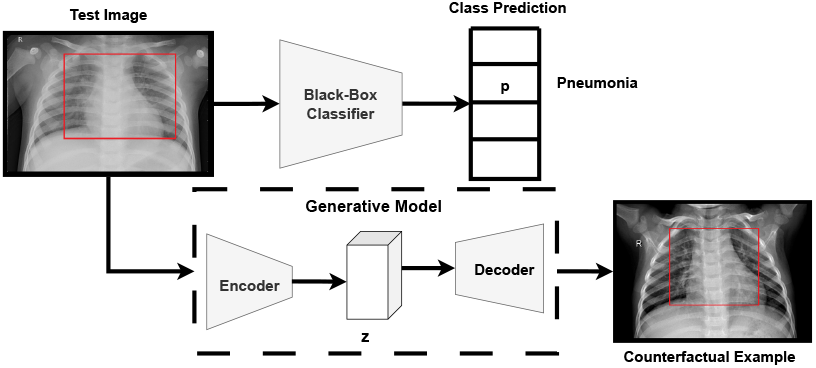
Counterfactual Explanations over Multimodal Data. Adapted from [21]. This image is licensed under CC BY 4.0.

One of the early model-agnostic counterfactual techniques for Explainable AI (XAI) is WatcherCF [313], which is particularly noteworthy. Another method, Feasible and Actionable Counterfactual Explanations (FACE) [226], produces comprehensible counterfactual explanations by calculating the shortest path distances weighted by density. Diverse Counterfactual Explanations (DiCE) [225] generate a variety of distinct counterfactual explanations for a given occurrence *x*, enabling users to select explanations that are easier to understand. CF-GNNExplainer [219] is designed to work with graph data. It uses sparsification methods to remove edges from the original adjacency matrix repeatedly. This process allows it to observe changes in prediction and determine the slightest perturbation needed based on edge reduction.

Zhou et al. [235] worked using Sparse CounterGAN (SCGAN) to create counterfactuals using modified MR images of breast cancer with desired traits or outcomes. Radiomic characteristics from MR images, alongwith tabular information from other patients, were used to train a generator-discriminator GAN. The generator generated masked residuals to construct a realistic yet counterfactual instance from the input data. The discriminator and classifier ensure that these counterfactuals are realistic and represent a distinct class outcome, such as a non-tumour condition. Oh et al. [314] proposed a framework that integrated counterfactual explanations in AI to improve Alzheimer’s disease diagnosis using MRI. It generates counterfactual maps that hypothetically transform normal brain images to show features of Alzheimer’s, guiding both model learning and refinement. They termed it a learn-explain-reinforce (LEAR) framework that uses these maps to help the model focus on relevant features and improve its generalisation capabilities. This cyclic process of learning and reinforcement was validated with qualitative and quantitative analyses on the Alzheimer’s disease neuroimaging initiative (ADNI) dataset [315], demonstrating enhanced model performance and comprehensibility compared to existing methods.

Metsch et al. [220] created the CLARUS platform (Figure 27 that integrated counterfactual XAI to enhance the understanding and trust in AI models used in clinical decision support systems. Holzinger et al. [316] addressed the inclusion of counterfactual reasoning into explainable AI (XAI) frameworks via Graph Neural Networks (GNNs). Their primary contribution is a multi-modal feature representation space that integrates data modalities such as images, text, and genomics, utilising interaction and correspondence graphs (ICGs) for fusion (Figure 18). Knowledge bases function as links to establish linkages among modalities. Graph Neural Networks (GNNs) are essential for establishing causal relationships between features using graph structures, hence allowing more robust and accessible explanations. This connects AI assessment to individual conceptual comprehension, improving the quality of explanations through causability, a term coined by the authors to assess the efficacy of explanations in facilitating human causal understanding.

**Fig. 18.**
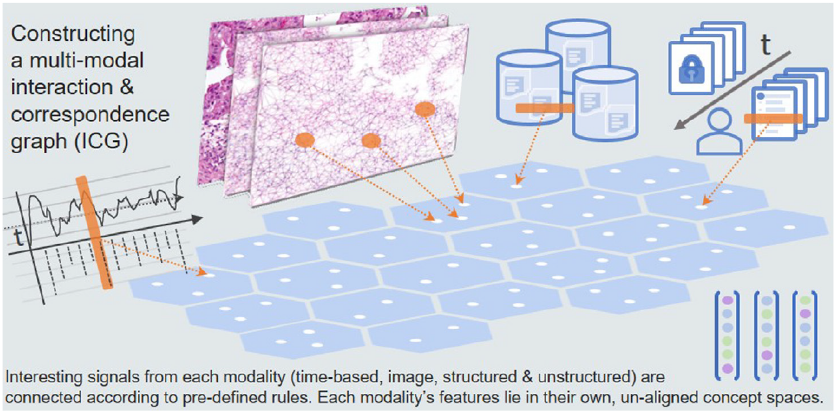
Graph Fusion [316]. This image is licensed under CC BY 4.0.

##### Case-based Reasoning

Vásquez-Morales et al. [233] implemented and validated Neural Network output explanation using case-based reasoning (NN-CBR) dual system. Bichindaritz et al. [317] proposed an explainable AI (XAI) framework for cancer survival analysis using a deep neural network (DNN) and case-based reasoning (CBR). This method largely used an autoencoder network and prototype layer. The prototype layer receives autoencoded training input features. This layer’s weight vector matches the encoded input, making input effects easy to explain. The model’s total loss function includes the negative log partial likelihood from the Cox model (common in survival analysis), the autoencoder loss, and two variables that guarantee intelligible prototype distances. This advanced loss function balances prediction accuracy and model interpretability. Shapley Additive Explanations (SHAP) quantify each feature’s contribution to the model’s predictions to aid explanation. This integrated SHAP-based prototype-based reasoning method completely explains model variables and particular predictions.

A strategy to increase clinician trust in AI-based clinical decision support systems by simplifying AI explanations was described by Corbucci et al. [318]. It presented an approach to use natural language processing (NLP) embedding methods and medical ontologies to enhance AI explanations with relevant clinical information about a patient’s health status. This technique tries to improve the relevance and dependability of the explanations produced by AI by linking data from the patient’s healthcare notes. First tests using a human specialist yielded encouraging findings in determining essential details regarding the patients’ illnesses.

Prototypical Part Network (ProtoPNet) [253] is a deep network architecture intended to improve image classification tasks by simulating human-like reasoning under the case-based reasoning category. ProtoPNet functioned by classifying images by combining evidence from the prototypes it found. This technique is similar to specialists like doctors and ornithologists who analyze images and identify important details to categorize them. ProtoPNet simplifies training by requiring simply image-level labels for training rather than thorough annotations for image components. Multi-modal Prototype Network (MProtoNet) [319] is an extension of ProtoPNet for brain tumour classification utilising 3D multi-parametric magnetic resonance imaging (mpMRI). MProtoNet includes a novel attention module that uses soft masking and an online CAM loss, providing more accurate and relevant attention maps than conventional models that depend on GradCAM. A significant difference from the 2D focus of current models is that this method improves the localisation of attention regions appropriate to the 3D structure of mpMRI data. Crucially, MProtoNet outperformed the metrics of localisation coherence and accuracy, obtaining these outcomes without requiring labels annotated by humans during training.

#### B.3 Interactive Tools

Utilising interactive tools in Explainable AI (XAI) is increasingly crucial for improving the comprehensibility and practicality of AI models, particularly in healthcare. Various platforms and toolkits have been developed to facilitate interactive explanations, allowing users to engage with AI models directly, thus enhancing trust and understanding.

The *Explainable AI Tools v0*.*6*.*0 Toolkit*^10^, created by Intel, provides a suite of tools for understanding and interpreting AI systems. It offers multiple techniques for model interpretability, including feature importance and visual explanations, helping users comprehend the model’s decision-making process. Similarly, *Vertex AI*^11^ integrates explainability capabilities to study and interpret model predictions. The *What-If Tool*^12^ [320], a component of Google Cloud, allows users to analyze model behaviour by modifying input data and observing the resulting prediction changes, which is valuable for detecting biases and understanding feature impacts.

Microsoft’s *InterpretML* framework^13^ [181] supports both interpretable (glass box) and opaque (black box) models. It incorporates SHAP and LIME to provide local and global interpretability of machine learning models, helping users better understand feature contributions and dependencies. Bhattacharya et al. [321] introduced *Explanatory Model Steering (EXMOS)*^14^, a framework enabling users to adjust prediction models interactively. By gathering input from healthcare professionals and patients, EXMOS provides data-based explanations that enhance user interactions and outcomes.

The *CLARUS* platform^15^ [220] is an interactive tool designed for generating counterfactual explanations for Graph Neural Networks (GNNs). It allows domain experts to fine-tune node placements in GNNs to produce accurate and actionable predictions, thereby improving the transparency and reliability of AI systems in clinical settings.

*ProtoPNet*^16^ [253] and *MProtoNet*^17^ [319] enhance image classification tasks by integrating detected prototypes. ProtoPNet is a prototypical part network that mimics human cognitive processes, crucial in medicine for understanding results. MProtoNet extends ProtoPNet to classify brain tumours using 3D multi-parametric MRI, incorporating an attention module to identify key regions in 3D medical data. *TSInterpret*^18^ [322] and *time interpret*^19^ [323] are specialised for interpreting time-series data, providing customised visualisations essential for medical applications where temporal patterns are significant.

*AutoXAI4Omics*^20^ [324] offers interactive visualisation for complex biological data, aiding users in understanding how omic features influence model decisions. The Magnitude Constrained Optimization (MACO) method [325] enhances visualisations in deep neural networks by maintaining the authenticity of generated images. Used in conjunction with the Concept Recursive Activation Factorization (CRAFT) technique [289], MACO visualises extracted feature vectors while ensuring they remain within the domain of natural images, thus improving interpretability. Although MACO does not directly handle multimodal data, its ability to provide spatial meaning to visualisations shows potential compatibility with multimodal medical data. By integrating various clinical data types or medical imaging, MACO offers comprehensive visualisations that enhance the understanding of complex medical processes. The DEEL AI project developed a neural network explainability toolbox named Xplique^21^ to support these advancements.

*Lucid*^22^ by Google improves the interpretability of neural networks, especially those using TensorFlow. It emphasises neuron, layer, and channel activations through techniques like Activation Atlases, saliency maps, Grad-CAM, and integrated gradients, optimising inputs in various representation bases such as Fourier and wavelet transforms. The primary goal of Lucid is to foster open research in neural network interpretability.

These interactive tools and platforms underscore the growing importance of explainability in AI, particularly in critical domains like healthcare. They facilitate user interaction with AI models and enhance understanding of their decision-making processes, bridging the gap between complex AI technologies and user-friendly applications.

### C Hybrid Approach

Alfeo et al. [326] proposed the Boundary Crossing Solo Ratio (BoCSoR) strategy as a means to improve XAI technologies, with a specific focus on classifying mental states based on fMRI data. This approach tackles the drawbacks of conventional counterfactual explanations, which usually concentrate on particular results rather than the general behaviour of the model, and SHAP, which requires a lot of computing resources and is influenced by correlated factors. BoCSoR combines feature importance analysis and counterfactual reasoning to identify how modifications in specific attributes can lead an instance to breach the decision boundary of the model, hence changing the classification outcome. BoCSoR comprehensively evaluates feature importance by combining local counterfactual discoveries and examining occurrences close to this boundary. This approach is specifically designed for analysing fMRI data and provides a more accurate and computationally efficient understanding of how AI models behave when categorising mental states. The authors have proven that BoCSoR improves the comprehensibility of AI models by emphasising the influence of discrete attribute modifications on the overall categorisation.

Wolf et al. [327] expanded upon the ProtoPNet framework [253] by introducing the Prototypical Part Faithful Explanation (ProtoPFaith) approach. ProtoPFaith utilises Shapley values obtained from prototype similarity scores to offer explanations. This approach guarantees that the contribution of each pixel is precisely quantified and elucidated, providing a more authentic representation of the data’s role in predictions. ProtoPFaith improves the clarity and reliability of explanations by following explanation axioms, which increases their comprehensibility and credibility.

These hybrid approaches demonstrate the creative combination of various XAI techniques to address specific restrictions, enhancing AI models’ interpretability and dependability in intricate fields like mental state classification and image-based predictions.

## D Discussion

Approaches to explainable artificial intelligence (XAI) including model-specific, model-agnostic, and intrinsic methods offer distinct advantages and limitations, each suited to different data types and clinical requirements. Model-specific techniques, such as Grad-CAM and CAM, are designed for convolutional neural networks (CNNs) and excel in imaging applications by providing visual explanations highlighting critical regions within medical images. These methods are particularly valuable in scenarios where spatial localisation of features is essential for clinical interpretation, such as identifying abnormalities in radiology scans. On the other hand, model-agnostic methods like LIME and SHAP are adaptable to various machine-learning architectures. They are particularly effective in analysing structured or tabular data, such as electronic health records (EHRs). By offering feature-level explanations, these approaches empower clinicians to understand the contribution of specific variables, making them a preferred choice for data-driven decision-making in healthcare contexts. Their flexibility enables their use across various tasks without being tied to a particular model type. Intrinsic models, including decision trees and Explainable Boosting Machines (EBMs), are naturally interpretable due to their straightforward structure. These methods prioritize simplicity and transparency, making them well-suited for use cases where interpretability is more critical than handling complex data. However, their capacity to address the intricacies of multimodal data, such as combining imaging and EHRs is limited. In such contexts, hybrid approaches that leverage the strengths of multiple techniques are essential. These integrative methods enable a unified interpretation of multimodal data streams, bridging gaps in understanding and supporting robust clinical workflows. This discussion connects the requirements of specific data modalities (outlined in Section 2) with the practical challenges of implementation (detailed in Section 5). While each XAI approach addresses different aspects of explainability, applying these methods in multimodal fusion workflows and frameworks (explored in Section 4) is an area of ongoing development. Continued innovation in this space is critical to ensuring that clinical insights are effectively derived while maintaining technical feasibility. This alignment between clinical utility and technological adaptability is vital for driving the adoption of XAI solutions in healthcare settings.

## 4 Frameworks in XAI Data Visualisation

The emergence of XAI in medical data analysis highlights the necessity for intuitive and robust visualisation frameworks that address the complexity and diversity of clinical settings. XAI visualisation strategies must evolve from static displays into dynamic tools that deliver actionable insights, foster collaboration, and adapt to varied data workflows.

This part examines significant developments in XAI data visualisation frameworks, concentrating on three essential domains: modality and data-specific visualisation techniques, collaborative analysis workflows, and feedback-based explanation enhancement. These frameworks address challenges such as handling high-dimensional data, integrating multimodal inputs, and iteratively improving interpretability through stakeholder feedback. This section analyses cutting-edge methodologies, emphasising their practical uses, intrinsic limitations, and prospects for additional innovation. This discussion explores how XAI visualisation bridges technical complexity and clinical applicability, enhancing transparency, trust, and alignment with healthcare needs. Each subject explores particular approaches, including insights from current studies and their practical consequences.

### A Visualisation techniques tailored to Modalities

Enhancing explainability in medical data analysis requires that techniques correspond to the varied demands of specific fields. Challenges include high-dimensional omics data, real-time decision-making requirements, and multimodal data integration, effectively handled through focused strategies that connect technical solutions with domain-specific workflows. Explainable AI for Dermatology (ExAID-2022) [328] presents an integration of Concept Activation Vectors (CAVs) and Concept Localisation Maps (CLMs) aimed at clarifying dermoscopic features such as colour streaks and irregular borders. This association allows practitioners to link visual patterns with classifications, improving transparency and clinical relevance. For example, heatmaps identify skin lesion characteristics, while textual explanations offer diagnostic context, significantly improving usability in dermatological processes. Despite its potential, ExAID faced challenges with imprecise concept localisation caused by hyperparameter sensitivity in CLM modules. This noise may confuse dermatologists and undermine confidence in model outputs. Resolving this necessitates comprehensive uncertainty quantification identifying uncertain regions, indicating the need for further imaging or physical examination. These modifications could more effectively align the system with physicians’ iterative diagnostic procedures, especially in identifying early-stage melanomas. Figure 19 showcases the overview of this framework where we see the test case presented with the classification of Nevus Lesion in contrast to Melanoma due to regular dots and globules. Concept classification is visible on the right of the test image.

**Fig. 19.**
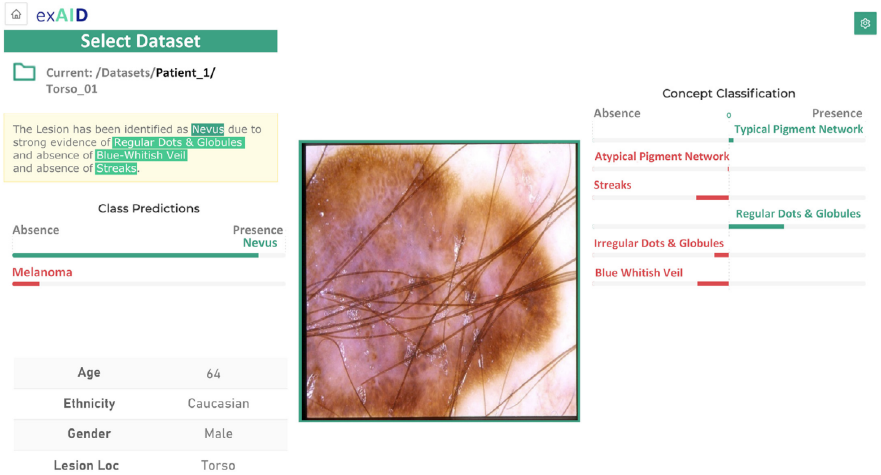
Overview of exAID [328]. This image is licensed under CC-BY-NC-ND.

An even broader approach is illustrated by the Holistic AI in Medicine (HAIM Framework-2022) [329] addressing the fusion of tabular, time-series, textual, and visual data into unified prediction models, utilising Shapley values to explain the contribution of each modality. In clinical applications like 48-hour mortality prediction, HAIM assesses the significance of vital sign changes (time series) and imaging characteristics (e.g., chest X-rays). This profound insight assists physicians in understanding the impact of each data stream on model predictions. Nonetheless, non-monotonic interactions where supplementary material does not uniformly enhance performance represent a considerable obstacle. Including redundant time-series data may diminish the prediction efficacy of imaging features. Dynamic gating methods could deactivate low-value modalities, enhancing computational efficiency and interpretability. This methodology guarantees that the system prioritises high-impact features, particularly in resource-limited healthcare environments. Figure 20 presents the modular architecture of the HAIM framework, demonstrating the integration of various data modalities such as tabular, time-series, textual, and visual into cohesive fusion embeddings. The figure presents the independent preprocessing pipelines for each modality, resulting in a unified multimodal embedding utilised for subsequent predictions. This design emphasises the framework’s adaptability, facilitating tasks such as 48-hour mortality prediction by using the distinct contributions of each data stream while tackling challenges like non-monotonic interactions and redundancy. The visual representation underscores the HAIM methodology’s ability to support scalable and interpretable multimodal AI applications within the healthcare sector.

**Fig. 20.**
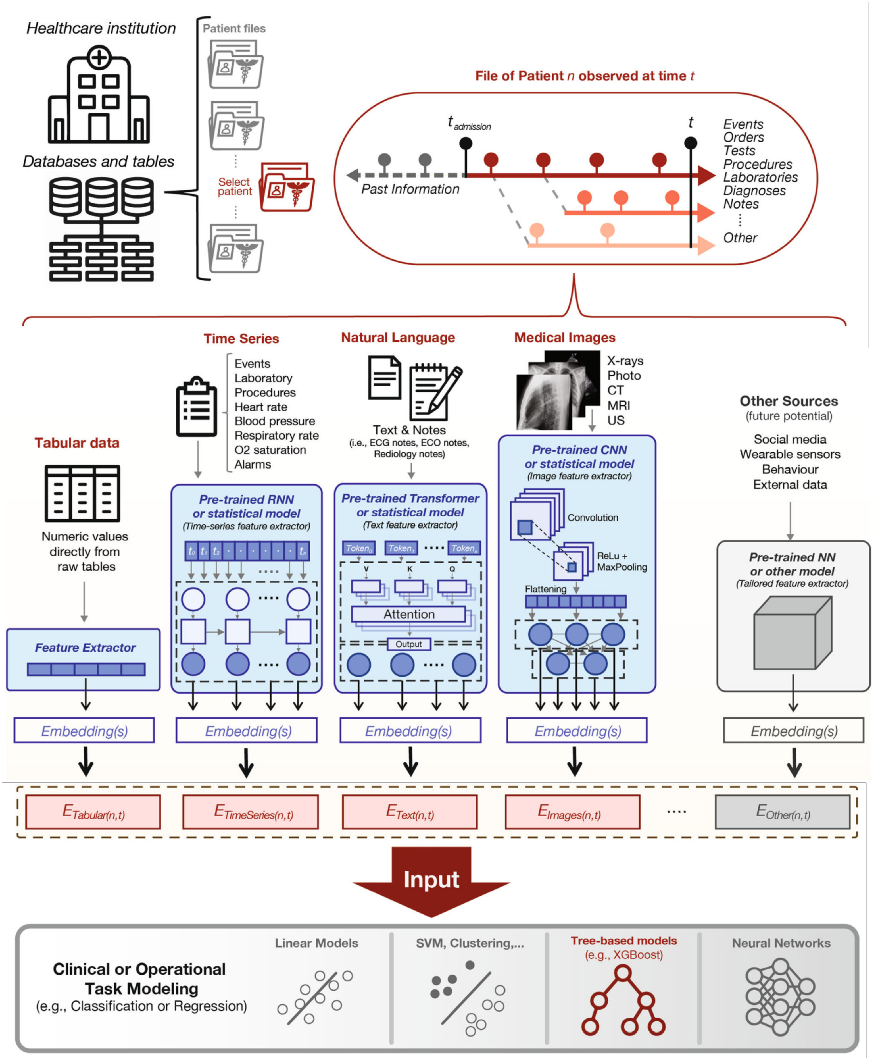
Overview of HAIM [329]. This image is licensed under CC-BY-4.0.

Text conditioned Latent Diffusion Model for Histopathology (PathLDM-2024) [330] utilises text-conditioned diffusion models to produce synthetic histopathology images derived from pathology reports. This novel method integrates GPT-based text summarisation with U-Net latent diffusion, facilitating the enhancement of datasets with high-resolution tumour– tumour-infiltrating lymphocyte (TIL) combinations. The system could generate biologically correct synthetic images, mitigating the deficiency of annotated datasets in pathology research. However, the system’s efficacy is significantly conditional upon the representativeness of its textual training material. Summaries generated by GPT may inadequately convey refined, pathology-specific terminology, restricting their generalisability. An improvement involves utilising extensive, varied pathology literature for training and gating systems to identify and eliminate biologically implausible synthetic results. This would guarantee the dependability of generated images, especially for instructional and diagnostic purposes. Figure 21 presents the PathLDM pipeline, highlighting its fundamental approach to text-conditioned histopathology image generation. The figure illustrates the integration of whole slide images (WSI) with pathology text reports via GPT-based summarisation and patch-level classification of tumours and tumour-infiltrating lymphocytes (TIL). The components are processed using CLIP embeddings and input into the latent diffusion model (LDM), utilising a Variational Autoencoder (VAE) alongside a U-Net denoiser. This configuration facilitates the generation of biologically precise, context-sensitive histopathology images, tackling issues such as restricted training data and involved multimodal interactions. The visual representation highlights PathLDM’s ability to improve pathology datasets and its potential utility in diagnostic applications.

**Fig. 21.**
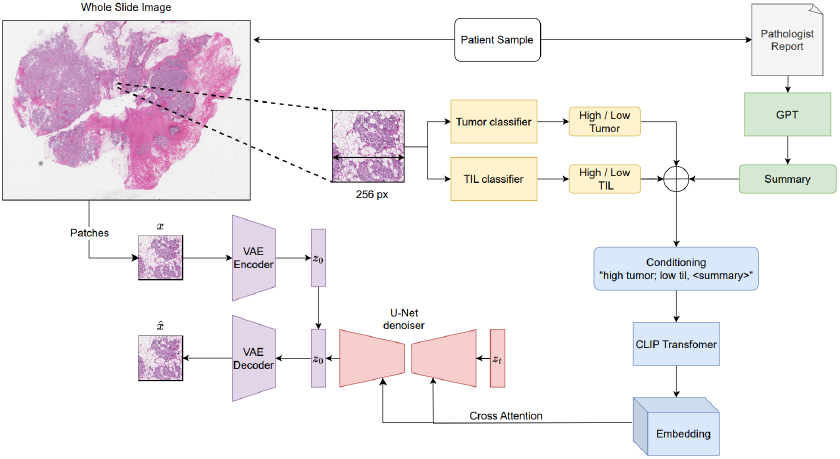
Overview of PathLDM [330]. This image is licensed under CC-BY-4.0.

Moving to neurosurgical applications, image-guided neurosurgery (NeuroIGN-2024) [331] framework combined pre-operative MRI with real-time intraoperative ultrasound (iUS) to assist surgeons during tumour removal. Certainty maps produced by Grad-CAM superimpose segmentation confidence scores onto live surgical displays, thereby improving decision-making in high-pressure environments. However, accurate synchronisation of sensors continues to be a challenge. Misalignments or latency may compromise the system’s reliability, perhaps resulting in erroneous surgical guidance. Advanced sensor fusion algorithms that consider patient movement and intraoperative variations could lessen these problems. Furthermore, integrating 3D visualisations of tumour margins might lessen the cognitive burden for surgeons alternating between MRI and ultrasound images. Implementing real-time confidence thresholds may notify surgeons when AI forecasts drop below acceptable accuracy levels, necessitating manual intervention. Figure 22 illustrates the NeuroIGN workflow utilised in multimodal image-guided neurosurgery. The process begins with acquiring pre-operative imaging data, comprising MRI and intraoperative ultrasound (iUS), which are subsequently integrated and aligned through AutoSeg for tumour segmentation (Step 1). Patient registration (Step 2) aligns the physical body with a virtual model, facilitating the precise tracking of surgical tools in real-time (Step 3) and their synchronisation with live ultrasound streams (Step 4). Explainable AI (Step 5) uses Grad-CAM to produce certainty maps, overlaying confidence scores onto real-time imaging. This research informs surgical navigation (Step 6) by integrating multimodal data, providing dynamic and interpretable visualisations for surgeons. The workflow enhances decision-making and surgical accuracy, ultimately facilitating successful tumour removal (Step 7). This figure highlights the modular, integrated, and explainable characteristics of NeuroIGN, emphasising its potential to improve accuracy and reliability in critical clinical situations.

**Fig. 22.**
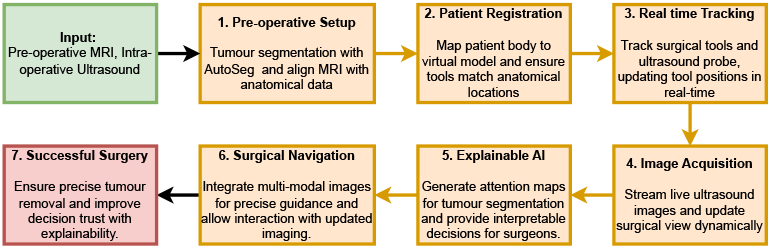
Overview of NeuroIGN [331]

In the domain of structured and unstructured data, EHR-KnowGen (2024) [332] presented a generative framework that integrated structured EHR data (such as laboratory findings and diagnostic codes) with unstructured clinical notes, converting them into textual prompts for a T5-based model. This method demonstrated superior efficacy in diagnosing intricate illnesses by capturing cross-modal interactions. Nonetheless, the framework’s dependence on rigorous hyperparameter optimisation presented scalability challenges across various healthcare systems. Automated calibration methods could identify domain drift changes in data distributions across geographies or demographics, and need to modify the model parameters accordingly. Moreover, improved dashboard designs could emphasise obstacles, such as absent lab tests, assisting physicians in prioritising data collection to boost diagnosis. Figure 23 presents the modular workflow of EHR-KnowGen, demonstrating the processing of structured and unstructured EHR data through its three primary components. The multimodal encoder integrates various data modalities into cohesive textual representations, forming the basis for further processing. The disease-related information extractor incorporates external domain knowledge through hierarchical ICD codes, facilitating the extraction of clinically relevant features and aligning multimodal embeddings to enhance interpretability and precision. The diagnosis results generator employs enhanced embeddings to produce disease diagnoses, utilising a generative approach to address complex conditions. This figure illustrates the framework’s ability to integrate multimodal data and domain knowledge into a unified diagnostic process.

**Fig. 23.**
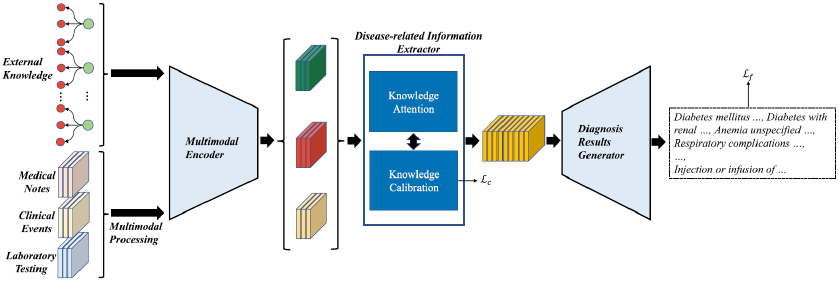
Overview of EHR-KnowGen [332]. This image is licensed under CC BY 4.0.

The XAI Orchestrator (2024) [333] united the imaging, genomics, and electronic health record data into a cohesive platform designed for oncology. Integrating layered saliency maps and aggregated Shapley values offered oncologists longitudinal insights into illness progression. A primary difficulty is addressing discrepancies in data quality across modalities, including inadequate electronic health record entries and inconsistent imaging techniques. Establishing a tiered validation system to pre-screen low-quality inputs could enhance the integration process. Furthermore, incorporating coherence scores would facilitate the identification of contradictory signals, such as a tumour showing malignant characteristics genomically while seeming benign radiographically, necessitating additional inquiry. Figure 24 presents a conceptual overview of the XAI Orchestrator framework, aimed at integrating multimodal and longitudinal data into a cohesive platform for oncology. The figure illustrates the integration of clinical guidelines and recent research as a foundational knowledge base, merged with patient-specific multimodal data to forecast outcomes. The framework employs layered saliency maps and aggregated Shapley values to produce modality-specific or time-specific explanations, subsequently synthesised by the Orchestrator into a comprehensive explanation. This explanation, made by an advanced XAI system, aims to assist oncologists by structuring and articulating insights and facilitating followup questions. The illustration emphasises the system’s modular architecture and capacity to manage various data inputs, ensuring adaptability and user-focused interaction.

**Fig. 24.**
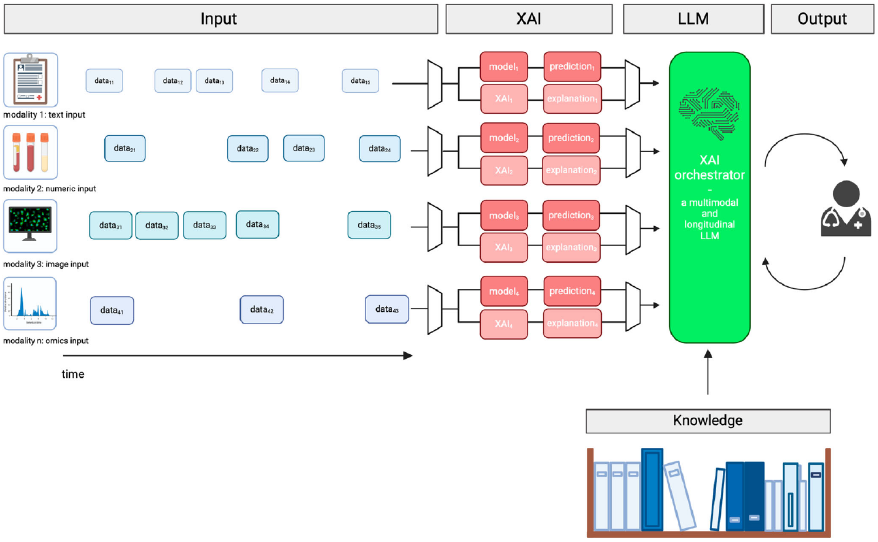
Overview of XAI Orchestrator [333]. This image is licensed under CC-BY-4.0.

Automated Explainable AI for Omics (AutoXAI4Omics-2024) [324] offered an automated pathway for omics analysis, utilising SHAP to associate genomic and proteomic characteristics with outcomes such as disease states or phenotypes. This framework is particularly good at discovering plant phenotyping or microbiome analysis. The dependence on human preparation for categorical data is a considerable obstacle for domain experts lacking familiarity with machine learning processes. Implementing automated feature encoding methods like natural language processing tokenisation may reduce the entrance barrier. Moreover, interactive lectures or organised training programs would assist users in more effectively interpreting SHAP attributions, thereby connecting AI outputs with biological pathways. Figure 25 illustrates the AutoXAI4Omics framework, showcasing its comprehensive workflow for analysing omics and tabular data. The figure portrays the sequential process, commencing with data ingestion and optional pre-processing steps, encompassing filtering, normalisation, and scaling, designed explicitly for omics data types. Subsequently, automated feature selection and hyperparameter tuning are utilised to determine the optimal machine learning models for the specified tasks, including classification and regression. The chosen models undergo evaluation and ranking according to various performance metrics, leading the framework to recommend the most appropriate model for subsequent analysis. Interpretability tools, such as SHAP, provide explainable insights into the relationships between features and predictions, thereby aiding in identifying actionable biological findings. This workflow highlights automation, interpretability, and adaptability, allowing domain experts to utilise machine learning without in-depth model development or optimisation knowledge.

**Fig. 25.**
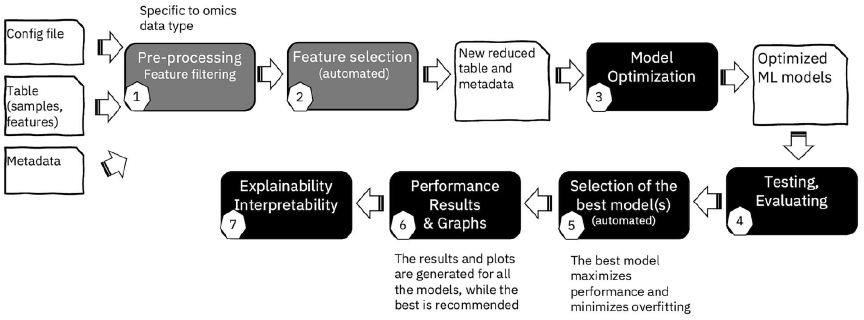
Overview of AutoXAI4Omics [324]. This image is licensed under CC BY 4.0.

These studies revealed the challenge of balancing domain-specific explainability with the need for generalisable frameworks that handle diverse datasets. Researchers must address challenges such as noisy localisations, real-time synchronisation, and multimodal fusion through improvements in dynamic gating, uncertainty quantification, and automated preprocessing. By tackling these issues, future systems can more effectively assimilate into clinical workflows, guaranteeing that AI tools not only forecast outcomes but also deliver actionable, reliable explanations that correspond with the requirements of healthcare professionals.

### B Workflows for collaborative analysis

XAI systems include uncovering model working and creating intuitive interfaces and feedback loops to promote collaboration among clinicians, doctors, patients, and data scientists. These methods cultivate a standard comprehension of AI results, improving trust, usability, and acceptance. We examined various collaborative frameworks designed to enhance the transparency and user-centricity of AI-driven multimodal healthcare systems.

In neurological disorders, Graph Convolutional Networks (GCN-2024) [334] are notable for their ability to engage clinicians directly in comprehending Alzheimer’s disease categorisation. The system utilised techniques such as adjacency matrices, bar graphs, and neighbourhood analyses, enabling doctors to assess the impact of patient similarities on diagnostic judgements. These discoveries are especially significant when examining the transition from moderate cognitive impairment (MCI) to Alzheimer’s disease. A considerable issue exists in differentiating MCI from normal ageing, as overlapping characteristics among subgroups might obscure diagnostic distinctions. One potential improvement is confidence-based stratification, which flags ambiguous cases as ‘inconclusive’ and prompts further testing or consultation. Furthermore, a reliability index demonstrating the consistency with which particular attributes distinguish MCI from normal cognition across diverse cohorts or model iterations could give doctors more confidence in the system’s recommendations. Figure 26 illustrates the computational architecture of Graph Convolutional Networks (GCNs) utilised to classify Alzheimer’s disease. Patient data, depicted as nodes, are interconnected by edges weighted according to their feature similarities, creating a population graph. The GCN utilises spectral convolutions to consolidate neighbourhood information, incorporating ReLU activation and self-connections to enhance the learnt embeddings. The model generates predictions for each node, categorising patients into Normal Cognition (NC), Mild Cognitive Impairment (MCI), or Alzheimer’s Disease (AD). The explanation module disaggregates feature contributions at individual and group levels, clarifying the reasoning underlying predictions. This visualisation highlights the graph-based methodology and technological advancements that facilitate precise integration and interpretation of multimodal data.

**Fig. 26.**
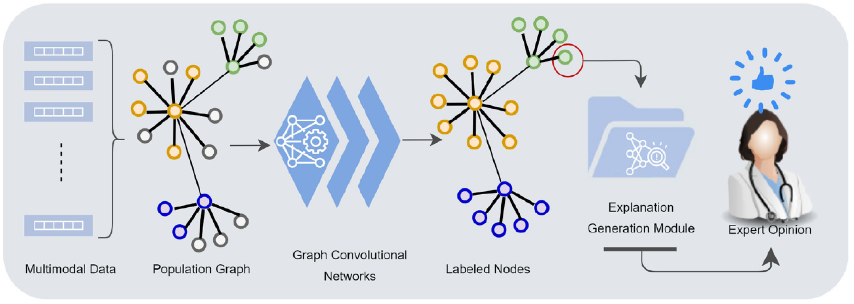
Overview of GCN [334]. This image is licensed under CC BY 4.0.

A similar drive toward interactive analysis characterised CLARUS (2024) [220], facilitating counterfactual reasoning for domain specialists utilising graph-based omics data. Clinicians and researchers could manually alter nodes or edges in a graph neural network (GNN) to evaluate “what if” situations. For example, eliminating a gene node in a protein-protein interaction (PPI) network may explain its influence on disease predictions, such as kidney cancer. These counterfactual experiments offer valuable clarity but require substantial manual intervention and domain expertise. The procedure can be optimised by integrating semi-automated recommendations that emphasise potentially significant nodes or edges informed by previous analysis or domain expertise. By easing cognitive strain, such enhancements would facilitate accelerated hypothesis testing while preserving the interactivity that renders the system very effective. Figure 27 presents the workflow of the CLARUS platform, an interactive explainable AI tool developed for counterfactual reasoning utilising graph neural networks (GNNs). The figure illustrates user engagement with the platform by selecting datasets, including synthetic protein-protein interaction (PPI) networks and real-world kidney cancer data. The workflow consists of several essential stages: exploration of visualised graphs, manual counterfactual manipulations (such as the addition or deletion of nodes or edges), prediction updates, and optional GNN retraining. These steps enable domain experts to assess the effects of structural changes on network predictions and explanation scores, thereby enhancing causal insights. The illustrated API connections (red arrows) indicate real-time communication between the user interface and the backend, facilitating dynamic updates to predictions and relevant visualisations. This integration enhances the platform’s functionality in iterative hypothesis testing, enabling users to analyse the impact of specific genes or interactions on disease classification results. The workflow illustrates the CLARUS system’s dedication to transparency, interactivity, and accountability in biomedical decision-making.

**Fig. 27.**
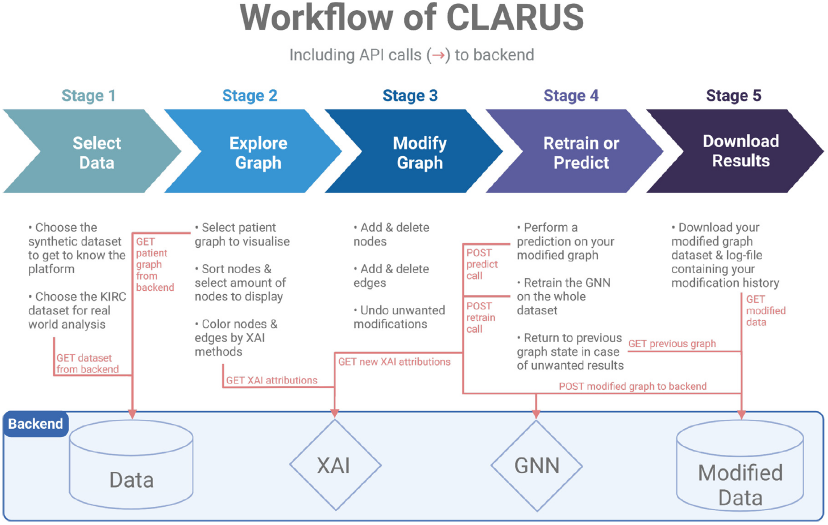
CLARUS workflow including API calls [220]. This image is licensed under CC BY 4.0.

Calibrated-XAI (2024) [222] illustrated collaborative workflows by integrating a trust calibration procedure into its multimodal AI architecture. The framework employed role-specific explanatory interfaces, clinicians authenticate AI outputs via “stamping” procedures while the IT teams investigate confidence intervals and evaluate system robustness. This participatory method guarantees that every stakeholder aids in enhancing the system. Nonetheless, sustaining this stakeholder engagement presents a logistical difficulty in busy clinical settings. Hospitals frequently lack the capacity for recurrent feedback sessions. Potential improvements may include streamlined user feedback systems, such as brief questionnaires embedded inside the workflow, that gather critical insights without necessitating time-consuming workshops. These micro-interactions may enhance regular feedback sessions to facilitate ongoing system enhancements. Figure 28 summarises the Calibrated-XAI (C-XAI) framework, highlighting its systematic approach to the design of trust-calibrated XAI interfaces. The diagram defines the four primary phases of the framework: Identification, Assessment, Selection and Implementation, and Evaluation. Each phase includes targeted activities designed to connect the technical attributes of XAI methods with human-centred design objectives. During the Identification phase, system analysts and stakeholders work together to collect requirements for explanations pertinent to the target human-AI task, ensuring inclusivity and representativeness. The Assessment phase analyses the technical and human factors associated with XAI methods, pinpointing risks that may result in trust calibration errors. The third phase of the framework, Selection and Implementation, focuses on risk mitigation through specific design interventions informed by C-XAI’s templates and best practices. The Evaluation phase facilitates iterative design refinement through behavioural metrics to evaluate trust calibration outcomes and user interactions. The figure illustrates the participatory design approach central to C-XAI, highlighting the active involvement of multidisciplinary stakeholders, including system analysts and psychologists, to ensure that the resulting XAI interfaces balance technical rigour and usability. This structured workflow facilitates the creation of interfaces that enhance trust in AI systems, especially in critical decision-making contexts.

**Fig. 28.**
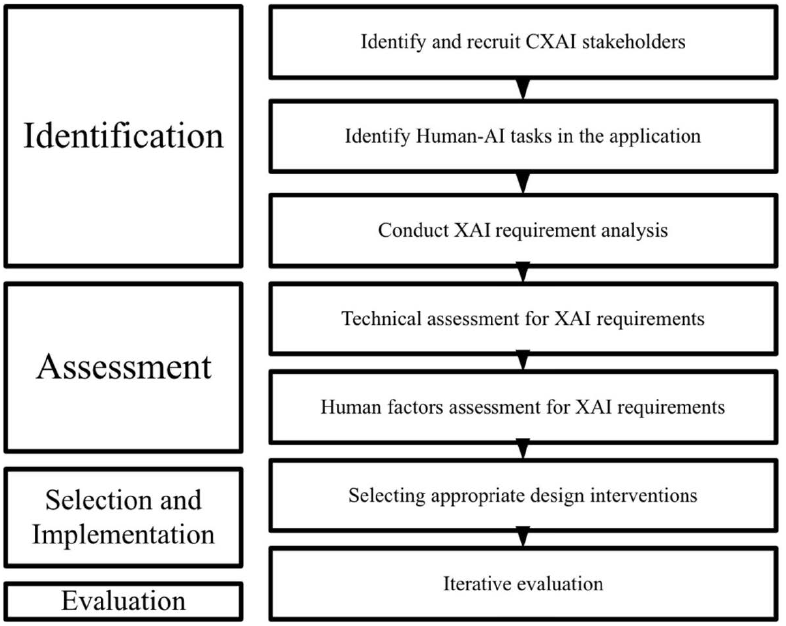
Overview of Calibrated-XAI [222]. This image is licensed under CC-BY-NC-ND.

In imaging-centric specialities, the Clinical Conversational AI Interface (2024) [335] enhanced radiological operations by enabling clinicians to communicate with the system via natural language queries. A radiologist may request a heatmap of a worrisome lesion and receive a visual representation and textual rationales. This dialogue-based design reduces adoption hurdles for radiologists, who often favour verbal communication over technical terminology. However, achieving a balance between diagnostic precision and user-friendly engagement continues to be complicated. For example, when confronted with complex enquiries or significant uncertainty, the system may yield partial or overly simplistic answers. To resolve this, establishing organised fallback mechanisms, such as an “expert mode” or a second-opinion feature, could direct users towards more comprehensive or alternate explanations, maintaining the system’s reliability in intricate situations. Figure 29 illustrates the Clinical Conversational AI Interface, emphasising its multimodal workflow to improve radiological operations. The figure describes the process, commencing with identifying user needs via analysing prevalent clinician enquiries, including diagnosis, location, and severity. The workflow advances the design of multimodal responses that integrate textual elements, similar to radiologist-style explanations, with visual aids such as heatmaps, segmentation maps, and contrastive images. The subsequent step focuses on validating the relevance of response content through dataset analysis to distinguish between essential details (e.g., location and severity) and superfluous information (e.g., general knowledge already known to clinicians). User testing and feedback follow, during which clinicians engage with prototypes that provide switching options between text-only and multimodal interfaces to assess usability and efficiency. The text illustrates the balance between diagnostic accuracy and usability, emphasising the need for effective performance in life-threatening and difficult-to-diagnose conditions while minimising false alarms. The iterative refinement process integrates clinician feedback, including larger images and additional switching options, to create a clinically optimised multimodal conversational system. This figure illustrates the system’s iterative design approach, focused on enhancing usability and adoption while preserving the accuracy required for clinical decisionmaking.

**Fig. 29.**
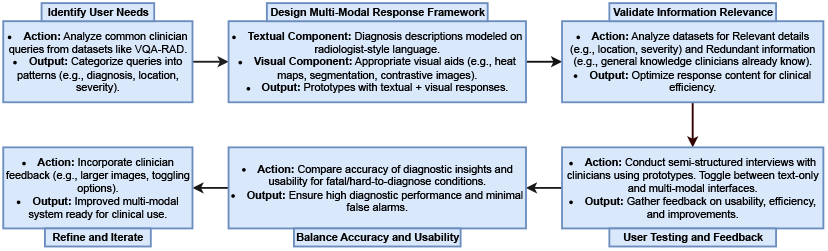
Overview of Clinical AI Interface [335]

A more text-focused approach appears in VLM-Radiology (2024) [309] that utilised vision-language models (VLMs) to integrate imaging and textual data smoothly. This framework enabled AI-assisted draft report generation and case retrieval through visual searches. The human-centred design, consistently enhanced by clinician feedback, promotes usability and guarantees conformity with radiological procedures. Nonetheless, maintaining automated textual outputs’ clinical accuracy and contextual relevance is a pivotal concern. Misclassifications, including mixing analogous circumstances or neglecting critical discoveries, can reduce trust in the system. An enhancement would be a domain-specific fine-tuning phase that cross-verifies outputs with a curated repository of prevalent diagnostic inaccuracies. Furthermore, a real-time validation system, incorporating crowdsourced input from various radiologists, could identify discrepancies or omissions before the finalisation of reports.

These collaborative workflows highlight the necessity of matching advanced AI systems with the actual requirements of clinical settings. These frameworks enhance comprehension of AI results and cultivate user trust through role-specific interfaces, interactive explanations, and participatory feedback systems. Nevertheless, difficulties such as minimising cognitive load, streamlining interfaces, and addressing ambiguity continue to be essential to their enhancement. By overcoming these obstacles, future systems can attain an optimal equilibrium among usability, diagnostic precision, and stakeholder involvement.

Table 8 presents the overview of various XAI studies over multimodal medical data, highlighting the objective of the study, various methods used, problems solved, and how it is helpful in multimodal medical data.

**Table 8.** Overview of XAI Studies in Multimodal Medical Data

| Study | Objective | Modalities Used | Fusion Strategy | Explainability Approach | Visualisation Tools | Performance Metrics | Applications and Challenges |
| --- | --- | --- | --- | --- | --- | --- | --- |
| AutoXAI4Omics (2024) [324] | Automated explainable AI tool for omics and tabular data | Genomics, Transcriptomics, Proteomics, Microbiome | Automates feature selection, hyperparameter tuning, and model selection for single-modality input; supports pre-processed multimodal integration | SHAP for global/local feature importance, combined with eli5's permutation importance | SHAP summary plots, feature importance bar plots, ROC curves, confusion matrices, interactive feature analysis | F1-Score, Accuracy, MAE, MAPE, R2 | Applications: Phenotype and disease state prediction, transcriptomics (e.g., infection response), environmental microbiome analysis<br>Challenges: Requires manual preprocessing for categorical features, adding complexity for non-technical users |
| Calibrated-XAI (2024) [222] | Participatory framework for trust calibration in XAI interfaces | EHR, Imaging data | Applies participatory design for multimodal data handling | Combines task-specific explanations with domain-context validation via interactive features and tailored outputs | Interactive prototypes, validation tools, visual feedback | Completeness, Usefulness, Effectiveness | Applications: Multimodal healthcare interfaces for trust calibration<br>Challenges: Stakeholder involvement extends design cycles, and scalability to diverse scenarios is limited by manual tailoring efforts |
| CLARUS (2024) [220] | Interactive XAI for counterfactuals in graph neural networks | Gene Expression, Methylation | Patient-specific graphs with multi-omics data, using node and edge relevances | Combines GNNExplainer for node importance with Integrated Gradients and Saliency for edge attributions; supports counterfactual generation via manual graph edits | Graph visualisations for node/edge relevance, interactive tools for graph manipulation, dynamic relevance score tooltips | Sensitivity, Specificity, Confusion Matrix | Applications: Cancer diagnosis (e.g., Kidney renal clear cell carcinoma-KIRC, other cancers)<br>Challenges: Limited datasets reduce generalizability; human-in-the-loop refinements increase expert workload |
| Clinical Conversational AI Interface (2024) [335] | Multimodal conversational interface for clinical imaging systems | X-ray, MRI, CT, Radiology Reports | Combines textual descriptions and visual explanations using rule-based matching and inductive coding for disease-specific responses | Aligns visuals with clinician queries via thematic analysis, using radiologist-style language and tools like segmentation maps and heatmaps | Heatmaps, segmentation maps, contrastive visuals, and progression tracking | Diagnostic precision, User satisfaction, Query response coverage | Applications: Enhances clinical workflows and AI adoption through conversational interfaces<br>Challenges: Balancing accuracy with usability, managing false alarms, and catering to diverse clinician preferences |
| EHR-KnowGen (2024) [332] | Generative framework for integrating multimodal EHR data for disease diagnosis | Medical notes, clinical events, lab results | Uses soft prompts for multimodal EHR unification; T5-based transformer encodes fine- and coarse-grained domain knowledge | Knowledge attention for disease identification, cross-attention for embedding alignment, and hierarchical calibration for integration | Attention maps, disease progression visualisations, textual diagnostic outputs with evidence | Micro/Macro F1, Precision, Recall, Accuracy | Applications: Rare/complex disease diagnosis using multimodal EHR<br>Challenges: Requires high-quality datasets, precise tuning of hyperparameters and soft prompts, increasing deployment complexity |
| ExAID (2022) [328] | Skin lesion diagnosis with human-aligned explanations | Dermoscopic, clinical images; textual explanations | Combines Concept Activation Vectors (CAVs) and Concept Localisation Maps (CLMs) for feature representation; rule-based explainers for visual and textual outputs | Uses CAVs to detect concepts, CLMs to localize features, and textual explanations for diagnostic insights, addressing uncertainty and contraindications | Heatmaps, progression maps, and textual narratives summarising diagnoses | Accuracy, Precision, Recall, AUC (Lesion Classification); Macro F1 (Concept Classification) | Applications: Dermatological diagnoses (e.g., melanoma, nevi)<br>Challenges: Limited concept-annotated datasets, noisy CLMs, hyperparameter tuning issues |
| ExPeRT (2024) [336] | Explainable prototype-based model for continuous regression tasks | Adult MRI, Fetal Ultrasound | Processes image patches via optimal transport to compute and aggregate patch-wise distances | Prototype learning explains predictions by mapping patches to prototypes in latent space, aligning predictions with label differences for interpretability | Prototype visualisations, patch-matching matrices, dynamic prototype weighting | Mean Absolute Error (MAE), with ablation studies for OT and consistency loss | Applications: Brain age prediction (adult MRI), fetal neurodevelopment patterns<br>Challenges: Requires high-quality images, precise preprocessing, and is computationally intensive |
| **GCNs (2024) [334]** | Graph convolutional network for Alzheimer's classification using multimodal data | MRI, PET, Cognitive Tests, Genetic, Biomarker Data | Population graph with patient nodes, feature-based edges; multimodal data fused into node attributes, edges reflect cognitive and biomarker similarities | Decomposition-based explanations for individual, group, and relational impacts; edge silencing for group influence; counterfactual/factual insights for clinicians | Adjacency matrices, feature importance plots, counterfactuals, pie charts for feature impacts, relational influence analysis | Accuracy, F1-Score, Explanation Stability, Expert User Agreement | Applications: Alzheimer's classification (Normal, MCI, Alzheimer's); biomarker identification<br>Challenges: Overlapping features (MCI/Normal), preprocessing complexity, high graph construction cost |
| **HAIM Framework (2022) [329]** | Holistic multimodal AI framework for healthcare prediction | Tabular, Time-series, Text, Images | Modular fusion embeddings aggregate standardised features across modalities for unified representation | Shapley values quantify contributions at feature and modality levels, providing insights into information flow and error propagation | ROC curves, Shapley value waterfalls, interaction heatmaps, gradient plots, and interactive tools for interpretability | AUROC, Precision, Recall, Average Improvement Metrics | Applications: Chest pathology diagnosis, length-of-stay, and mortality predictions with improved accuracy over single-modality models<br>Challenges: Non-monotonic fusion effects, data sparsity, redundancy, preprocessing demands, and reliance on high-quality multimodal data |
| **LAVA (2024) [337]** | Neuron-level explainable AI framework for Alzheimer's assessment | Retinal Fundus Imaging, Cognitive Tests | Fuses retinal biomarkers and cognitive scores in a hierarchical framework using vessel segmentation, neuron probing, and granularity explanations | Two-phase explanation: (1) Neuron probing via MI identifies critical neurons; (2) Granularity explanation clusters subpopulations with hierarchical clustering. Visual backpropagation aids biomarker localisation | Dendrograms, feature attribution maps, radar plots, progression visualisations, gradient saliency maps | Accuracy (AD vs. Normal), Clustering Metrics (Calinski-Harabasz, Adjusted MI), Vascular Morphology (Fractal Dimension, Vessel Density) | Applications: Alzheimer's classification and progression mapping for early intervention<br>Challenges: Small dataset (100 images, 61 patients), demographic confounders, and high sensitivity to clustering/neuron selection parameters |
| **NeuroIGN (2024) [331]** | Explainable multimodal system for brain tumour surgery | Pre-operative MRI, Real-time iUS, Tracking Data | Integrates MRI with iUS for real-time navigation, dynamically updating surgical views using multimodal data fusion | Grad-CAM for interpretable segmentation, ExplainAI module for certainty maps and guided visualisation, counterfactuals for tumour boundary delineation | Certainty maps, segmentation overlays in 3D Slicer, guided saliency maps, interactive navigation with MRI and iUS | Dice Similarity Coefficient (DSC), Hausdorff Distance (HD95), Target Registration Error (TRE), Real-time frame rate (19 FPS) | Applications: Brain tumour resection, adaptable to neurosurgery tasks, improving precision and AI trust<br>Challenges: Balancing segmentation accuracy with explainability, UI intuitiveness, support for 3D imaging, and real-time data calibration |
| **PathLDM (2024) [330]** | Text-conditioned Latent Diffusion Model for histopathology image generation using pathology reports | Whole Slide Images (WSIs), Pathology Reports | Fuses global WSI-level data with local tumour/TIL statistics using GPT summaries and CLIP embeddings for unified text-visual representation | GPT-based text summarisation and CLIP text embeddings guide U-Net diffusion in latent space. Includes cyclical positional embedding and classifier-free guidance for improved generation | High-resolution histopathology images, progression maps, tumour/TIL heatmaps, FID plots | FID, SSIM, CAS, Tumour/TIL Class Consistency | Applications: Synthetic histopathology for breast cancer research, aiding data augmentation and tumour biology insights<br>Challenges: Relies on text summaries, computationally intensive, complex integration of tumour/TIL at patch level |
| **Prosthetic Hand Control (2022) [338]** | Optimisation of multimodal sensors for prosthetic hand control using explainable AI | EMG, Accelerometer, Gyroscope, Magnetometer | Combines sensor data at feature and decision levels; uses explainable AI to optimize sensor configurations and analyze modality contributions | Gradient-based methods (IG, Gradient SHAP, Saliency Maps) ensure robust attribution via metrics like faithfulness, sensitivity, and complexity | Heatmaps for temporal/spatial attributions, contribution maps for sensor placement, and modality-specific importance charts | Recognition Accuracy (91.5% healthy, 79.2% amputees), Attribution Faithfulness, Sensitivity, Complexity | Applications: Enhances prosthetic hand control, reducing sensors by 40% while maintaining performance<br>Challenges: Sensitive to sensor placement and data quality, computationally intensive, and requires fine-grained tuning |
| **RSA (2024) [339]** | Evaluates explanation stability in XAI across ML pipelines | Chest X-rays, Tabular Data (Heart Disease, Housing Prices, Multi-class Data) | Post-hoc RSA methodology to measure explanation similarity across pipelines, models, preprocessing, and feature engineering | Uses RSA to analyze Representational Dissimilarity Matrices (RDMs) and similarity metrics (Cosine Similarity, PCC, Kendall's Tau) for explanation robustness under varied conditions | Heatmaps for RDMs, scatter plots for similarity, feature contribution, and pairwise similarity visualisations | Classification: Accuracy, Recall, Precision, F1;<br>Regression: R2, MAE, RMSE;<br>Similarity: Sc, PCC, $\tau_A$ | Applications: Stability analysis of XAI explanations in healthcare (COVID-19, heart disease) and real estate. Aids robust explainer selection.<br>Challenges: Requires high-dimensional data, performance depends on similarity measures, and RDM calculations are computationally intensive for large datasets |
| **VLM-Radiology (2024) [309]** | Vision-Language Models (VLMs) for radiology workflows | X-ray, CT, Textual Reports | Combines multimodal embeddings of image-text pairs to enable report generation, visual search, and workflow enhancements | Iterative feedback from clinicians shapes four use cases: draft report generation, augmented review, visual search, and imaging history highlights | Interactive tools for image annotation, report generation, visual queries, and longitudinal comparisons | Focus on clinical usability and qualitative feedback. No standardised metrics reported | Applications: Report generation, clinical visual search, imaging history highlights, and decision support<br>Challenges: Clinical integration barriers, trust in AI, scalability with iterative feedback dependency |
| **XAI Orchestrator (2024) [333]** | Framework for orchestrating XAI in multimodal and longitudinal medical data | Radiology Imaging, EHRs, Genomic Data | Fuses multimodal and longitudinal data with adaptive, uncertainty-aware mechanisms; employs modular architectures for dynamic updates | Integrates saliency maps, Shapley values, delta saliency maps, and hierarchical layers for context-aware, user-adapted explanations | Visualisations include color-coded saliency maps, feature importance charts, delta saliency maps, and interactive tools for clinician engagement | Faithfulness, Robustness, Localisation, Complexity, and Randomisation metrics; AUROC, precision, and F1-score for modular evaluations | Applications: Oncology and radiology workflows (e.g., tumour boards), supporting multimodal interpretation and longitudinal tracking<br>Challenges: Data sparsity, high-dimensional -omics data, adversarial vulnerabilities, variability across sources, and computational demands for scalable integration |

### C Feedback-based explanation refinement

Many AI systems generate fixed explanations, but newer approaches treat interpretability as a dynamic, iterative process. These frameworks integrate feedback from domain experts or models to enhance explanations progressively, enhancing their resemblance with actual medical knowledge and clinical practices.

In rehabilitation engineering, the study on Prosthetic Hand Control (2022) [338] employed gradient-based attribution methods to determine the essential sensors, including electromyography (EMG) and accelerometers, for precise hand-motion identification. Through the iterative analysis of feature importance metrics, the framework attains a 40% reduction in sensor utilisation while maintaining recognition accuracy. This decrease streamlined the hardware combinations while preserving system performance. The reliance on precise sensor placement introduced variability, as misaligned sensors can reduce performance. The computing burden of techniques like Integrated Gradients or Saliency Maps complicates the real-time implementation of prosthetic devices. A potential enhancement would be to create on-device approximation techniques for feature attribution, potentially utilising hardware accelerators to preserve computational efficiency in embedded settings. A further improvement may include a customisation phase, wherein the system adaptively modifies sensor significance according to each user’s limb morphology and residual muscular signals. This method would create a flexible pipeline tailored to individual anatomies, supporting personalised rehabilitation. Figure 30 illustrates the workflow of the XAI-enhanced human-machine interface system developed for prosthetic hand control. The process initiates with Input Data Collection, involving acquiring multimodal sensor data, such as electromyography (EMG) and inertial measurement units (IMU), from healthy individuals and amputees. The subsequent phase entails the construction of the Target Model, wherein a recognition model is formulated to analyse sensor data, thereby identifying hand gestures and their corresponding actions. Explainable AI (XAI) techniques, including saliency, integrated gradients, and gradient SHAP, are utilised to reveal the contributions of individual sensors or modalities. These methods produce Contribution Maps that visually emphasise the sensors or modalities that substantially affect recognition accuracy. The system performs a Quantitative Evaluation utilising metrics such as faithfulness, robustness, complexity, and randomisation to verify the reliability and interpretability of the explanations. Ultimately, the system optimises sensors by removing redundant sensors and streamlining the hardware configuration while preserving or enhancing recognition accuracy. This figure highlights the iterative method for optimising multimodal sensor systems, minimising complexity while preserving explainability and precision. The methodology is enhanced by visually illustrating the systematic steps for improving prosthetic hand control using XAI techniques. ExPeRT (2024) [336] illustrated the iterative refinement in continuous regression tasks, such as foetal brain age estimation. The system offered straightforward, specific reasons for its predictions by correlating visual patches with prototypes in a learnt latent space. Experts can identify mismatched prototypes which are the instances when the model erroneously correlates attributes like bone density with advanced brain age, necessitating recalibration of the latent space in subsequent training iterations. This approach facilitated a continuous feedback loop, enhancing the model’s dependability and domain alignment. Nonetheless, using ExPeRT in resource-limited clinical settings presented difficulties because of its need for extensive collections of high-quality pictures. A viable alternative is establishing a federated learning system consolidating datasets from several sources while safeguarding patient privacy. This method would enable the model to enhance its prototypes, utilising various data sources without necessitating the distribution of raw images. Furthermore, employing data augmentation approaches may improve prototype generalisability, allowing the model to function effectively even with limited information. Figure 31 presents a schematic representation of the ExPeRT model architecture and its methodology for explainable prototype-based regression tasks. The workflow comprises three primary components. The model architecture initially extracts latent features from an input image using a convolutional encoder and supplementary processing layers. The prototypes represent learnt embeddings associated with actual training samples and are embedded within this latent space. The distance computation utilising Optimal Transport (OT) assesses patch-wise similarities between the input image and prototypes. OT aligns corresponding patches from the input and prototypes, consolidating patch-wise distances into a singular image-level distance. This process guarantees structural consistency within the latent space, facilitating anatomically relevant comparisons. The visualisation of patch-wise correspondences improves explainability by emphasising particular regions that affect predictions. During inference, predictions are generated through a weighted average of the labels of prototypes located within a defined distance radius in the latent space. This mechanism enables the model to utilise only applicable prototypes, enhancing interpretability and minimising the influence of unnecessary examples. This figure highlights ExPeRT’s emphasis on producing spatially grounded and interpretable predictions while achieving state-of-the-art performance in regression tasks such as brain age estimation. Integrating prototype visualisation and OT matching provides a transparent and reliable methodology for clinical applications.

**Fig. 30.**
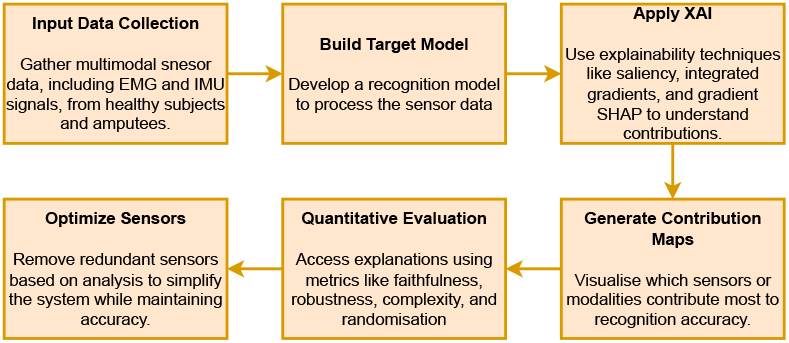
Overview of XAI-enhanced human machine interface system

**Fig. 31.**
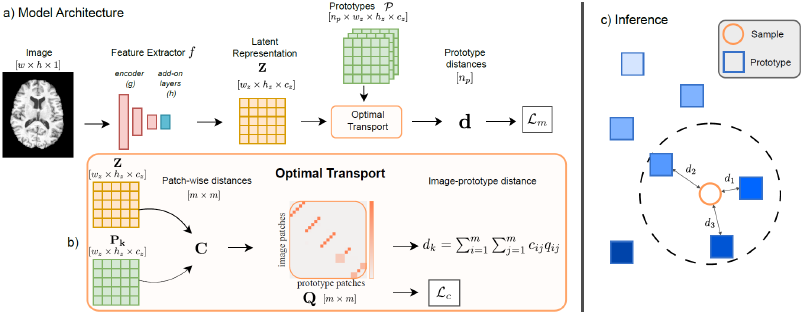
Overview of the ExPeRT model: (a) Model architecture extracts features from an image and compares them to prototypes using Optimal Transport (OT). (b) OT computes patch-wise distances to measure similarity between image and prototype. (c) During inference, predictions are based on prototypes within a defined distance radius [336]. This image is licensed under CC BY 4.0.

Focusing on Alzheimer’s disease, LAVA (2024) [337] utilised a dual-phase explanatory approach to enhance the classification. Initially, essential neurons are identified by maximum likelihood-based mutual information analysis. Subsequently, hierarchical clustering categorised patients into more specific subpopulations, elucidating nuanced illness phases. Table 8 illustrates how experts can manually reclassify or amalgamate clusters that seem clinically incongruous, thereby directing the model towards more significant subgroup delineations. Despite LAVA’s limited dataset of 100 fundus images from 61 Alzheimer’s patients, this feedback-oriented refinement method provides substantial insights for future research. Integrating supplementary patient-level information, such as genetic markers or blood test results, may enhance clustering accuracy by adding new dimensions of distinction. Furthermore, utilising transfer learning with pre-trained models from comparable domains may ease the constraints of limited datasets, allowing LAVA to tailor its explanations to more intricate clinical situations. Figure 32 depicts the LAVA framework’s detailed workflow for the classification of Alzheimer’s disease and the evaluation of its progression scale. The process begins with acquiring and preprocessing fundus images, during which retinal vasculature maps are produced utilising the AutoMorph pipeline to guarantee high-quality input data. The framework uses a VGG-16 model for binary classification, distinguishing Alzheimer’s disease (AD) from normal cognition (NC), with feature attribution maps improving interpretability. Neurone probing identifies essential neurones within intermediate network layers by applying Maximum Likelihood Estimation (MLE). The neurones that significantly influence predictions serve as the foundation for granularity explanation. Hierarchical clustering categorises patients into latent subpopulations according to activation patterns, revealing distinct disease stages within the Alzheimer’s continuum. The framework integrates neuron-level insights with hierarchical clustering, yielding interpretable and clinically relevant explanations for classification and disease progression.

**Fig. 32.**
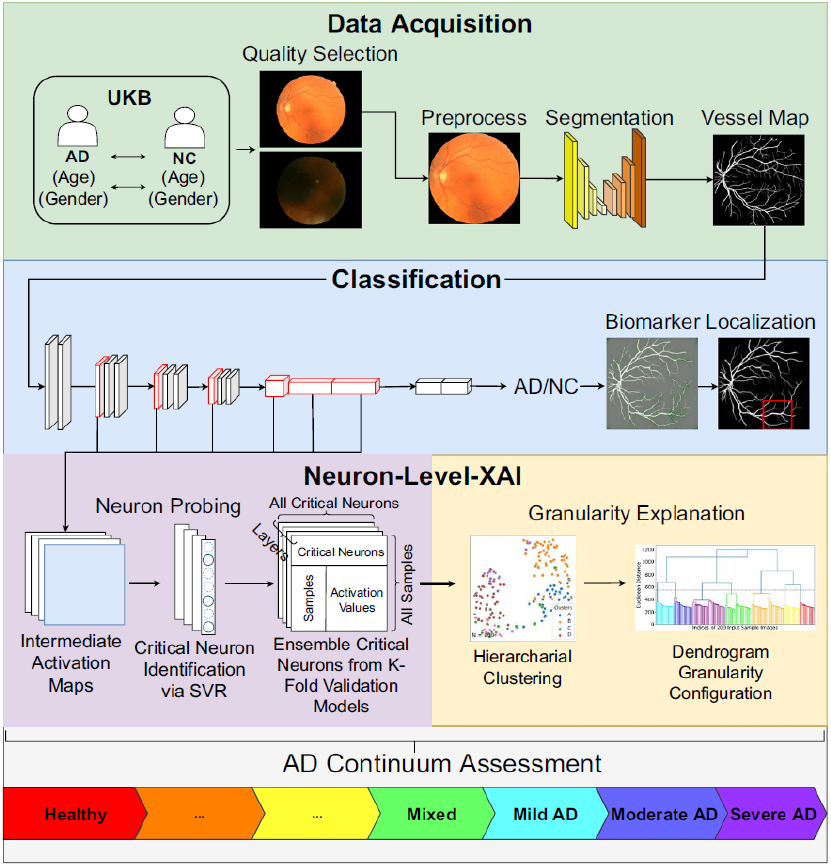
Overview of LAVA [337]. This image is licensed under CC BY 4.0.

RSA (2024) [339] employed a meta-level strategy for explanation refining by evaluating the consistency of explanations across various machine learning pipelines or preprocessing methods. The approach used Representational Dissimilarity Matrices (RDMs) to assess the consistency of explanations when small changes, such as substituting a random forest with a gradient-boosting model, are implemented. Sudden changes in RDMs can expose tenuous explanations that may lack generalisability. RSA could adopt adaptive explanation strategies to adjust attribution methods based on dataset characteristics or model design. For instance, if RSA identified significant instability in specific features, the pipeline may autonomously transition to a more resilient explanation technique, such as SHAP or Integrated Gradients, or ask a human expert to reevaluate feature engineering decisions. Moreover, employing ensemble-based methods to consolidate explanations from several models may improve consistency, particularly in critical medical applications. Figure 33 presents the framework highlighting its effectiveness in assessing the stability of machine learning explanations. The workflow initiates with model training, involving training one or more machine learning models on preprocessed datasets. Subsequently, explanation generation leads to applying posthoc explainers, including SHAP or LIME, to derive feature contribution values for a subset of test observations. The subsequent phase results in the development of Representational Dissimilarity Matrices (RDMs), in which dissimilarity metrics, such as Euclidean or correlation distance, assess variations in feature contributions among observations. The RDMs form the basis for the similarity analysis phase, wherein comparisons between pairs of RDMs demonstrate the consistency of explanations across different variations, including model architectures, preprocessing techniques, or datasets. This step identifies patterns and inconsistencies in explanatory outputs by visualising and quantifying explanation similarities. The figure highlights the framework’s capacity to conduct a meta-level evaluation of XAI methods through a systematic comparison of explanations, thereby enhancing understanding of the robustness and generalisability of machine learning pipelines. This method guarantees the maintenance of explanation consistency across various scenarios, thereby improving the reliability of decision-support systems.

**Fig. 33.**
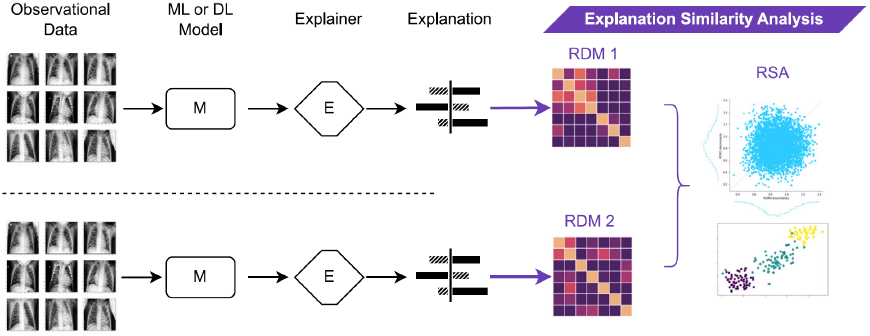
Overview of RSA [339]. This image is licensed under CC BY 4.0.

These feedback-driven frameworks illustrate that explainability is a dynamic dialogue among algorithms, domain experts, and advancing clinical insights rather than a static procedure. Through the iterative integration of expert feedback, model introspection, and error analysis, these systems enhance their explanations to reflect real-world requirements accurately. Nonetheless, obstacles persist in reconciling computational efficiency, customisation, and scalability. Implementing tactics such as on-device processing, federated learning, and adaptive explanation approaches would facilitate the development of AI systems that clarify their predictions and enhance their transparency and utility in clinical practice over time.

### D Multimodal explanation evaluation

The evaluation of model explanations remains an active area of research, particularly within the medical domain, where seeking expert opinions from radiologists and clinicians is a standard method to assess interpretability. However, this approach is time-consuming. Therefore, there have been efforts to propose objective criteria for evaluating the quality of explanations. Figure 34 presents various types of multimodal explanation evaluations.

**Fig. 34.**
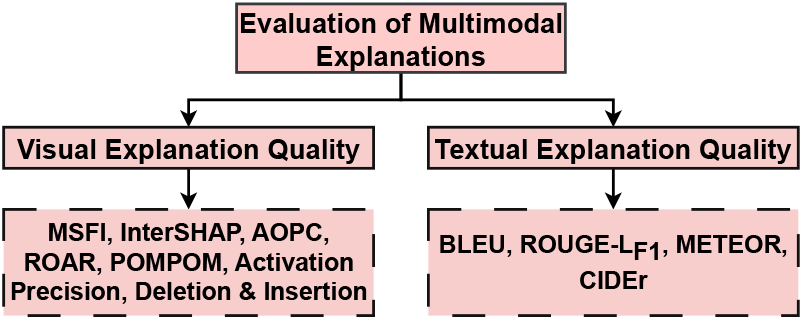
Types of Multimodal Explanation Evaluations

#### Visual Explanation Quality

For AI model predictions to enhance clinical decision support systems, they must be clearly explained to physicians. Heatmaps are commonly used to highlight relevant features in these predictions, but their effectiveness for multimodal medical images is uncertain, given the diverse clinical information they display across various channels.

The Modality-Specific Feature Importance (MSFI) [340] metric was developed to address this issue by evaluating and highlighting the significance of features specific to each modality. MSFI determines the importance or contribution of features from each modality towards the model’s predictive performance. A feature *x*_*i*_ in modality *m* is mathematically represented by its impact on the model’s output, which can be approximated using techniques like the derivative of the output for the feature or more complex techniques like Shapley values.

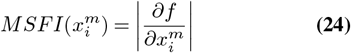

Where *f* is the model’s output function, 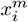 is the *i*-th feature of the *m*-th modality, and 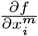 is the partial derivative of the function to that feature, representing the sensitivity of the output to changes in 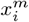.

InterSHAP [341] introduces a novel dimension to multimodal explanations by differentiating between inter-patient and intra-patient explanations. Intra-patient explanations emphasise the analysis of how various modalities influence individual predictions, particularly in assessing the relative importance of imaging data versus lab results in diagnosing a specific condition. Conversely, inter-patient explanations synthesise insights from the entire dataset, uncovering population-level patterns and identifying the most predictive modalities for specific subgroups. This technique works on multiple modalities, essential in healthcare (see Figure 35).

**Fig. 35.**
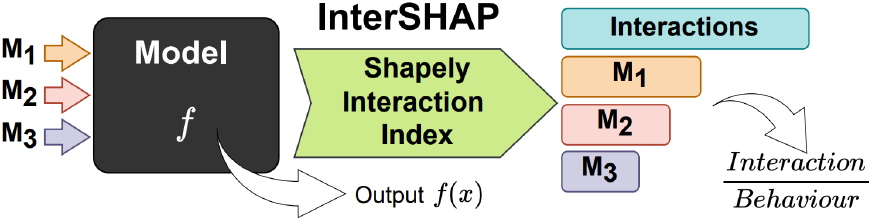
InterSHAP analyses a model’s behaviour by input perturbation from three modalities. It is defined as the ratio of interactions to overall model behaviour [341]. This image is licensed under CC-BY-NC-ND.

Samek et al. [342] introduced the Area Over the Most Relevant First (MoRF) perturbation curve (AOPC) metric. This metric uses a regional perturbation technique where regions of an input image are removed in order of their relevance to the predicted class, observing the effect on the model’s performance. The approach uses heatmaps to show that a higher AOPC value indicates greater model sensitivity to these perturbations.

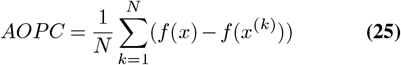

Where *f* (*x*) is the model’s output for the original image *x, f* (*x*^(*k*)^) is the model’s output after the top *k* relevant regions are removed, and *N* is the total number of perturbed regions. Inspired by [342], Petsiuk et al. [270] introduced two causal metrics, deletion and insertion, for evaluating explanations of black-box models. The deletion metric measures the drop in class likelihood as critical pixels in a saliency map are progressively removed, while the insertion metric tracks the increase in class probability as these pixels are gradually reintroduced.

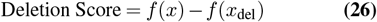

Where *f* (*x*) is the model’s confidence in the class for the original image, and *f* (*x*_del_) is the confidence after removing salient pixels.

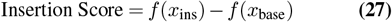

Where *f* (*x*_ins_) is the confidence after reintroducing the pixels, and *f* (*x*_base_) represents the baseline confidence without those pixels.

Hooker et al. [343] raised concerns about traditional metrics like those discussed by Petsiuk et al. [270], arguing they do not fully capture the root causes of performance declines, which might be due to artefacts from pixel replacement. They proposed the RemOve And Retrain (ROAR) technique. ROAR assesses a model’s interpretability by examining the impact on accuracy after removing key features identified by various interpretability methods and retraining the model.

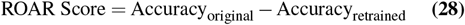

Where Accuracy_original_ is the accuracy of the original model and Accuracy_retrained_ is the accuracy after retraining the model without the identified critical features.

Rio-Torto et al. [344] developed the Percentage of Meaning Pixels Outside the Mask (POMPOM) metric to assess the effectiveness of explanations by calculating the percentage of meaningful pixels that fall outside a defined area about the total pixel count.

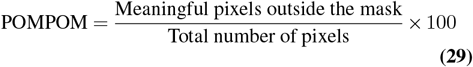

Barnett et al. [345] introduced the Activation Precision (AP) metric, which evaluates how well the information in the “relevant region” aligns with radiologist annotations in classifying mass margins.

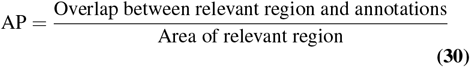

However, both metrics require manual annotations, which can be demanding and challenging for specific medical image datasets. By applying intersection or union procedures to examine consistency among highlighted Regions of Interest (ROI), we may effectively assess the interpretability effectiveness of explainable AI for multimodal medical data.

#### Textual Explanation Quality

Evaluating the quality of textual explanations involves metrics such as BLEU [346], METEOR [347], CIDEr [348], and *ROUGE* − *L*_*F* 1_ [349].

The Bi-Lingual Evaluation Understudy (BLEU) score assesses the ability of machine-translated text to match human translations. Papineni et al. [346] developed this metric to determine the level of agreement between machine-generated text and human translations. BLEU compares the n-grams of the candidate translation with reference translations and assigns a weight to these matches to calculate a comprehensive score.

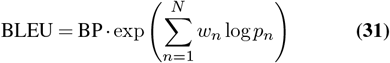

Where *p*_*n*_ is the precision of n-grams, *w*_*n*_ are weights summing to 1, typically equal for all n-grams, and BP (Brevity Penalty) addresses translation length to penalize overly short translations.

The Brevity Penalty (BP) ensures the BLEU score balances brevity with accuracy.

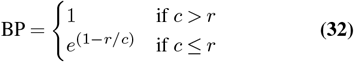

Where *c* is the length of the candidate translation and *r* is the effective reference corpus length.

The Metric for Evaluation of Translation with Explicit Ordering (METEOR) assesses translation quality, improving upon BLEU by integrating synonymy, stemming, and giving more importance to recall. Lavie et al. [347] developed METEOR to provide a more precise evaluation by balancing precision and recall.

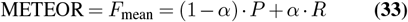

Where *P* is precision, *R* is recall, and *α* balances precision and recall. METEOR includes a penalty for poor word ordering.

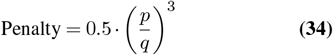

Where *p* is the number of chunks and *q* is the number of unigrams in the match.

The final METEOR score is:

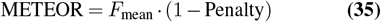

The Consensus-based Image Description Evaluation (CIDEr) score assesses the quality of textual descriptions in image captioning by measuring consensus between a candidate caption and reference captions. Vedantam et al. [348] introduced CIDEr, which uses TF-IDF weighting for each n-gram in captions.

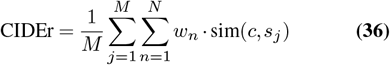

Where *c* is the candidate caption, *s*_*j*_ are the reference captions, *M* is the number of reference captions, *N* is the maximum n-gram length, *w*_*n*_ are weights for each n-gram, and sim(*c, s*_*j*_) is the cosine similarity between the candidate and reference captions, using TF-IDF weighted n-gram representations.

TF-IDF for each n-gram is calculated as:

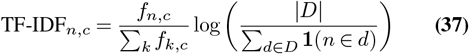

Where *f*_*n,c*_ is the frequency of n-gram *n* in capLtion *c*, |*D*| is the total number of documents (captions), and ∑_*d*∈*D*_ **1**(*n* ∈*d*) is the number of documents where the n-gram *n* appears. The *ROUGE L*_*F* 1_ score, part of the ROUGE metric suite, evaluates the quality of summaries by comparing them to reference summaries. The ROUGE-L variant focuses on the longest common subsequence (LCS) between the candidate summary and reference summaries.

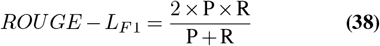

Where 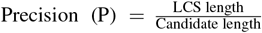 and 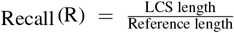.

### E Discussion

The frameworks presented in this section can be categorised according to their main objectives and approaches to data management. A primary group concentrates on specialised, domain-specific tasks to generate explanations that closely correspond to specific clinical workflows or modalities. For example, ExAID explains dermoscopic features relevant to skin-lesion classification; PathLDM produces synthetic histopathology images based on pathology reports; NeuroIGN assists neurosurgeons by integrating MRI and ultrasound data in real-time; GCN (for Alzheimer’s) discriminates relationships between biomarker and imaging features; Prosthetic Hand Control employs sensor attributions to enhance EMG hardware; and LAVA emphasises critical neurones and clusters for Alzheimer’s diagnosis using retinal fundus images. Each field, ranging from dermatology to neurosurgery, addresses a specific area while sharing a common theme: maintaining the interpretability of the output for specialised end users, including dermatologists, pathologists, and surgeons.

A second group focuses on integrating diverse data and providing more generalised, frequently automated methodologies. HAIM, EHR-KnowGen, XAI Orchestrator, and AutoXAI4Omics integrate various data streams, such as imaging, omics, lab tests, and textual notes, to develop cohesive predictive models. Visual explanations, including saliency maps and Shapley-value plots, indicate modalities significantly influencing the final decision. Frameworks such as Calibrated-XAI, CLARUS, Clinical Conversational AI Interface, VLM-Radiology, RSA, and ExPeRT focus on user interaction, feedback mechanisms, and the stability of explanations. Clinicians, data scientists, and other stakeholders are encouraged to refine system outputs by adjusting graph structures in CLARUS, rating AI confidence in Calibrated-XAI, interacting with an AI-driven radiology interface, or assessing explanation consistency through RSA. These studies demonstrate a significant transition towards XAI solutions that provide transparent predictions while also adapting to real-world clinical input and diverse data types.

Various metrics are employed across the studies to assess the effectiveness of these visual and textual explanations. Techniques such as AOPC, Deletion, Insertion, and ROAR assess the impact of modifying or removing top-ranked features on model performance, thereby identifying the critical components of the dataset. Text-based outputs are evaluated using BLEU, ROUGE, and METEOR scores to assess the clarity and accuracy of machine-generated explanations compared to human-language references. Each framework aims to connect AI with clinical reasoning while supporting its interpretability claims through measurable indicators of explanation quality.

## 5 Current Practices, Challenges, and Future Directions

Analysing complicated data in the medical industry calls for combining information from many sources, including electronic health records (EHRs), behavioural data, genetic profiles (omics), sensor readings, and medical imaging like radiology scans. The explanations these sophisticated tools generate must be clear, understandable, and aesthetically interesting to be useful for doctors. Structured approaches to link many data types with explainability techniques have been presented by recent frameworks such as CLARUS [220] and AutoXAI4Omics [324]. These models centre on tools including saliency maps emphasising the most critical areas of focus in data, such as particular regions in an image, feature-importance methods which identify the key variables or features driving artificial intelligence decisions, and counterfactual and example-based approaches letting users investigate “what-if” scenarios or review related cases to grasp the system’s reasoning better. This part summarises knowledge on present visualisation techniques, points out the challenges in generating practical and user-friendly visualisations, and provides doable suggestions for implementation. It also describes potential avenues of study meant to improve these techniques so they might support clinical decision-making more.

## A Current Visualisation Practices

### A.1 Commonly Applied Visualisation Methods

#### Backpropagation-based Saliency Maps

Medical imaging extensively uses backpropagation-based saliency maps to describe which portions of an image a model focuses on during prediction. These methods, for instance, can show suspicious areas on MRI or CT images, helping doctors find anomalies. Popular techniques such as Grad-CAM and Grad-CAM++ superimpose heatmaps onto the original image, displaying the spatial areas most likely influencing the model’s choice. Layer-wise Relevance Propagation (LRP) offers pixel-level relevance ratings, providing a more comprehensive picture of the model’s emphasis. Recent studies show that these methods are especially successful for convolutional neural networks (CNNs) since they can visually localise areas of interest, including tumour tissues or lesion boundaries. For example, although LRP has helped define lesion borders in transfer learning applications [249], Grad-CAM has been utilised to identify tumour areas [337].

#### Feature Importance Techniques

Making meaning of organised data like electronic health records (EHRs), genetic profiles, and time-series sensor data depends critically on feature significance methods. These techniques provide greater understanding and confidence in AI systems applied in healthcare by helping to clarify the contribution of particular features to the model decision [350]. LIME is a generally applied technique approximating complicated models with simpler local surrogate models. LIME is very helpful for low-dimensional reduction by feature identification for every prediction [337, 350]. Using Shapley values from cooperative game theory, SHAP is another widely used method that computes feature contributions at both global and local levels. Whether for a single patient or over a whole patient cohort, SHAP offers insights into which factors, e.g., age, blood pressure, and glucose levels influence a model’s judgements most. These instruments increase interpretability and help doctors concentrate on the most pertinent factors, increasing the actionability and dependability of artificial intelligence systems in clinical environments.

#### Counterfactual and Example-Based Explanations

Two powerful methods for improving the interpretability of artificial intelligence models in healthcare are counterfactual and example-based explanations. Particularly helpful for personalising and decision-making, they offer insights by demonstrating how modifications to specific input features could affect the model predictions. Emphasising “what-if” situations, counterfactual explanations show how changing particular variables such as a test result or genetic marker may produce a different conclusion. For example, tools like DiCE [225] and ExPeRT [336] let people investigate practical insights including suggested lifestyle modifications that might lower disease risk. These clear-cut justifications help to close the distance between sophisticated artificial intelligence systems and pragmatic healthcare interventions.

To make forecasts more relevant, example-based methods employ representative examples, including ProtoPNet [253] and MProtoNet [319]. These approaches enable doctors to grasp better how a model classifies inputs by showing typical cases inside every class. For example, they might draw attention to a typical tumour image to explain why a particular scan came out malignant. Recent developments, including the CLARUS framework [220], have integrated interactive dashboards with counterfactual explanations. These systems let doctors examine explanations in real-time, helping them hone or better grasp the AI’s thinking mechanism. By including counterfactuals in user-friendly tools, AI systems are guaranteed to be accurate but also open and practical in clinical settings.

### A.2 Visualisation Techniques for Bridging Modalities

#### Integrated Attention Maps

Attention mechanisms are fundamental in multimodal systems, facilitating the integration of various data types, including imaging and electronic health records (EHRs) while improving interpretability. These mechanisms dynamically assess the contribution of each modality, ensuring that both the model and the user comprehend the relative significance of different data sources in prediction-making. An integrated attention map may indicate that particular genetic markers and lesion locations identified in MRI scans significantly affect diagnostic outcomes. Techniques like Gradient-weighted Class Activation Mapping (Grad-CAM) and attention-weighted embeddings enhance these capabilities, facilitating detailed interpretability.

Recent frameworks such as NeuroIGN [331] combine MRI and intraoperative ultrasound (iUS) data, providing real-time surgical visualisations that are dynamically updated with attention-based certainty maps to assist decision-making. EHR-KnowGen [332] utilises attention-based mechanisms to align structured and unstructured EHR data, highlighting cross-modality interactions that clarify essential clinical variables influencing outcomes. These frameworks illustrate the role of attention maps in integrating various data sources, offering clinicians nuanced insights into contributions specific to each modality. This increased transparency fosters trust in AI-driven decisions and enables clinicians to prioritise the most significant data sources for patient-specific situations.

#### Hybrid Fusion Visualisation

Recent advancements in models such as HAIM [329], NeuroIGN [331], PathLDM [330], and LAVA [337] have markedly enhanced the integration of explainability within multimodal medical data workflows, equipping clinicians with practical tools for interpreting intricate predictions. HAIM employs modular fusion embeddings to combine modalities, including imaging, tabular, and textual data, enhancing tasks like 48-hour mortality prediction. It emphasises high-impact features and ensures explainability through Shapley values. NeuroIGN integrates preoperative MRI with real-time intraoperative ultrasound (iUS) data through hybrid fusion techniques, aligning multimodal imaging to provide interpretable real-time segmentation during neurosurgical procedures. This integration assists surgeons in making informed decisions in critical clinical environments. PathLDM integrates pathology reports and histopathology whole slide images (WSIs) via text-conditioned latent diffusion, successfully uniting global textual information with localised tumour imaging characteristics. This hybrid method improves synthetic image generation and assists clinicians in situations with scarce annotated data. LAVA utilises a hybrid framework to integrate retinal fundus imaging and cognitive assessments, clarifying Alzheimer’s progression through hierarchical clustering of multimodal data. By incorporating various modalities into unified visualisations, these frameworks enable clinicians to obtain actionable insights, thereby reducing the disparity between artificial intelligence-driven discoveries and their practical clinical applications.

#### Temporal Visualisations

Artificial intelligence models must address specific challenges to interpret wearable sensor data, and time-series signals such as ECG or EEG while capitalising on significant clinical insight opportunities. Visual aids, including heatmaps and trend graphs, effectively emphasise critical intervals or patterns relevant to diagnosis. Heatmaps can identify periods of increased activity or anomalies in EEG data, whereas trend graphs can depict long-term changes, emphasising irregularities associated with clinical outcomes. These visualisations allow clinicians to concentrate on the most relevant data segments, enhancing their capacity to correlate model outputs with clinical observations.

Frameworks like HAIM [329] employ temporal saliency maps to illustrate the contribution of specific time intervals or sequences to mortality prediction, thereby assisting clinicians in identifying critical trends within time-series data. Similarly, EHR-KnowGen [332] integrates structured EHR data with time-series features, using attention-weighted embeddings to emphasise impactful temporal patterns dynamically. Integrating multimodal fusion methodologies demonstrates how temporal relationships among data points can improve interpretability. Advanced layered designs improve the comprehension of multimodal interactions, including integrating sensor readings with physiological signals. Frameworks such as NeuroIGN [331] synchronise real-time imaging with temporal variations in patient vitals, providing interpretable feedback during neurosurgical procedures. These visualisations maintain chronological and contextual significance, enabling clinicians to make informed decisions based on communicated temporal patterns.

#### Dimensionality Reduction

High-dimensional data analysis, including coupled omics and imaging features, is greatly enhanced by dimensionality reduction techniques such as t-SNE, UMAP, and PCA. By projecting intricate representations into interpretable two-dimensional or three-dimensional spaces, these methods reveal clusters, patterns, and relationships that may remain obscured in complex data. t-SNE is particularly effective in preserving local relationships, which enhances its ability to identify small-scale clusters within data. UMAP balances the preservation of regional and global structures, frequently resulting in more precise and cohesive visualisations for multimodal datasets. PCA reduces the complexity of high-dimensional data by identifying principal components that account for the most significant variance, proving particularly useful for datasets characterised by linear relationships.

When applied carefully, these techniques preserve critical interconnections across modalities and decrease complexity, facilitating intuitive and efficient data exploration. Frameworks such as HAIM [329] employ dimensionality reduction techniques to handle multimodal inputs, effectively prioritising the most relevant features from imaging, time series, and tabular data for enhanced interpretability. PathLDM [330] integrates feature selection and dimensionality reduction to analyse high-dimensional pathology data, creating interpretable synthetic histopathology images. The EHR-KnowGen [332] framework illustrates the potential of dimensionality reduction to improve the integration of structured and unstructured EHR data by refining feature embeddings, allowing researchers to concentrate on clinically relevant patterns. These frameworks demonstrate the capability of advanced AI systems to manage diverse, highdimensional datasets while maintaining interpretability and usability for medical practitioners. Integrating dimensionality reduction and intelligent feature selection methodologies simplifies the exploration of large-scale genomic and imaging datasets while preserving their clinical relevance. This method improves multimodal AI systems’ interpretability and practical use in healthcare.

### A.3 Interactive Visualisation Tools

#### Dashboard-Based Exploration

Interactive tools to examine and comprehend AI model behaviour are fundamental to modern XAI-enabled platforms, including CLARUS [220] and EHR-KnowGen [332]. These systems offer clinicians and researchers dynamic dashboards for real-time visualisation of model predictions, facilitating the editing of parameters, modification of input data, and adjustment of settings. This interactivity creates a setting where clinicians can actively investigate AI models, assessing the influence of specific factors or changes on the outcomes. EHR-KnowGen integrates structured and unstructured EHR data into unified explanations, enabling users to dynamically explore the relative importance of various data modalities.

These tools are especially useful in complex multimodal contexts, where comprehending the interactions among various data types, such as imaging, textual reports, and time-series data can pose significant challenges. Dashboards in PathLDM [330] and HAIM [329] offer layered visualisations that integrate global summaries with detailed insights. PathLDM employs user-friendly interfaces to aid in synthesising pathology reports and histopathology images. In contrast, HAIM highlights the roles of individual modalities, allowing clinicians to customise analyses for particular patient cases. Interactive platforms improve trust and comprehension by connecting AI systems with domain experts. Visualising the real-time effects of parameter changes enables clinicians to refine models with AI engineers collaboratively. This capability is essential for modifying AI solutions to address the distinct requirements of particular clinical settings or patient groups. Systems such as CLARUS, EHR-KnowGen, and HAIM enhance the usability of AI in healthcare by facilitating dynamic exploration, promoting deeper insights, and aligning AI outputs with clinical priorities. These platforms illustrate the capacity of dashboard-based exploration to convert AI into a collaborative, transparent, and actionable resource for medical decision-making.

#### What-If Tools

Clinicians and researchers are transforming their engagement with AI models by using dynamic tools like Google’s What-If [320], which facilitates real-time scenario analysis. Users can modify patient-specific variables such as BMI, laboratory values, or genomic risk factors and analyse the resulting prediction changes. This interactive method enhances model interpretability, providing clear insights into the contribution of features to clinical outcomes.

In addition to these fundamental tools, advanced frameworks like SCouT [351] and GCNs [334] enhance “what-if” capabilities through spatiotemporal synthetic data generation and adjacency-based visualisations, respectively. SCouT addresses the issue of sparse datasets by modelling realistic variations, facilitating the exploration of patient trajectories in data-constrained scenarios. GCNs enhance this by visualising patient relationships through graph networks, highlighting individual and group-level features essential for differentiating clinical subgroups, including mild cognitive impairment and Alzheimer’s disease progression. The incorporation of counterfactual generation, as illustrated in ProtoPNet [253], identifies minimal alterations to input parameters that would modify a diagnosis, offering practical insights. LAVA integrates imaging and cognitive assessment data for prototype-based reasoning, enabling users to evaluate hypotheses regarding the influence of patient features on diagnostic categories.

These tools demonstrate the integration of “what-if” capabilities in multimodal AI systems, enhancing clinician trust and aiding clinical decision-making. They facilitate real-time interaction, scenario testing, and comprehensive feature exploration, connecting abstract AI outputs with practical medical insights and supporting AI integration in critical healthcare applications.

## B Challenges in Visualisation

Although the above mentioned techniques show considerable potential, using them in practical clinical environments involves numerous major difficulties, particularly considering different data sources.

### B.1 Balancing Complexity and Interpretability

In multimodal environments, where many data sources such as MRI scans, EHR data, and genetic markers must be combined, the difficulty of balancing complexity and interpretability is more evident. On one side, very complicated, multi-layered explanations might confuse doctors and clinicians, making them useless for practical application [352, 353]. Conversely, too simplistic models, such as single saliency maps, risk obscuring critical interactions between various modalities.

#### High Dimensionality

Computational efficiency and interpretability are challenged by datasets including hundreds of features such as genetic variations, imaging pixels, and time-series sensor data. Not only are these high-dimensional datasets resource-intensive to handle, but they also challenge meaningful and accessible presentation for clinicians [350]. Focuses on strong feature selection to rank the most significant variables, lowering complexity and maintaining important information from PSGIF [350]. Integrating genetic and imaging data also helps multimodal approaches. LAVA [337] investigates intermediate layers of convolutional neural networks (CNNs) to understand model processing, therefore facilitating the improved interpretation of important characteristics derived from high-dimensional input. Although these technologies offer great value, their accessibility in clinical environments is limited since they are usually targeted at technical users. User-friendly pipelines are essential to enable the management and interpretation of high-dimensional data. Methods including t-SNE, UMAP, and PCA provide promising techniques for dimensionality reduction. Still, they need adaption to protect clinical relevance, verifying that dimensionality reduction does not cause loss of essential links between imaging characteristics and genetic markers. Additionally, simplified interfaces offer clinicians easily comprehensible outputs, including 2D or 3D visualisations with overlays or annotations. Incorporating clever preprocessing and parameter optimisation helps lower the hand corrections requirement. Frameworks such as those suggested in LAVA and PSGIF should be developed to encompass these capabilities, producing technically strong tools that are both technically strong and understandable to non-technical users [326]. Translating high-dimensional insights into practical medical decisions will be simpler by increasing dimensionality reduction techniques and smoothly including them in clinical processes. Future studies should concentrate on closing the technical complexity-usability gap to guarantee that these technologies are as helpful as they are strong.

#### Accuracy–Interpretability Trade-off

Deep neural networks (DNNs), among other models with high prediction accuracy, are sometimes considered “black boxes” because of their complexity and lack of openness. On the other hand, simpler models such as linear regression or tiny decision trees are easier to understand but find difficulty managing the intricacy of multimodal data and complicated feature relationships [325, 336]. Implementing Explainable AI (XAI) in clinical settings presents great difficulty in managing this trade-off. DNNs shine in capturing complex interactions across multimodal data but typically lack simple mechanisms for physicians to grasp their thinking, posing challenges in balancing accuracy and interpretability. Simpler models may miss significant insights or patterns in high-dimensional datasets, even if they are interpretable. Clinicians need accurate forecasts and clear, understandable explanations to help decision-making and establish confidence.

Advanced and hybrid XAI techniques aiming to preserve the predictive capacity of complicated models while improving their interpretability have evolved to handle these challenges. ExPeRT [336] explains predictions utilising exemplary examples inside every class. Linking the outputs of a model to relevant, real-world instances closes the distance between clinical intuition and abstract model thinking. This method is very successful for multimodal systems, where examples might span several data, such as integrating an imaging scan with crucial EHR features. MACO [325] presents limitations during model training to generate more natural and human-friendly visual explanations. This guarantees the interpretability of outputs without sacrificing the model’s prediction accuracy. MACO can be used in situations such as saliency maps to ensure that highlighted areas or characteristics match human expectations, increasing its practicality for doctors.

Maintaining a clinically acceptable trade-off calls for ongoing innovation in hybrid XAI methods combining high-accuracy models with interpretable overlays, including saliency maps, feature attributions, or counterfactual scenarios. Designing interfaces that show explanations in a clinician-friendly way will help you to emphasise usability. Including strong validation will help to guarantee that interpretability characteristics complement clinical knowledge and expectations. The healthcare sector may fully utilise high-performance AI systems using ExPeRT and MACO, ensuring their forecasts stay clear and actionable in practical clinical environments.

The balance between model complexity and interpretability is essential in high-stakes medical applications, as clinicians need dependable explanations while maintaining predictive accuracy. The eXirt method [171], although computationally demanding due to its perturbation-based analysis, offers organised and probabilistic insights into feature relevance. In contrast to conventional post-hoc explainability methods, eXirt quantifies interpretability using metrics of *discrimination, difficulty*, and *guessing*, thereby providing a robust framework for feature evaluation. The methodological rigour of eXirt makes it particularly significant in critical domains such as oncology, neurology, and cardiology, where AI-driven diagnostics necessitate transparency. In tumour classification models, eXirt can identify MRI-based imaging biomarkers with the most significant discrimination power, thereby enhancing clinicians’ comprehension of the model’s rationale for malignancy predictions. In cardiovascular risk assessments, difficulty scores can identify ambiguous cases that require clinical judgment. eXirt balances computational cost with interpretability, allowing AI-driven medical tools to be both explainable and clinically actionable, thereby ensuring that the need for transparent, trustworthy explanations surpasses the computational expense.

### B.2 Standardising Visualisation Practices

Unlike single-modality data, multimodal datasets mix several sources, each with unique qualities and visualisation needs. This variability seriously challenges the development of consistent methods for visualising and assessing Explainable AI (XAI) outputs [339]. Often, lacking set norms results in inconsistent or inferior findings across studies, impeding field advancement.

#### Heterogeneous Data Types

One of the key challenges is heterogeneous data types. Multimodal data combines structural, unstructured, geographical, and temporal components and calls for different visualisation methods for every modality. In particular, Imaging data, for instance, gains from saliency maps or attention overlays; EHR data is better represented with feature attribution scores or counterfactuals. Combining these still presents difficulty. Unlike single-modality explainability, whereby standard metrics (e.g., authenticity, sparsity) are more defined, multimodal XAI lacks uniform criteria to evaluate whether explanations are trustworthy, interpretable, and therapeutically significant. Visualising the total contributions of all modalities in a clear, interpretable manner often results in oversimplification, maybe hiding crucial information. Comparing results across research or including findings in meta-analyses becomes challenging without uniform approaches to evaluate the quality of XAI outputs.

#### Domain-Specific Constraints

Every modality in multimodal datasets has unique qualities that make developing consistent evaluation rules more difficult. Radiology, for example, typically uses localised saliency maps or attention overlays to emphasise areas of importance, such as MRI scan tumour margins. Conversely, genomic data calls for explanations that address high-dimensional and feature-specific interactions, such as the influence of gene variations. EHR data gains from feature attribution scores linked to particular patient criteria and textual rationales. These variations make it challenging to create a one-size-fits-all assessment since a single “explanation quality” score usually misses the subtleties of localised visual explanations against textual reasons. Dealing with these domain-specific limitations calls for tailored evaluation plans that fit the modality, maintaining coherence in multimodal environments.

#### Lack of Unified Protocols

A significant obstacle to XAI’s progress in multimodal settings is the lack of standardised benchmarks and metrics to evaluate dependability and interpretability. Designed for particular domains, current techniques such as AOPC and ROAR are suitable with saliency maps, while BLEU and CIDEr are useful as text-based metrics. These measures, however, cannot meet the particular needs of multimodal medical data, so explanations must coincide across several data kinds, including imaging, genomics, and EHRs. Verifying the dependability and interpretability of XAI outputs is a subjective process without a universal framework, impeding outcomes’ comparability between investigations.

#### Framework Development

The potential solutions include framework development. Provide consistent models for multimodal XAI visualisation addressing several data formats. These models should include models for combining across modalities saliency maps, feature importance, and counter-factual examples. Establish benchmarks for assessing multi-modal XAI outcomes by combining criteria, including evaluating whether explanations fit domain-specific patterns. Ensure proper representation of interactions between modalities is cross-modal consistency, and use physician comments and human-centred metrics to test interpretability and usability. Create tools that let doctors interactively investigate XAI outputs, cycling between modalities and degrees of explanation granularity to grasp the system’s reasoning better. Encourage multidisciplinary cooperation among physicians, domain experts, and artificial intelligence researchers to create and implement homogenous standards for visualisation and evaluation. Advancement of the area depends on requirements for multimodal XAI. Uniform methods would raise the consistency and quality of research and help AI systems in clinical practice to be more interpretable, actionable, and trustworthy.

### B.3 Bridging Modalities in Visualisations

A significant difficulty in Explainable AI (XAI) for healthcare is developing unified, coherent visualisations that include many data sources. Effective multimodal visualisations must be interpretable for topic experts yet capture cross-modal interactions.

#### Late Fusion Complexity

Late fusion techniques aggregate outputs from independently trained models for every modality, often find providing aligned and significant feature attributions challenging. For instance, disjointed or poorly synchronised saliency maps from imaging and tabular data can complicate understanding how each modality’s factors contribute to the general forecast. These approaches run the danger of excluding cross-modal interactions crucial for proper diagnosis, such as the interaction between lesion locations in an MRI and genetic markers discovered in omics data [354]. Dealing with this difficulty requires designing early or hybrid fusion pipelines that combine modalities during model training, enabling interaction-aware attributions. Using post-hoc tools allowed visualising interactions across several modalities instead of considering every input as autonomous.

#### Knowledge-Domain Gap

The variation of knowledge among doctors adds even more difficulty to understanding multimodal visualisations. Imaging experts might not have the background to decipher genomic patterns or gene-expression saliency maps. Geneticists could thus find overlays less relevant or even deceptive on MRI scans without background [355]. This mismatch in domain knowledge reduces the utility of homogeneous visualisations. Platforms such as CLARUS [220] let doctors fine-tune and adjust AI explanations to their particular fields of expertise through interactive feedback. Customised visualisations should show explanations in domain-relevant forms, e.g., thorough genetic attributions for molecular scientists and image overlays for radiologists.

## C Guidance for Effective Visualisation Practices

Establishing efficient visualisation techniques for multimodal medical data calls for resolving the complexity of aggregating several data sources and guaranteeing that the resultant tools remain clinically helpful and practical. This requires careful design ideas to strike a mix of thorough coverage, usability, and clarity.

### C.1 Modular Visualisations

One good tactic is using modular visualisations that divide the presentation of every data modality. Imaging data, for example, can be shown by heatmaps; bar charts can show EHR characteristics. These visual panels let doctors concentrate on each modality separately, lowering the cognitive strain. Turning between these panels helps users grasp individual contributions clearly before synthesising ideas from the aggregated data [67].

### C.2 Layered Visualisations

Layering visualisations is another helpful strategy for offering high-level summaries and thorough explanations. Starting with summaries, like confidence scores for every modality, for instance, enables doctors to evaluate the general performance of the model quickly. They can then probe into particular elements, such as key time intervals in sensor data or localised saliency maps for MRI areas. This hierarchical system guarantees that the visualisation is flexible enough for quick evaluations and more thorough investigation by experts [328].

### C.3 Fusion Visualisation

Though it brings unique difficulties, combining modalities in one visualisation is essential for multimodal artificial intelligence. Clearly expressed should be the approach employed to mix modalities. Whereas hybrid fusion could reveal the extracted features of each modality before they are integrated, early fusion could show combined feature embeddings. Late fusion can show the different forecasts for every modality and their aggregations. Clear vision of these fusion processes helps clinical interpretation [149, 153]. It also makes the relationships between modalities more visible.

### C.4 Domain-Specific Visual Aids

Development of trust and usability depends on domain-specific visual aids. Visualisations should fit the procedures of particular medical specialities. Heat maps should highlight suspicious areas in radiography, while text annotations help aggregate patient statistics. Genomics visualisations might link gene-expression data onto pathways or biomarkers the domain specialists already know. This congruence with current knowledge systems facilitates physicians’ understanding and confidence in the AI-generated outputs [220, 355].

### C.5 User-Centric Design

Another essential component of good visualisation is prioritising user-centric design. Interactive dashboards give doctors more control and help them better comprehend by letting them modify model parameters or thresholds. Feedback loops, including clinicians, also assist models and their visualisations to better align with clinical priorities and remain relevant to changing processes [324, 354].

### C.6 Centralised Multimodal Dashboards

Eventually, centralised multimodal dashboards showing the value of every modality side by side can simplify cross-modal study. These dashboards help doctors link imaging results with lab results or genomic data by combining visual components, including saliency maps, bar charts, and idea highlights. This combined perspective helps improve decisionmaking and more accurate diagnostics [332].

Combining modularity, layering, fusion clarity, domain alignment, interactivity, and centralised dashboards will help to create powerful and readily available visualisations driven by artificial intelligence in real-world clinical environments.

## D Future Research Directions

Though explainable artificial intelligence (XAI) visualisation for healthcare uses has come a long way, much space for improvement and innovation still exists. These developments can enable XAI to be more deeply integrated into healthcare processes, producing broader and more significant effects.

### D.1 Evaluate Visualisation Effectiveness

Evaluating how successfully XAI visualisations satisfy the needs of its end users helps one to maximise the value of these tools. Methodical user studies help to shed light on how various explanation strategies are seen and applied in reality. Eye-tracking and usability assessment methods can expose minute interactions between users and visual aids. Furthermore, providing quantifiable standards to evaluate the advantages of new visualisation approaches [326] are objective measures, including diagnostic accuracy, speed to diagnosis, and user happiness. Such assessments point to areas needing work and open the path for user-centric design ideas in XAI for healthcare.

### D.2 Automate Visualisation Refinement

Dynamic refinement is an interesting way to improve the usability of visualisations depending on user feedback. In response to real-time physician inputs, interactive refinement systems could dynamically change saliency levels or reorganise feature lists. These systems might learn over time to provide ideal degrees of information catered to particular modalities and user preferences. Through emphasising the most important findings, such automated methods could enable faster, more precise decision-making [321].

### D.3 Advance Interactive Tools

Interactive visualisation tools can transform how doctors interact with varied multimodal data. Improved “what-if” analytic tools would let users concurrently change several kinds of data, including electronic health records (EHRs) and imaging markers, and view the recalibrated forecasts in real time. This interaction may also incorporate counterfactual analysis, showing how particular changes such as choosing a different imaging slice or fixing a laboratory value, might greatly influence model outputs [235]. These developments might help doctors better grasp model behaviour and have more confidence in AI-generated insights.

### D.4 Data Scarcity and Synthetic Data

Particularly with rare diseases or under-represented populations, healthcare databases frequently suffer from problems, including data imbalance and shortage. Transformer-based models and generative adversarial networks (GANs) are starting to show remarkable ability for creating synthetic datasets to close these gaps. Pipelines such as SCouT [351] show, for example, the capacity to replicate varied patient populations. Particularly for neglected diseases and sensitive demographics [350], future research should seek to build generative approaches capable of precisely capturing the complexity of multimodal data while maintaining ethical and practical considerations.

### D.5 Explainability for Reinforcement Learning

Applications in healthcare, such as patient management and treatment planning, are finding favour for reinforcement learning (RL). Its opaque decision-making policies, however, provide significant adoption difficulties. Visualising RL components such as reward trajectories, policy decisions, and hypothetical alternatives helps close the trust gap. For instance, clarifying why a given treatment (e.g., chemotherapy) is recommended over another (e.g., surgery) at a given level of treatment will help Adapting MACO [325] to RL scenarios may help to preserve interpretability in sequential decisionmaking further, so strengthening trust and utility in clinical contexts.

### D.6 Integration of Human Factors

Future research must prioritise refining human-centred XAI to ensure alignment with clinical workflows and patient requirements. Existing approaches enhance interpretability; however, adaptive explanation systems that adjust complexity according to user expertise such as simplified patient narratives and uncertainty aware insights for clinicians may improve usability [355].

Future XAI tools should integrate interactive feedback mechanisms, enabling domain experts to iteratively refine explanation strategies, building on frameworks such as CLARUS[220]. The eXirt framework [171] may be enhanced to deliver real-time adaptive IRT-based visualisations, customising explanations according to diagnostic uncertainty and clinician preferences. Subsequent research should examine the influence of explainability on decision confidence in high-stakes contexts, ensuring that AI-generated recommendations are both transparent and actionable.

## 6 Conclusion

This survey shows that Explainable AI (XAI) in healthcare has advanced from generic model introspection to providing customised, multimodal explanations appropriate for complex medical data. Research integrating imaging, laboratory results, electronic health records (EHRs), and omics data demonstrates that multimodal strategies improve diagnostic efficiency. However, they present difficulties comprehending how many data sources collectively influence predictions. Research advancements utilising LIME, SHAP, and other perturbation-based methodologies have progressed significantly, coupled with concept-based approaches and prototype-driven explanations that facilitate clinicians’ intuitive interpretation of model outcomes. Moreover, counterfactual explanations have become an effective tool for medical practitioners to investigate “what if” scenarios, offering an enhanced understanding of model-based suggestions. Despite these positive advances, significant obstacles exist. A significant drawback is the lack of defined metrics for assessing the quality and reliability of explanations generated by XAI. Although image-based interpretability is supported by defined evaluation criteria in medical imaging, textual and numerical data do not possess globally recognised benchmarks. Integrating XAI into practical clinical workflows such as real-time monitoring, decision support systems, and extensive genomics platforms is a developing effort. Effective visual representation is essential, as medical professionals need clear, simple, and actionable information that corresponds with their cognitive workflows. Cooperative initiatives between AI researchers and physicians are crucial for developing interactive, multi-faceted explanations that merge transparency with usability.

Multiple intriguing pathways require additional investigation to solidify XAI’s position in patient-centred treatment. Federated learning methodologies may improve explainability while safeguarding patient privacy, promoting wider acceptance in decentralised healthcare settings. Reinforcement learning models that use frameworks that are easy to understand can adapt to patient answers better than blackbox AI. Furthermore, generative models may be utilised to produce synthetic variations of under-represented clinical cases, fixing dataset imbalances and explaining regions of model uncertainty. Interdisciplinary collaboration is essential for advancement, bringing together data scientists, machine learning specialists, and healthcare practitioners to create transparent and therapeutically applicable solutions. If XAI can resolve algorithmic predictions with human thinking, its revolutionary impact in healthcare will be unmatched. Explainability-driven AI can improve diagnostic precision, enable personalised treatment approaches, and cultivate trust between physicians and patients, promoting a more transparent and ethically accountable future in medical AI. This systematic review of XAI data visualisation in multimodal medical contexts enhances understanding of Responsible AI, facilitating more transparent healthcare solutions.

## Statements & Declarations

### Ethical Approval and Consent to participate

Not applicable.

### Human and Animal Ethics

Not applicable.

### Consent for publication

Not applicable.

### Availability of supporting data

Data sharing does not apply to this article as no datasets were generated or analysed during the current study.

### Data Availability

Data sharing does not apply to this article as no datasets were generated or analysed during the current study.

### Code availability

My manuscript has no associate code.

## Footnotes

1 https://www.darpa.mil/

2 https://www.engineeringvillage.com/home.url

3 https://www.scopus.com/

4 https://apps.webofknowledge.com/

5 https://pubmed.ncbi.nlm.nih.gov/

6 https://scholar.google.com/

8 https://pair-code.github.io/what-if-tool/

9 https://github.com/ClearExplanationsAI/CLEARImage

10 https://github.com/IntelAI/intel-xai-tools

11 https://cloud.google.com/vertex-ai/docs/explainable-ai

12 https://pair-code.github.io/what-if-tool/

13 https://github.com/interpretml/interpret

14 https://github.com/adib0073/EXMOS/

15 https://rshiny.gwdg.de/apps/clarus/

16 https://github.com/ai-med/KeepTheFaith

17 https://github.com/aywi/mprotonet

18 https://github.com/fzi-forschungszentrum-informatik/TSInterpret

19 https://github.com/josephenguehard/timeinterpret

20 https://github.com/IBM/AutoXAI4Omics

21 https://github.com/deel-ai/xplique

22 https://github.com/tensorflow/lucid

